# Structure-function analysis of a calcium-independent metacaspase reveals a novel proteolytic pathway for lateral root emergence

**DOI:** 10.1101/2023.01.15.523950

**Authors:** Simon Stael, Igor Sabljić, Dominique Audenaert, Thilde Andersson, Liana Tsiatsiani, Robert P. Kumpf, Andreu Vidal-Albalat, Cecilia Lindgren, Dominique Vercammen, Silke Jacques, Long Nguyen, Maria Njo, Álvaro D. Fernández-Fernández, Tine Beunens, Evy Timmerman, Kris Gevaert, Jerry Ståhlberg, Peter V. Bozhkov, Anna Linusson, Tom Beeckman, Frank Van Breusegem

## Abstract

Metacaspases are part of an evolutionarily broad family of multifunctional cysteine proteases, involved in disease and normal development. Despite the extensive study of metacaspases in the two decades since their discovery, the structure-function relationship of metacaspases remains poorly understood. Furthermore, previous studies on their function have been thwarted by the redundancy in gene copy number and potential phenotypic suppression of genetic mutations, especially in plants. Here, we have solved the X-ray crystal structure of an *Arabidopsis thaliana* type II metacaspase (AtMCA-IIf) that belongs to a particular sub-group that does not require calcium ions for activation. Compared to crystal structures of other metacaspases and caspases, the AtMCA-IIf active site is structurally similar and poses a conundrum for the catalytic mechanism of the cysteine-histidine dyad. To study metacaspase activity in plants, we developed an *in vitro* chemical screen to identify small molecule metacaspase inhibitors. Several hits with a minimal thioxodihydropyrimidine-dione (TDP) structure were identified, some being specific inhibitors of AtMCA-IIf. We provide a mechanistic basis for inhibition by the TDP-containing compounds through molecular docking onto the AtMCA-IIf crystal structure. Finally, a TDP-containing compound (TDP6) was effective at inhibiting lateral root emergence *in vivo*, likely through the inhibition of metacaspases that are specifically expressed in the endodermal cells overlaying developing lateral root primordia. In the future, the small compound inhibitors and crystal structure of AtMCA-IIf can be used to study metacaspases in various other species, such as important human pathogens including those causing neglected diseases.

## Introduction

Metacaspases are part of the C14 family of cysteine dependent proteases together with caspases and paracaspases (Rawlings *et al*, 2018). Metacaspases are conserved throughout plants, fungi, protists and bacteria, whereas caspases and paracaspases are present in mammals. The interest in metacaspases was initially sparked by their sequence similarities, entailing a common hemoglobinase structural fold and a conserved histidine/cysteine dyad in the catalytic domain, with the well-studied mammalian caspases and hence a suspected functional involvement in programmed cell death (PCD) outside metazoa (Uren *et al*, 2000; Van Doorn *et al*, 2011). Two decades later it becomes clear that metacaspases are a diversified clade of multifunctional proteins that diverge from caspases in protein structure, substrate preference and functionalities (Vercammen *et al*, 2004; Tsiatsiani *et al*, 2011; Lam & Zhang, 2012; Minina *et al*, 2017, 2020; Klemenčič & Funk, 2019). Unlike caspases that cleave their substrate proteins after aspartate, metacaspases cleave exclusively after the basic amino acids arginine and lysine. Metacaspases are classified according to their domain organization: Type-I metacaspases have an N-terminal pro-domain that is absent in type-II metacaspases, and the latter have a longer linker connecting the p20 and p10 regions. Type III metacaspases were more recently identified in genomes of phytoplanktonic protists and have a rearrangement of the p10 region preceding the p20, but otherwise have a similar activity profile to type I and type II metacaspases (Choi & Berges, 2013; Klemenčič & Funk, 2018). The conserved p20 and p10 regions together build up the core protease domain. They were designated p20 and p10 because activated enzymes typically show two bands at 20 kDa and 10 kDa, respectively, on SDS-PAGE gels. Most metacaspases require calcium ions (Ca^2+^) and a neutral pH for activity, with the notable exception of a sub-group of type II metacaspases present only in vascular plants that function optimally at mildly acidic pH (around pH 5.5) and do not require Ca^2+^ (Vercammen *et al*, 2004). Apart from their role in pathogen-induced PCD (in plants also called the hypersensitive response) (Coll *et al*, 2010; Watanabe & Lam, 2011a), metacaspases function in senescence, aging and protein aggregate clearing (Lee *et al*, 2010; Coll *et al*, 2014; Hill *et al*, 2014), clearance of cell corpses (Bollhöner *et al*, 2013), wound-induced damage associated molecular pattern signalling (Hander *et al*, 2019; Shen *et al*, 2019) and developmental cell death events (Bollhöner *et al*, 2013).

To date, two protein crystal structures of type I MCAs are published from *Saccharomyces cerevisiae* and *Trypanosoma brucei* (TbMCA-Ib) (McLuskey *et al*, 2012; Wong *et al*, 2012), and one type II calcium dependent MCA from *Arabidopsis thaliana* (AtMCA-IIa/AtMC4) (Zhu *et al*, 2020). The core of all three structures is a typical caspase/hemoglobinase alpha-beta-alpha sandwich fold (Aravind & Koonin, 2002). Differences exist within the type-specific regions: In the type I MCA structures, the N-terminal part of the protein binds in the active site, which explains why these enzymes are not fully active before removing the N-terminal pro-domain. However, it is not yet understood how Ca^2+^ activates the protease since the proposed Ca^2+^ binding sites are far away from the active site. Furthermore, the cysteine and histidine residues that make up the catalytic dyad are too far apart and, in a conformation where the catalytic histidine cannot act as base responsible for cysteine deprotonation, which is otherwise usually proposed as mechanism for other cysteine proteases like papains. This conundrum presents itself also in the caspases for which the catalytic mechanism is not completely explained (Ramoso-Guzmán *et al*, 2019). In the AtMCA-IIa structure, part of the linker region binds in the active site, which explains the need for auto-cleavage in this region for activation (Zhu *et al*, 2020). The resolved structures of AtMCA-IIa have no Ca^2+^ structure is observed, therefore it remains unclear where Ca^2+^ binds in type II MCAs, and how it triggers activation. Similar to the conundrum in type I MCAs, AtMCA-IIa structures also show an inactive form of the enzyme with the catalytic residues too far apart. In addition, for the Ca^2+^-independent metacaspases, a representative crystal structure is lacking, which further impedes the understanding of metacaspase activation mechanisms.

Lateral roots make up a significant part of the plant root system and ensure optimal nutrient absorption, water uptake and soil anchorage of plants. Their developmental trajectory is remarkable, as they originate from precursor cells deep within the main root and have to push through the terminally differentiated outer cell layers of the root. Lateral roots originate from pericycle cells that acquire lateral root founder cell identity and divide asymmetrically, a process that is tightly regulated by the plant hormone auxin. Under the control of auxin, these founder cells develop into a lateral root primordium (LRP), which then grows through the overlying endodermis and cortex until it eventually breaks through the epidermis and emerges from the primary root. Evidence has been provided that a tight integration of chemical and mechanical signalling between the LRP and the surrounding tissue is essential for its proper development while the overlying layers have to accommodate the outgrowth of lateral roots (Vermeer *et al*, 2014). In Arabidopsis, the pericycle-overlying endodermal cells represent a major obstacle for the developing lateral root primordium because of the presence of their lignified Casparian strips in the primary cell wall. Both endodermal cell elimination and accommodation due to cell shape changes have been reported to contribute to the outgrowth of primordia through the endodermis (Vermeer *et al*, 2014; Escamez *et al*, 2020). The Arabidopsis genome contains three type I and six type II MCAs (Uren *et al*, 2000; Vercammen *et al*, 2004; Tsiatsiani *et al*, 2011). The reported specific spatio-temporal transcriptional induction of *AtMCA-IIf/AtMC9* in the endodermis in front of an initiation LRP, suggested its involvement in PCD of the endodermis to clear the way of the developing primordium. However, most likely due to functional redundancy, single *atmca-IIf* mutants do not experience problems during lateral root emergence (Escamez *et al*, 2020).

Here, we took a chemical biology approach to address the role of metacaspases in lateral root formation. First, we solved the crystal structure of AtMCA-IIf, the sole Ca^2+^-independent metacaspase of Arabidopsis. We then developed an in vitro chemical screen and identified a series of small molecule metacaspase inhibitors with a minimal thioxodihydropyrimidine-dione (TDP) structure, some of which could specifically inhibit activity of AtMCA-IIf. Finally, we show that an AtMCA-IIf-specific TDP-containing inhibitor (TDP6) was effective at suppressing LRP emergence in vivo.

## Results

### Preparation of the AtMCA-IIf crystal structure, a representative of Ca^2+^-independent metacaspases

For structure determination of AtMCA-IIf, recombinant wildtype and catalytically inactive C147A mutant proteases with N-terminal His-tag (Vercammen *et al*, 2004) were expressed in *E. coli* and purified by IMAC and size exclusion chromatography, yielding 11-26 mg protein/L culture. No crystals were obtained of the wildtype protease, presumably due to autocleavage, but the C147A mutant did crystallize. Structures of two crystal forms were obtained, tetragonal P4322 and monoclinic P21, at 2.6 and 1.95 Å resolution, respectively. Apart from the crystal packing the two structures are practically identical, and the P21 structure refined at R/R_free_ values of 0.17/0.21 was chosen for analysis. The four protein chains in the asymmetric unit are very similar with RMSD of 0.18–0.19 Å. At position 271 we found an unintended Phe to Tyr mutation in all four chains, but it does not seem to induce any structural disturbances. Chain C is shown because it has lowest number of amino acid residues that are not seen in the structure. In total, 20 residues are missing from chain C, the 6 first N-terminal residues, 7 in the first gap (102-VKSAHPF-108) and 7 in a second gap (166-SSNISPA-172). Statistics from diffraction data processing and structure refinement are summarized in Appendix Table S1.

Type II metacaspases are built from a core domain comprising the p20 and p10 regions, a linker region, an autoinhibitory motif (AIM) bound at the active site, and a domain of unknown function (UNK; Fig. 1a). The core domain exhibits the typical caspase/hemoglobinase alpha-beta-alpha sandwich fold, with a central β-sheet, containing six strands in the order 213456 where strand 6 is antiparallel to the rest, surrounded by five α-helices and a β-hairpin motif (Fig. 1b). In the following, residue numbers refer to AtMCA-IIf unless otherwise indicated. The catalytic dyad residues His95 and Cys147 (mutated to Ala in the structure) are located in the p20 region after the β3 and β4 strands, respectively. The p20 region is followed by the linker, which is probably flexible as judged from the absence of electron density for six amino acid residues that could not be built here. Then comes the AIM embedded in the active site with the Arg183 side chain inserted in the S_1_ pocket (Fig. 1c), immediately followed by the UNK domain, built from three α-helices and a short 3_10_ helix, and finally the p10 region of the core domain.

**Fig. 1:**
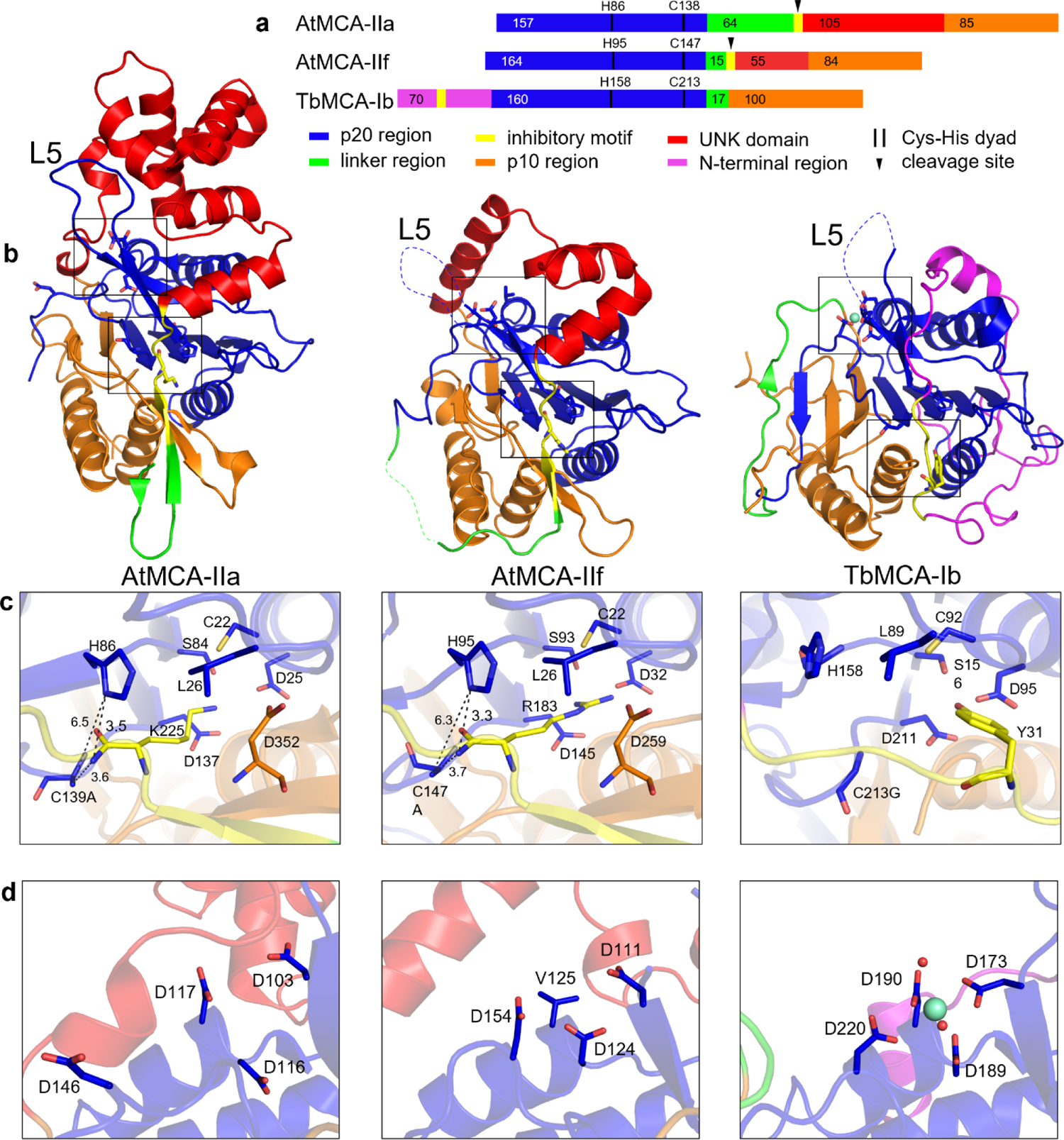
Comparison of the newly obtained AtMCA-IIf C147A crystal structure to available MCA structures. **a**, Scheme of the protein sequences drawn to scale and colored by region (color key in figure). Regions p20 and p10 where designated as described in [32]. The positions of the catalytic dyad His and Cys residues are indicated as well as the number of amino acids in each region. **b**, Overall structures of AtMCAII-a (PDB 6W8S; C139A mutation), AtMCA-IIf (PDB 8A53; C147A mutation), and TbMCA-Ib (PDB 4AFP; C213G mutation) color-coded by region. Selected residues are shown in stick representation, at the catalytic center (lower box), and at the putative calcium binding site (upper box). The position of loop L5 is indicated, which is visible in AtMCA-IIa but not in the two other structures. **c**, Catalytic center with autoinhibitory motifs bound. (d) Putative calcium binding site, with Sm^3+^ bound in the TbMCA-Ib structure (right).

### Structure comparison of AtMCA-IIf reveals an overall similarity with AtMCA-IIa

Superposition of the AtMCA-IIf and AtMCA-IIa structures gives a backbone RMSD value of 1.1 Å, meaning the proteases are overall similar, but with distinct features. The core domain is very similar in size and shape with RMSD value of 0.81 Å. Both proteases contain an AIM embedded in the active site, albeit with different sequences (Fig. 1c). In AtMCA-IIf, Arg183 (180-ITS<u>R</u>ALP-186) is a key residue that has to be cleaved in order for the protease to be active, whereas Lys225 (222-AKD<u>K</u>SLP-228) is the corresponding residue in AtMCA-IIa (Vercammen *et al*, 2004;Watanabe & Lam, 2011b). The linker region (from 165 to 179) is much shorter in AtMCA-IIf with 15 amino acid residues (6 of which are missing in the structure), compared to around 70 residues in AtMCA-IIa where 60 are missing from the current structure. The UNK domain (position 187 to 241), not present in the type I structures, is also shorter in AtMCA-IIf. It contains only helices, 4 in AtMCA-IIf and 8 in AtMCA-IIa, with 55 and 105 amino acid residues, respectively (Fig. 1b).

### Binding of the autoinhibitory motif (AIM) in the active site resembles AtMCA-IIf substrate preference

At the catalytic center, the AIM is positioned with its backbone carbonyl of Arg183 placed between the catalytic dyad residues (His95 and Cys139Ala; Fig. 1c). Thus, with the catalytic residues on opposite sides of the peptide bond to be cleaved, the histidine will not be in direct contact with and cannot act as activating proton acceptor to the catalytic cysteine (Cys147 in AtMCA-IIf wildtype). Instead, one of the His95 nitrogen atoms (ND1) points directly at the backbone nitrogen of Ala184 at hydrogen bond distance (3.4 Å), suggesting a role for His95 in the first, acylation step of the reaction. The backbone carbonyl oxygen of Arg183 is pointing into a presumed oxy-anion hole forming two hydrogen bonds with the backbone nitrogens of Gly96 and Ala147 (catalytic Cys147 in the wildtype enzyme) (Appendix Fig. S1a).

Whereas the AIM is quite different in TbMCA-Ib, it is similar in the type II enzymes. The sidechain of Arg183 (and Lys225 in AtMCA-IIa) is entirely buried in the adjacent S_1_ pocket, built from the same amino acids in type I and type II, except for one Asp (Asp259) from the p10 region in type II. In TbMCA-Ib this space is instead occupied by Tyr31 of the N-terminal region (Fig. 1c). In the N-terminal direction from the cleavage site, four additional amino acid residues (179-TITS-182) form a beta-strand, bound antiparallel with the 258-261 beta-strand in p10. Their sidechains are exposed and no well-defined pockets are distinguished, except for a predominantly hydrophobic S_4_ pocket, suggesting a preference for a hydrophobic residue at the P4 position of substrates (Appendix Fig. S2). In the C-terminal direction, no S_1_’ pocket can be distinguished and all residues in the vicinity are the same in AtMCA-IIf and AtMCA-IIa (Appendix Fig. S1b). There is a clear hydrophobic S_2_’ pocket though, harbouring leucine in both structures, where several residues are conserved while some are different (Asn24/Ala17, Phe119/Met111, Ala189/Thr231, Val190/Leu232; AtMCA-IIf/AtMCA-IIa numbering). Also, the first helix of the UNK domain (187-197) is oriented slightly different between the two AtMCAs, which may influence the S_2_’ pocket formation. Next, there is no clear S_3_’ pocket. A proline sidechain points out in solution in both structures, after which the peptide chain turns about 90 degrees and continues into the first α-helix of the UNK domain.

To assess substrate specificity, a positional proteomics approach was used to screen the root proteome of Arabidopsis plants for proteins that are differentially cleaved in AtMCA-IIf loss- or gain-of-function mutants by N-terminal combined fractional diagonal chromatography (COFRADIC; Staes *et al*, 2011). Two analyses were done in which the root N-terminome of *atmca-IIf* loss-of-function (knockout, KO) plants was compared to wild type (WT) or AtMCA-IIf gain-of-function (overexpression, OE) plants (Appendix Fig. S3a). Additionally, an *in vitro* analysis was performed in which an *atmca-IIf* KO root proteome was treated with recombinant AtMCA-IIf. Identification of so-called neo-N-terminal peptides revealed a subfraction of proteins in the Arabidopsis proteome that are potentially cleaved by AtMCA-IIf and the cleavage sites in those proteins (Appendix Fig. S3b-c; Appendix Table S2). In line with our previous findings on metacaspase cleavage sites, 70% and 72% of the significant differentially abundant neo-N-terminal peptides were generated after Arg or Lys in the KO/OE *in vivo* and the *in vitro* analysis, respectively (Vercammen *et al*, 2006). In the KO/WT *in vivo* study this was only 13%, which is likely due to the lower amounts of AtMCA-IIf in the WT samples compared to OE and *in vitro* studies. Enriched amino acid sequences within the P4 to P4′ position spanning the cleavage site (nomenclature according to Schechter & Berger, 2009) of the Arg/Lys-cleaved proteins display a cleavage signature that is similar to the one previously described for AtMCA-IIf in protein substrates from 2-day-old seedlings (Tsiatsiani *et al*, 2013) (Appendix Fig. S3e). Based on the crystal structure, we were now able to match the substrate cleavage specificity to the active site pockets of AtMCA-IIf. The P1 site entailing Arg or Lys obviously fits the negatively charged S1 pocket. No clear explanation is at hand for the enrichment of basic amino acid residues in P3, neither for acidic side chains in P1’ position, as clear pockets were missing from the AtMCA-IIf structure. Convincingly, mostly hydrophobic amino acids were found at the P2’ position that can fit the hydrophobic S2’ pocket. To a lesser extent there was a fit for the hydrophobic S4 pocket, where only in the previous study by Tsiatsiani *et al*. there was an enrichment for Ala in P4 (as well as Glu and Val; Appendix Fig. S3e). Altogether, an AtMCA-IIf substrate cleavage signature emerged wherein P3 and P1 are occupied by basic, P1’ by acidic, and P4 and P2’ by hydrophobic amino acid residues. This is reflected at least in the active site pockets S1 (acidic) and S2’ (hydrophobic) from the newly acquired AtMCA-IIf crystal structure.

### Absence of calcium binding sites in AtMCA-IIf

To assess the calcium independency of AtMCA-IIf, we compared the proposed Ca^2+^ binding sites between the available 3D crystal structures (Fig. 1d). TbMCA-Ib has a Sm^3+^ ion bound as a Ca^2+^ surrogate and this site is almost identical in both type I structures (McLuskey *et al*, 2012; Wong *et al*, 2012). The Sm^3+^ ion is coordinated by four aspartic acid residues and two water molecules, while in AtMCA-IIf only three of those four are conserved. In the fourth position a valine instead of an aspartic acid residue is present, which could be connected to the AtMCA-IIf Ca^2+^-independency (Fig. 1d). However, as stated previously by Zhu *et. al.*, the equivalent site in the AtMCA-IIa structure is blocked by a loop at the end of the UNK domain, just before p10 (Zhu *et al*, 2020). Instead, they proposed an alternative Ca^2+^ binding site composed of four negatively charged amino acids in a row (96-EDDD-99) at the turn of loop L5 that is embedded in a positively charged pocket of the UNK domain (Fig. 1b). Negatively charged residues are conserved here in both type I and II of Arabidopsis (three in AtMCA-Ia-c and four in AtMCA-IIa-e), except for AtMCA-IIf, which is also lacking the part of the UNK domain that supports loop L5 in AtMCA-IIa (Appendix Fig. S4). The corresponding loop of AtMCA-IIf is disordered and not visible in the crystal structure (102-VKSAHPF-108) and does not contain a similar cluster of negatively charged residues. These structural differences could explain why AtMCA-IIf is not activated by Ca^2+^.

### A chemical screen finds metacaspase small molecule inhibitors with a common thioxodihydropyrimidinedione (TDP) scaffold

To overcome potential metacaspase functional redundancy and phenotypic plasticity as a result of stable metacaspase mutations, we aimed for a chemical approach to inhibit the enzymatic activity in situ. Therefore, we developed a high-throughput adaptation of an *in vitro* biochemical assay based on cleavage of a fluorescently-labeled VRPR-AMC tetrapeptide by recombinant AtMCA-IIf (Vercammen *et al*, 2004, 2006). The miniaturized assay was used to screen a chemical library of 10,000 small organic compounds at a concentration of 10 µM to identify inhibitors of AtMCA-IIf (Fig. 2a). The screen was performed twice against the 10,000 compounds and together led to 177 compounds that showed at least 75% inhibition. These 177 compounds were then selected for a third screen. Together, 81 compounds were identified that reduced the metacaspase activity on average with at least 75% in all three screens (Appendix Table S3). Investigation of the chemical diversity of the inhibitors by performing a clustering based on the compounds’ chemical structures revealed an enrichment for hits containing a thioxodihydropyrimidinedione (TDP) scaffold (Fig. 2b; Appendix Table S3). This TDP substructure was found in 30 (37%) of the hit compounds, compared to 79 (0.8%) present in the entire screening library, which made us to prioritize the TDP-containing hits for further investigation.

**Fig. 2:**
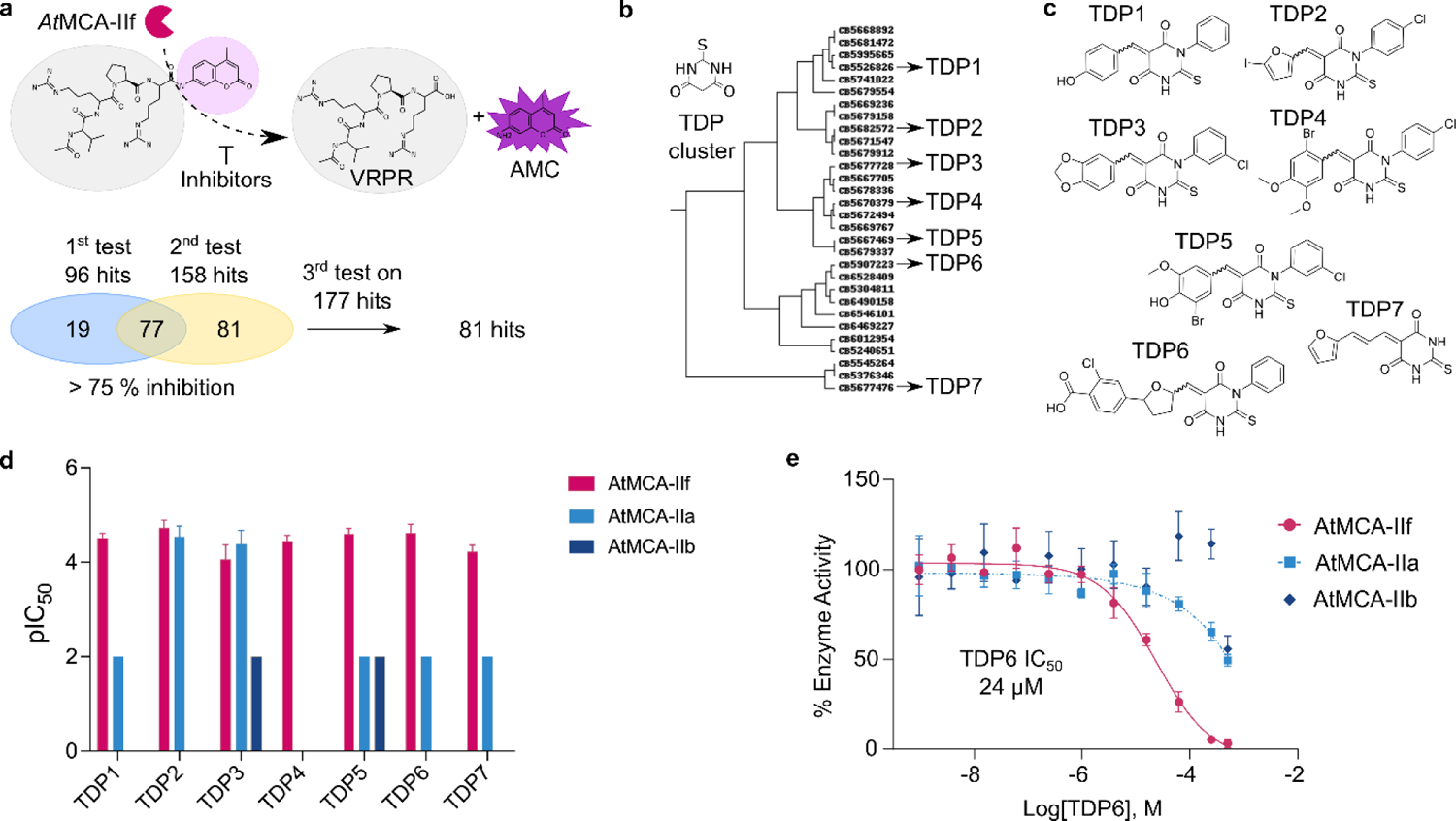
A chemical screen reveals a cluster of AtMCA-IIf protease inhibitors that share a TDP substructure. **a**, Schematic overview of the chemical screen against AtMCA-IIf. The full screen (10.000 compounds) was performed twice resulting in the identification of 96 and 158 compounds with an inhibitory effect larger than 75%. The resulting 177 compounds were re-tested and 81 compounds were retained with an average inhibition larger than 75% in the 3 tests. **b**, Structure-based clustering of chemical compounds with 75% AtMCA-IIf inhibitory effect. From 81 final hits, 30 compounds contain a thioxodihydropyrimidinedione (TDP) substructure. **c**, Chemical structures of the seven representative compounds from each TDP subcluster based on their inhibitory effect against the VRPR-ase activity of AtMCA-IIf. **d**, Inhibitory activity of the TDP compounds against three metacaspases expressed as pIC50. Compounds that showed some inhibitory activity (IC50> 200 µM) at the highest tested concentrations are set to pIC50=2 to illustrate the inhibitory difference between the enzymes. **e,** Dose-response curve for inhibitory activity for TDP6 against three metacaspases, obtained from the activity-based in vitro assay.

### Dose-response analysis of TDP-containing compounds to probe selectivity towards AtMCA-IIf, AtMCA-IIa and AtMCA-Iib

Based upon their inhibitory activity and chemical diversity, seven compounds containing the TDP scaffold were selected for a full dose-response analysis (TDP1 to TDP7; Fig. 2c; Appendix Fig. S5a; Appendix Table S4). In addition to AtMCA-IIf, the inhibitory activities of the compounds were evaluated on AtMCA-IIb/AtMC5, because of its overlapping gene expression pattern in roots (Appendix Fig. S8), and AtMCA-IIa, the most abundant and relatively well-studied Arabidopsis metacaspase (Watanabe & Lam, 2011a, 2011b; Fortin & Lam, 2018; Hander *et al*, 2019; Zhu *et al*, 2020). Measurements were performed on commercially available compounds, of which the chemical structures had been confirmed by nuclear magnetic resonance (NMR) characterization (Appendix Table S5). The compounds all carry different aromatic or heteroaromatic moieties linked to the TDP scaffold via a carbon-carbon double bond, generating two possible isomers, depending on the substitution pattern (E and Z). Before testing, the ratio between the E and Z isomers was determined by ^1^H-NMR to be 1:1 for all TDPs, except TDP7 that only was found in the E configuration (Appendix Table S5). Thus, six out of seven TDPs are a mixture of two compounds with the same chemical formula (diastereomeric). The full dose-response analysis of the TDPs towards AtMCA-IIf corresponded well with the screening data, as all hits showed dose-dependent inhibition towards the protease, with IC_50_ values ranging from 19 to 86 µM (Fig. 2d; Appendix Fig. S5a; Appendix Table S4). A comparison of the inhibition profiles between the proteases showed that the majority of the TDPs selectively inhibited AtMCA-IIf. Only two of the compounds (TDP2 and TDP3) inhibited the activity of AtMCA-IIa in a dose-dependent manner, exhibiting an inhibitory effect similar as for AtMCA-IIf, while additional four showed some activity at the highest concentrations. The compounds showed even less inhibitory activity against AtMCA-IIb; only two (TDP3 and TDP5) where active at the highest tested concentrations. Investigation of the inhibition profile towards AtMCA-IIf, revealed a steep dose-dependency (Hillslope >1) for some of the TDPs. This might be related to the diastereomeric mixture of the TDP compounds, where the two compounds possible can interact differently with the protease causing the steepness of the curve. We developed an orthogonal in vitro assay to probe inhibition of metacaspase activities based on the cleavage of a substrate protein, PROPEP1 (Hander *et al*, 2019; Shen *et al*, 2019; Zhu *et al*, 2020). TDP6 was selected as a representative of the inhibitors with the lowest IC_50_ of the compounds that are selective towards AtMCA-IIf in the VRPR-AMC cleavage assay. Increased cleavage of a recombinant GST-TEV-PROPEP1 fusion protein by AtMCA-IIf correlated with decreased concentrations of TDP6 in the reaction mixture (Appendix Fig. S6a-b), while this correlation was obscure for AtMCA-IIa and AtMCA-IIb (Appendix Fig. S6c-d). In conclusion, several new TDP-containing small molecules were found to inhibit AtMCA-IIf enzymatic activity in vitro, without causing inihibition of the two other AtMCAs of the same type.

### Molecular docking simulations to the crystal structure of AtMCA-IIf suggest binding of TDP compounds to the active site pocket

To understand how the TDPs could interact with AtMCA-IIf and inhibit the catalysis, we performed molecular docking to the active site of AtMCA-IIf of TDP6, as selective inhibitor (Fig. 2d,e), and TDP2 and TDP3, which inhibit both AtMCA-IIf and AtMCA-IIa (Fig. 3; Appendix Fig. S7). The autoinhibitory motif was excluded from AtMCA-IIf to expose the active site prior to docking. Since the TDPs were tested experimentally as a mixture of isomers, both the E- and Z-configurations of TDP2, TDP3 and TDP6 were considered. The molecular docking revealed multiple poses of the TDP inhibitors due to the E- and Z-configurations and the apparent symmetric nature of the TDP scaffold with its phenyl- and benzylidene substituents. Two molecular interaction types appeared to be the driving force for the generated binding poses in the active site of *At*MCA-IIf: (i) an aromatic moiety of the inhibitors (the phenyl- or the benzylidene substituent) projected into the S1 pocket of the protein for electrostatic and dispersion interactions and (ii) hydrogen bonding interactions between a carbonyl oxygen of the TDP scaffold and one or more –NH of the protein, (the –NH+ of His95, and the backbone –NH’s of Gly96 and Cys147). Note that in the crystal structure where the autoinhibitory motif is included, (i) corresponds to the pocket where the side chain of Arg183 is bound and (ii) is the presumed oxy-anion hole where the peptide carbonyl oxygen of Arg183 is located. In the molecular dockings, it was the type of substituents on the phenyl- or the benzylidene moieties together with the E- and Z-configurations that determined which one of the aromatic moieties was projected into the S1 pocket and thus which of the two carbonyls of the TDP scaffold that formed the hydrogen bonding interactions. In addition to the key interactions that were observed for all docked compounds, the TDP inhibitors with E-configuration placed the other aromatic substituent along the β4 sheet (Fig. 3a), while the docking poses for the Z-isomers varied more. In many docking poses, the other aromatic moiety of the Z-isomers was projected out to the solvent in the opposite direction of the β4 sheet, or in a few poses in the S2’ pocket where Leu185 of the autoinhibitory motif is located. The multiple alternatives of binding poses for this class of inhibitors might explain its unpronounced structure-activity relationship. Inspection of the docking poses showed that the larger benzylidene substituent of E-TDP6 had a more extensive interaction surface with the β4 sheet directed towards the L5 loop, that connects β4 with β5, compared to the non-selective TDP2 and TDP3 (Fig. 3a; Appendix Fig. S7). Interestingly, while the amino acid residues forming the β4 and β5 sheets are highly similar between the two proteases, there are large differences in the L5 loop. The ten amino acid residue Pro100-Lys109 loop of AtMCA-IIf shows no sequence similarity, except for Pro91 in the L5 loop of AtMCA-IIa (Fig. 3b). Instead, there are significant differences that likely influence the dynamics of the loops and thus also the character of the β-sheet surfaces. The loop of AtMCA-IIa comprises two glycines (Gly95 and Gly101), while the loop of AtMCA-IIf instead has an additional proline (Pro107). Thus, we propose that the loss of activity for TDP6 towards AtMCA-IIa, while TDP2 and TDP3 maintain inhibition, was because the β4-sheet of AtMCA-IIa could not accommodate the larger substituent of TDP6 in a similar manner as for AtMCA-IIf due to differences in the dynamic pattern of the β-sheet surfaces.

**Fig. 3:**
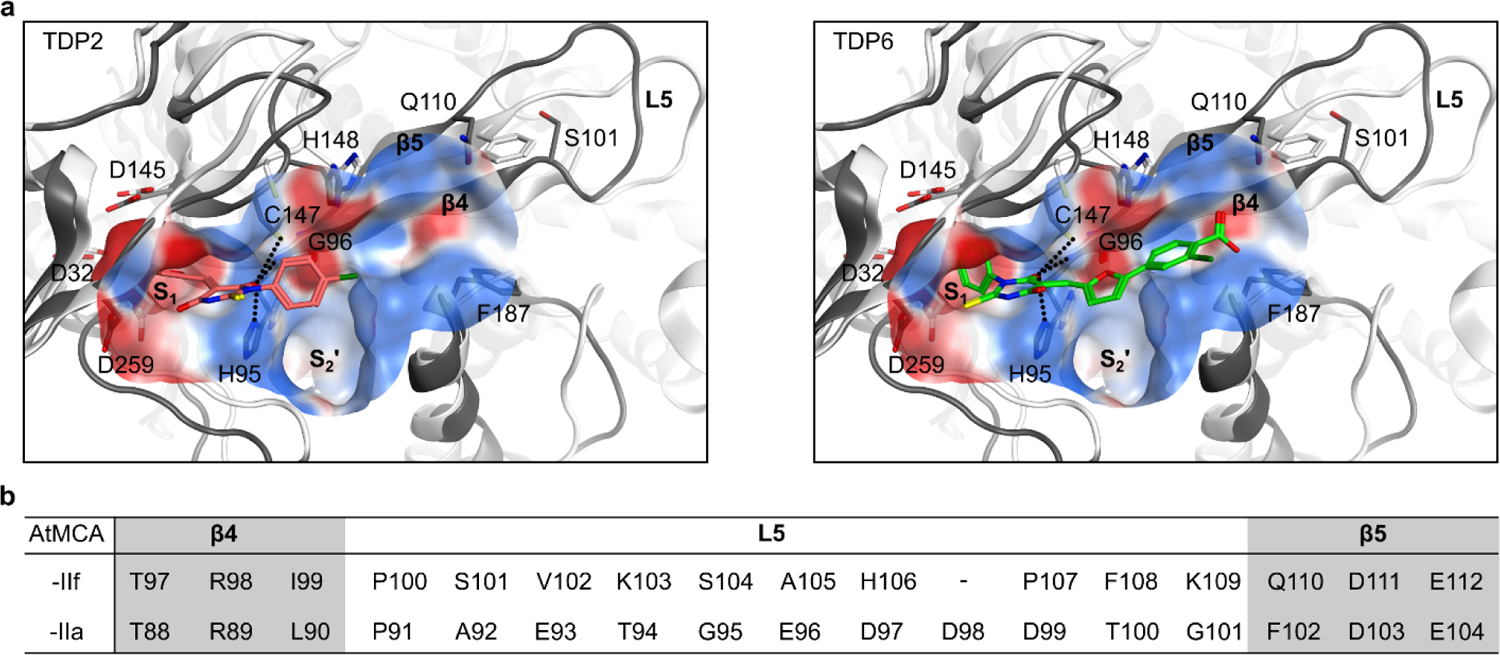
Potential binding modes of TDP6 based on docking to AtMCA-IIf. **a**, The tertiary structure and selected residues of AtMCA-IIf (PDB 8A53) are displayed in dark grey and AtMCA-IIa (PDB: 6W8S) in white. One of the aromatic substituents of the thiobarbituric acid scaffold of the E-configuration of TDP2 and TDP6 is placed in the S1 pocket of AtMCA-IIf, and the second aromatic substituent is placed along the β4 sheet. Possible hydrogen bonding interactions to the carbonyl oxygens of the thiobarbituric acid moiety are shown with dotted lines. The electrostatic surface of the active site is displayed. The L5 loop of AtMCA-IIf has been modelled for visualization purposes. Residue numbers for AtMCA-IIf are shown. **b,** Comparison of the sequence of the L5 loop of AtMCA-IIf and AtMCA-IIa.

### AtMCA-IIf spatiotemporal expression pattern suggests a role in lateral root emergence

As demonstrated earlier (Escamez *et al*, 2020), *AtMCA-IIf* is expressed in the endodermis in cells overlying developing lateral root primordia (LRP; Appendix Fig. S8a), which suggests a role in the emergence of lateral roots through the endodermis (Fig. 4a). Furthermore, we studied the spatiotemporal expression of the other Arabidopsis metacaspase genes in roots using promoter GUS-GFP reporter lines. Similar to *AtMCA-IIf*, *AtMCA-IIb* is expressed in endodermal cells overlying developing primordia (Appendix Fig. S8b). Besides *AtMCA-IIc/AtMC6*, all metacaspase genes were expressed in the root, although with distinctive expression levels and spatial patterns (Appendix Fig. S9). The expression pattern of *AtMCA-IIf* was investigated in more detail with a translational promoter AtMCA-IIf-GFP fusion construct confirming expression in the endodermis during the first stages of lateral root development (Fig. 4b). As previously reported (Bollhöner *et al*, 2013), AtMCA-IIf was also expressed early on in the xylem strands (adjacent to the stage II LRP, Fig. 4b; Appendix Movie S1), which disappears in the later stages of LRP development during which the xylem loses the GFP signal (Stage V LRP, Fig. 4b; Appendix Movie S1). Next to the tight engulfment of the endodermal cell(s) surrounding the LRP, a striking partitioning of the AtMCA-IIf-GFP fusion protein in the endodermis to the side of the cell bordering the developing lateral root could be discerned (Fig. 4c). LRP emergence in T-DNA knock-out lines of *atmca-IIb*, *atmca-IIf*, and the double *atmca-IIb atmca-IIf* mutant were measured (Appendix Fig. S10a,b). Surprisingly, no significant differences were observed between wild-type plants and single or double knock-out lines in the ratio of non-emerged lateral roots (Appendix Fig. S10c). This confirms previous findings that phenotypes in single or double mutants have no effect on LRP emergence possibly because of functional redundancy of metacaspases (Escamez *et al*, 2020).

**Fig. 4:**
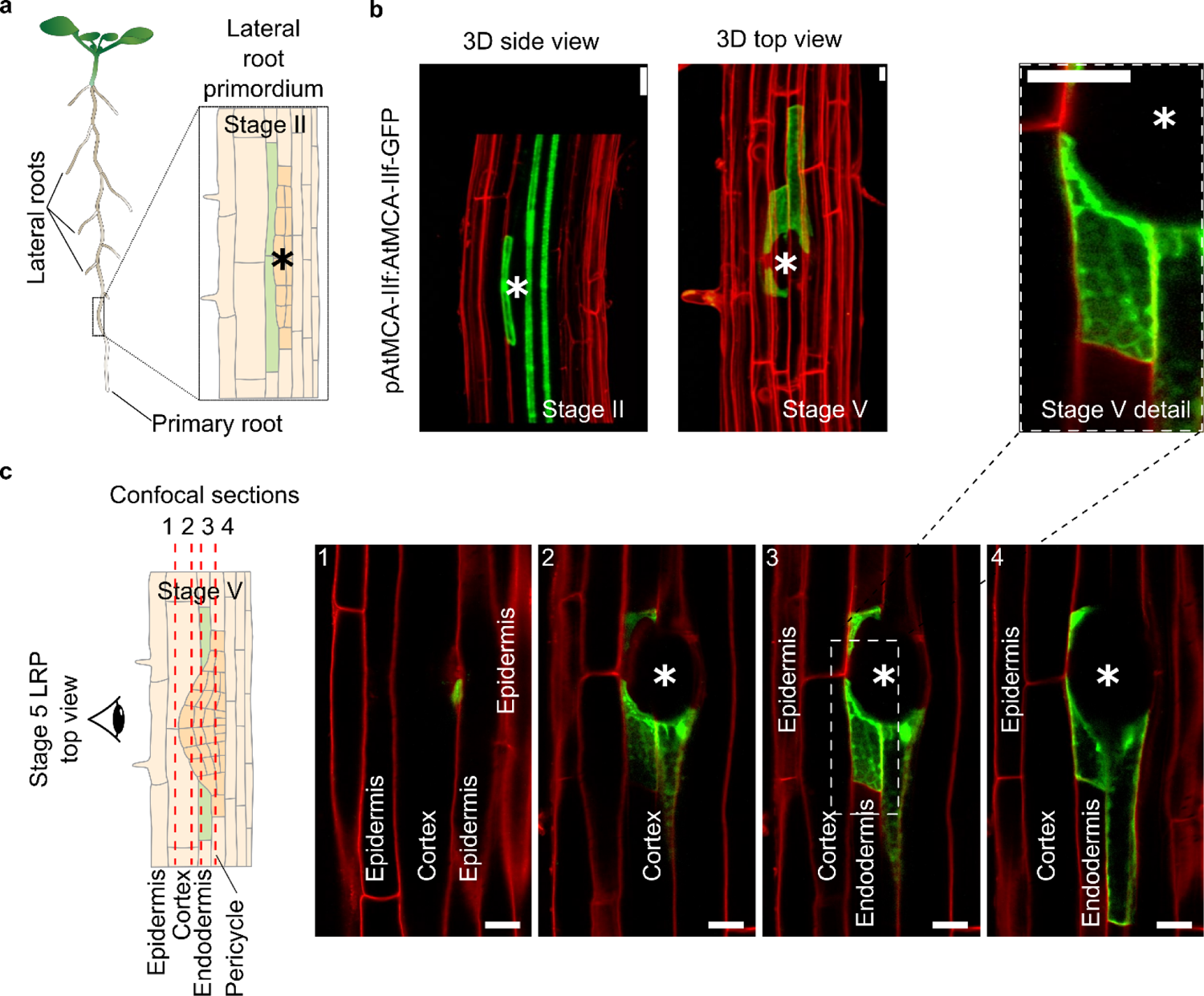
AtMCA-IIf is specifically expressed in endodermal cells overlaying the lateral root primordium in Arabidopsis. **a**, Schematic overview of the site of lateral root development in the primary Arabidopsis root. **b**, A 3D reconstruction of pAtMCA-IIf:AtMCA-IIf-GFP fluorescence from confocal sections of stage 2-3 and stage 5 lateral root primordia. The primordia is indicated with an asterisk (*). Cell walls are stained with PI (red). **c**, Confocal sections through the tissue layers overlaying the stage 5 LRP. Scalebar is 20 µm.

### Inhibition of metacaspase activity by TDP6 suppresses lateral root emergence

To assess the phenotypic effect of metacaspase inhibition on LRP development, 8-day old Arabidopsis seedlings were analysed after a 5-day growth period in media supplemented with 50 µM of the inhibitors TDP1-7. Several compounds decreased the number of lateral roots, with TDP3 and TDP6 having the most pronounced effect on lateral root density (Fig. 5a). Unlike TDP3, TDP6 did not have a significant effect (p<0.05) on primary root length, indicating that TDP6 was a more specific inhibitor of lateral root development (Fig. 5a,b; Appendix Fig. S11). Interestingly, TDP2, which showed a high inhibitory activity towards both AtMCA-IIf and AtMCA-IIa (Fig. 2d; Appendix Table 4), affected lateral root development only to a minor degree. This suggests that factors unaccounted for in an in vitro assay, such as chemical stability and tissue uptake, are critical for the in vivo effect of a chemical probe.

**Fig. 5:**
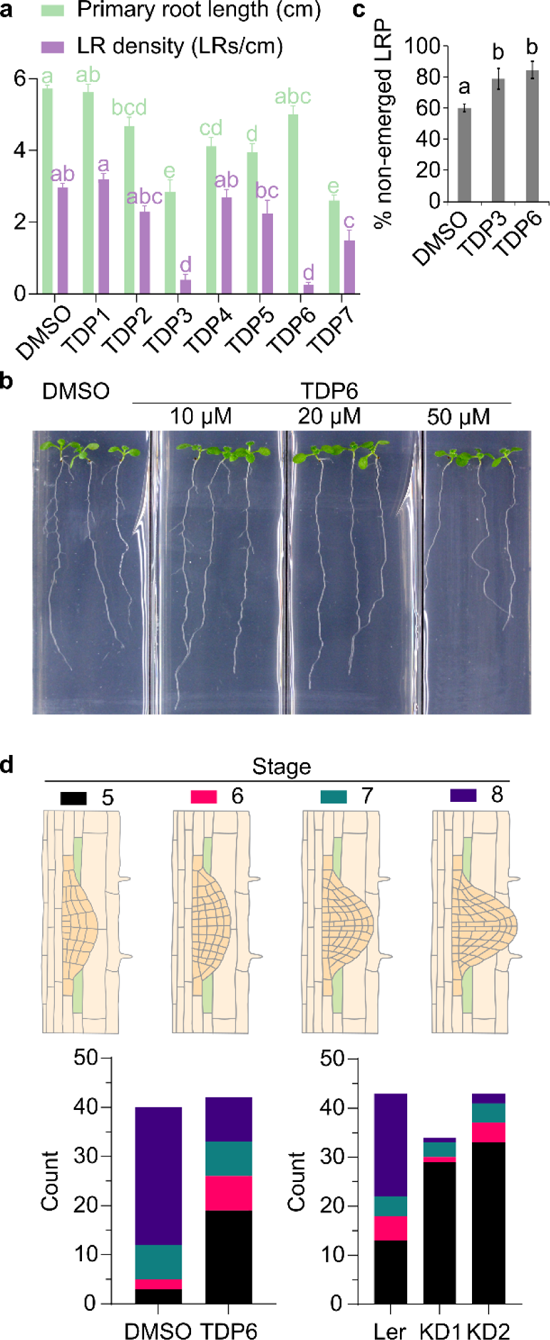
TDP6 inhibits lateral root emergence. **a**, The effect of the selected TDP-containing inhibitors on primary root length and lateral root density compared to control (DMSO). Error bars represent SEM, treatments were compared separately for primary root length and lateral root density by one-way ANOVA post-hoc Tukeýs test (p*<*0.05). **b**, Visual phenotype of TDP6 treatment on lateral root growth. **c**, The number of non-emerged lateral root primordia (LRP) is increased by 20% on average by TDP3 and TDP6. Error bars represent SEM; treatments were compared with one-way ANOVA post-hoc Tukeýs test (p*<*0.0001). **d**, Stages of LRP development scored in a root bending assay for TDP6 and two metacaspase II family amiRNA knock-down lines (KD1 and KD2) compared to their controls (DMSO and wild type Landsberg erecta seedlings, respectively).

A detailed assesment of the TDP3 and TDP6 effect on lateral root emergence showed that the proportion of non-emerged primordia in treated seedlings was on average 20% higher compared to the untreated control (Fig. 5c). Lateral root primordia go through various stages of development (Malamy & Benfey, 1997), which can be conveniently scored by a root bending assay in which seedlings are turned 90° after which the root tips grow towards the shifted vector of gravity, resulting in a bend in the root and the development of a LRP in the bend. In this assay, primordia emergence appeared to have slowed down as they accumulate primarily at stage 5 in TDP6 treated seedlings (Fig. 5d). To confirm these findings by genetic means, we reduced type II metacaspase gene expression with an amiRNA construct directed against *AtMCA-IIa,b,c,d* and *-IIf* in an *AtMCA-IIe/AtMC8* T-DNA background line (Appendix Fig. S12a,b). Similar to TDP3 and TDP6 treatment of wild type seedlings (Fig. 5c), two independent knock-down lines (KD1 and KD2) contained more non-emerged lateral root primordia than wild type (Appendix Fig. S12c), although the effect was more pronounced with the TDP compounds. Detailed analysis of lateral root primordia stages showed a significantly increased number of lateral root primordia (p<0.05) that have not yet protruded the endodermis (stages 1-3; Appendix Fig. 12d), suggesting that the endodermis is the restraining tissue for lateral root emergence when type II metacaspase levels are down-regulated. These findings are recapitulated in a root bending assay of the KD1 and KD2 lines (Fig. 5d). Unlike with the knock-down lines, inhibition of lateral root emergence can be reverted by transfer of seedlings from TDP6 to control medium (Appendix Fig. S13a,b). Importantly, a reversal was observed in the part of the root that was previously exposed to TDP6 (Appendix Fig. S13c), underscoring the importance of TDP6 as a versatile tool to tune lateral root emergence. In conclusion, where genetic ablation of AtMCA-IIf in previous and current research failed to uncover an effect on lateral root development, application of the novel small molecule AtMCA-IIf inhibitor TDP6 reveals a positive effect of metacaspase on LRP emergence.

## Discussion

### The AtMCA-IIf crystal structure has led to an improved understanding of metacaspase activity

We report here the crystal structure of a Ca^2+^-independent type II metacaspase, AtMCA-IIf, which was needed in order to obtain a better understanding of metacaspase activity. At the same time, this study raised further questions about the catalytic mechanism of the cysteine protease clan CD that contains, among others, metacaspases, paracaspases such as Mucosa-associated lymphoid tissue lymphoma translocation protein 1 (MALT1), caspases, legumain and separase. It was concluded already by Polgár (2004) and is declared in the MEROPS database (Rawlings *et al*, 2018) that the His-Cys catalytic dyad acts with a mechanism distinct from the Cys-His dyads in clan CA (papain and homologs). Yet, several later studies have used the mechanism of papain to describe the role of the His-Cys dyad in clan CD proteases. In papain, the Cys and His residues are positioned in direct contact with each other, on the same side of the peptide bond to be cleaved, and the mechanism has been well established (Polgár, 2004). The role of the histidine residue is to act as proton acceptor to the cysteine, and the deprotonated, negatively charged form of the cysteine makes a nucleophilic attack at the peptide carbonyl carbon, which initiates cleavage and formation of the covalent acyl-enzyme intermediate. In clan CD on the other hand, the His and Cys residues are positioned on opposite sides of the scissile peptide bond, making it unlikely that the His residue can act as a proton acceptor for deprotonation of the catalytic Cys residue. It may be argued that the catalytic centre may undergo substantial conformational changes during the reaction, but such changes have not been reported for any clan CD proteases. Like in clan CA proteases, it is well established that the catalytic cysteine forms a covalent bond with the peptide carbonyl in the acyl intermediate. However, QM/MM simulations of the reaction mechanism for human legumain indicated that the cysteine needs to be in its protonated, neutral state for a productive nucleophilic attack and rupture of the scissile peptide bond (Elsässer *et al*, 2017). In this proposed catalytic mechanism, a water molecule is bound by the His residue in the enzyme-substrate complex, but in the structures of AtMCA-IIf and AtMCA-IIa there is no apparent room to host a water molecule at the corresponding position. It is thus uncertain how the His residue participates in the first acylation step. In the second hydrolysis step, the role of the His residue is presumably to bind, orient and activate the catalytic water by helping to shuttle one proton from the water molecule to the carbonyl oxygen of the acyl-enzyme intermediate.

For AtMCA-IIa, no major structural differences were noted at the active site between the inactive C139A mutant and wildtype proteases upon treatment of wildtype crystals with Ca^2+^, apart from the disappearance of electron density for Lys255 (presumably due to cleavage) (Zhu *et al*, 2020). Also, the positions of the His-Cys dyad-bearing loops are very similar when we superpose a wide range of clan CD protein structures, inactive mutants and zymogen forms as well as activated enzymes with trapped covalent intermediates (Appendix Fig. S14), indicating that zymogen activation does not involve large movements of the His-Cys dyad residues for recruitment of the catalytic machinery. Rather, the machinery is already in place and preconfigured for peptide bond cleavage. The complementarity of the AIM sequence to the active site suggests that the AtMCA-IIa and AtMCA-IIf structures may be close to the conformation of the Michaelis complex, i.e. how the substrate is bound for cleavage. If this is the case, why is the AIM not cleaved then in the zymogen state of the enzymes, or rather, why is it so slow? The autoinhibition by the AIM of type II metacaspases is analogous to the function of the Bovine pancreatic trypsin inhibitor (BPTI). BPTI binds to trypsin with high affinity, in the position for peptide bond hydrolysis, but is hydrolyzed ∼10^11^ times more slowly than small peptide substrates. The question of slow hydrolysis was addressed in a QM/MD simulation study of the binding and reaction of BPTI with trypsin (Peräkylä & Kollman, 2000). The calculations showed higher activation free energies for BPTI corresponding to 10^6^-10^9^ decrease in catalytic rate, due to a strong and rigid binding that makes the transition state formation more difficult. The authors also hypothesize that the remaining 10^2^-10^5^ decrease in rate is because the peptide bond in BPTI, once cleaved, is more likely to re-ligate than to be further hydrolyzed to a product that dissociates from the enzyme. We propose a similar inhibitory function of the AIM in type II metacaspases as for BPTI. Strong and rigid binding of the AIM sequence on both sides of the catalytic center may hold both the peptide carbonyl and nitrogen atoms so firmly in place that they are less likely to undergo the conformational changes required for bond cleavage. The activation energy barrier will thus be very high and the reaction rate very low. Furthermore, if the peptide bond between Arg183 and Ala184 is cleaved, the new N-terminal amino group may block the access for a water molecule to attack the acyl-enzyme intermediate and complete the hydrolysis, while the amino group itself will be well positioned for nucleophilic attack and relegation of the peptide bond. An additional feature of the AIM is that it is covalently linked to the rest of the protein. If hydrolytic cleavage does occur, and the newly formed ends dissociate from the active site, their effective concentrations will be very high, and they are likely to re-bind and re-ligate. Stronger binding of the non-cleaved substrate than after cleavage will affect the equilibrium of the reaction and favor non-cleaved over cleaved AIM sequence. Nevertheless, hydrolysis does seem to happen at low rates also in the absence of Ca^2+^ or low pH as indicated by (i) The fact that both recombinant AtMCA-IIa and AtMCA-IIf are partially autoprocessed when produced in *E. coli* (Vercammen *et al*, 2004), and (ii) the presence of a p20 band on western blots against AtMCA-IIa and AtMCA-IIf from plant protein extracts in steady state (Watanabe & Lam, 2011a; Belenghi *et* al, 2007; Hander *et al*, 2019). The conservation of an LPL/F motif in the AIM sequence across type II metacaspases suggests an important role (Appendix Fig. S4). The first Leu of this motif is deeply embedded in the S2’ pocket and is represented by Ile and Met in some homologs, while the Pro is strictly conserved and is partially exposed at the S3’ site (Appendix Fig. S1b and S2a). After the Pro residue, the peptide chain makes a sharp turn into the first alpha helix of the UNK domain, and the N-terminal end of the helix is anchored by a conserved hydrophobic residue (L/I/Y/F). We hypothesize that the binding of this motif is crucial for autoinhibition and that the function of the UNK domain is to keep this region firmly bound at the active site to prevent cleavage of the AIM. We also speculate that the activation mechanism involves increased dynamics of the UNK domain, at least of the first alpha helix, leading to larger conformational freedom of the AIM so that it is more readily cleaved. In AtMCA-IIf, an additional cleavage site was found in the UNK domain in the degradome study by Tsiatsiani *et al*. (2013) that may play a role here. The identified fragment shows cleavage at Arg216 in a surface turn between the second alpha-helix and the short 3-10 helix on the opposite side of the UNK domain from the active site. The sequence cleaved is LFGR(216)DAGLKF. If AtMCA-IIf is cleaved both at Arg183 of the AIM and at Arg216 of the UNK domain, the resulting 33-residue peptide could thus be free to detach and diffuse away from the activated enzyme. Similarly in AtMCA-IIa, a cleavage site has been identified at Lys249 in the UNK domain (Hander *et al*, 2019). Cleavage there and at Lys225 in the AIM will generate a 24-residue peptide comprising the 1st helix and the following loop.

Lastly, the UNK domain in AtMCA-IIa contains a rather large number of charged residues (11 Lys, 2 Arg, 7 Asp, 7 Glu) that stabilize the structure through numerous salt bridges. Several positive residues are buried within the domain, although Lys/Arg residues are usually located at the surface of proteins. Buried or partially buried Lys/Arg residues make salt bridges at both of the proposed Ca^2+^ binding sites mentioned earlier in the text: Lys276 and Lys 320 bind to Glu96 in the negative cluster at the tip of loop L5, Arg251 binds to Glu93 and Asp103 at either end of the L5 loop, and Lys265 and Lys268 bind to Asp103, 117, 118 and 121 at the site where Sm^3+^ binds in the TbMCA-Ib structure. Binding of Ca2^+^ at these sites would disrupt the salt bridges and thus destabilise the structure of the UNK domain, which may play a key role in the calcium activation mechanism. In the UNK of AtMCA-IIf on the other hand, there are fewer Lys/Arg, none of which are buried, few salt bridges, and only one of these is similarly located (Lys220 to Asp111 at the base of the L5 loop), which may further help explain why this enzyme is Ca2^+^ independent. However, future studies are needed to understand if, and in that case how, a low pH destabilizes the UNK domain of AtMCA-IIf and strong binding of the AIM at the catalytic center.

### A new group of small molecules for in vivo inhibition of metacaspase activity

To overcome functional redundancy and to develop a tunable inhibition strategy, we performed a target based chemical screen for small molecules that inhibit type II metacaspases. Small molecule inhibitors are often used as a means to inhibit therapeutic targets in a clinical setting, including proteases (Wu *et al*, 2022), but were also successfully applied to unravel biological mechanisms in plants, such as perception of the plant hormones ABA and brassinosteroid, membrane trafficking and primary metabolism (De Rybel *et al*, 2009; Kim *et al*, 2011; Kerchev *et al*, 2015; Dejonghe & Russinova, 2017; Rohzon *et al*, 2019; Vaidya *et al*, 2019, 2021). In contrast, few examples exist in the literature of small molecules that modulate plant protease activity in vivo. Mainly those can be found that are substrate mimetics: a peptide based on the substrate cleavage signature and modified with a ‘warhead’ that covalently binds and inactivates the protease active site during catalysis (Morimoto & Van Der Hoorn, 2016). For metacaspases, the most used substrate mimetic inhibitor is z-VRPR-fmk, which was derived from a preferred tetrapeptide substrate as determined by scanning of a combinatorial tetrapeptide library for AtMCA-IIf activity (Vercammen *et al*, 2006) and has a fluoromethyl ketone (fmk) warhead. Not only z-VRPR-fmk is used in enzymatic and in vivo assays in plants (Hander *et al*, 2019; Berenguer *et al*, 2021; Graff van Creveld *et al*, 2021; preprint: Cornblatt *et al*, 2021), but also in vivo against human paracaspase MALT1. However, in human studies z-VRPR-fmk suffers from poor cell permeability, probably due to the two Arg residues (Fontán *et al*, 2018), and biological toxicity (Demeyer *et al*, 2016; Hamp *et al*, 2021). More recently, new activity based probes with an acyloxymethylketone (AOMK) warhead were developed based on the AtMCA-IIf tetrapeptide library study and COFRADIC analysis (Vercammen *et al*, 2006; Tsiatsiani *et al*, 2013), but in vivo efficacy was not tested (Štrancar *et al*, 2022). Substrate mimetics and derivatives were also designed for inhibition of protozoan metacaspases (Berg *et al*, 2010). Few studies of small molecule inhibitors in *Trypanosoma* sp. and *Plasmodium* sp. are available, and even fewer that are not peptide or substrate-like, that would contain more diversity in their chemical structure. Nonetheless, through rational design based on the observation that z-FA-fmk can inhibit *Plasmodium falciparum* metacaspase-2 (PfMCA-2), Vandana *et al*. (2020) synthesized a non-peptidyl molecule named SS-5 that induces cell death in *P. falciparum*. More recently, two small molecules derived from SS-5, called C532 and C533, were found to effectively block transmission of *P. falciparum* and *P. berghei*, reducing the parasite burden in a mosquito host (Kumari *et al*, 2022). Bear in mind that protozoa exclusively have type I metacaspases (Tsiatsiani *et al*, 2011). In this study we screened against 10,000 chemicals thereby enlarging the chemical space of potential type II metacaspase inhibitors. The uncovered small molecule inhibitors in this study and their derivatives could prove to be active against medical important type I metacaspase in the future. However, given that highly similar TDP-containing small molecules can have very different effects in vivo (for instance, on LRP emergence), it is hard to predict which small molecules will be effective without further detailed studies in vivo.

### MCA-dependent substrate cleavages as a potential driver of cell separation and heightened immune response in lateral root emergence

Lateral root emergence requires an interplay between the emerging LRP and the overlaying tissues, including endodermal cells (Banda *et al*, 2019; Sager *et al*, 2021). Auxin signaling, mechanical feedback, regulated cell death, cytoskeleton dynamics and symplastic isolation are involved in this tightly regulated process of new organ development (Richter *et al*, 2009; Marhavý *et al*, 2016; Vilches Barro *et al*, 2019; Escamez *et al*, 2020; Sager *et al*, 2020). Genes expressed specifically in the endodermal cells adjacent to LRPs, such as *MYB36* and *PLDP5* (Fernaández-Marcos *et al*, 2017; Sager *et al*, 2020), can control the process of lateral root emergence. Here we demonstrated that metacaspases with a defined spatiotemporal endodermal expression pattern are necessary for a proficient outgrowth of LRPs. Endodermal cells are differentiated cells containing two impermeable polymers in their cell walls: lignin and suberin. Both the lignified Casparian strips and suberized lamellae are critical to form a tightly controlled bidirectional barrier between the environment (e.g. soil nutrients, water, microbiota) and the plant vasculature (Doblas *et al*, 2017).

Cell separation is required to allow for the developing LRP to protrude through the endodermal cell layer, while at the same time, remodelling of the lignin and suberin barriers is required to regulate nutrient and water flow, and likely to prevent colonization of the root interior by pathogens (Vermeer *et al*, 2014; Zhou *et al*, 2020; Ursache *et al*, 2021). The substrate specificity studies with a COFRADIC analysis on *atmca-IIf* single mutants in whole Arabidopsis roots revealed potential substrate proteins of AtMCA-IIf related to cell wall remodelling. Considering the extracellular localization of AtMCA-IIf (Vercammen *et al*, 2006), a subset of *AtMCA-IIf* co-expressed potential substrates (Appendix Table S2.1) that localize in the extracellular space, such as several glycosyl hydrolases and peroxidases, could be of key importance in cell wall remodelling. Glycosyl hydrolases (GHs) are encoded by multiple gene families in Arabidopsis and hydrolyze glycosidic bonds between two or more carbohydrates. AtMCA-IIf substrates of the GH family 1 (AT3G09260), 32 (AT1G12240), 38 (AT3G26720) have not yet been correlated with cell wall remodelling. However, since cell walls consist of 90% carbohydrates and only for 10% of proteins (Wojtaszek, 1997), modulation of GH by AtMCA-IIf might have an impact on cell wall structure. For example, N-glycan processing of glycosylated proteins by the class I α-mannosidase MNS2 (AT3G21160; GH family 47), together with its close homologs MNS1 and MNS3 is important for root development and cell wall biosynthesis (Liebminger *et al*, 2009). Interestingly, a *mns1mns2* double mutant displayed increased lateral root formation, whereas the *mns1mns2mns3* triple mutant had a short radially swollen root and showed defects in pectin accumulation (Passardi *et al*, 2004). Three different peroxidases (AT4G30170, AT1G05240, AT5G17820) were identified exclusively in the AtMCA-IIf-KO proteome, possibly as a result of their degradation in the WT proteome by AtMCA-IIf. Secreted class III peroxidases regulate the level of apoplastic H2O2 which is implicated in cell wall loosening and cross-linking to phenolic compounds (Lee *et al*, 2013). The precise subcellular distribution of a peroxidase (PER64) and reactive oxygen species (ROS) formation was linked to lignin crosslinking and casparian strip formation along the longitudinal plane of the endodermis (Bartels *et al*, 2013). Although not detected in the current COFRADIC analyses, PER64 might be degraded by AtMCA-IIf at the sites of lateral root development. Finally, UDP-D-apiose/UDP-D-xylose synthase 2 (AT1G08200), a protein whose depletion leads to cell wall thickening and cell death (Ahn et al, 2006), underwent a 17-fold higher processing in the AtMC9-OE plants compared to the AtMC9-KO. Together, this suggests the possibility of metacaspase-dependent cell wall remodelling to allow cell separation of the endodermal cells around the developing lateral root. Alternatively, AtMCA-IIf could be important for execution of cell death or clearance of dead cells (Bollhöner *et al*, 2012), as cell death of the endodermal cells overlaying the LRP was found to be important for lateral root emergence (Escamez *et al*, 2020). As not all proximal endodermal cells die during passage of the developing lateral root (Escamez *et al*, 2020), it is possible that a mixture of cell death and cell wall remodelling of endodermal cells takes place during LRP development. Furthermore, cell wall remodelling and cell death are not necessarily mutually exclusive events, as AtMCA-IIf can be active post-mortem for example in the clearance of cellular remnants in ‘dead’ xylem cells (Bollhöner *et al*, 2013). In the future, a detailed spatiotemporal analysis of cell wall remodelling-mediated cell separation and PCD in the root endodermis should facilitate our mechanistic comprehension of the role of AtMCA-IIf in lateral root emergence.

Besides their potential roles in cell wall remodelling, we can also speculate on a defensive role for metacaspases during LRP emergence. Physical damage-induced or temporary developmental breaches of root tissue integrity might create the opportunity for soil pathogens to invade the host plant. Zhou *et al*. noted that cortical cells in the vicinity of LRPs have a heightened responsiveness to bacterial elicitors such as flg22 (Zhou *et al*, 2020). We and others have previously shown that upon wounding of plants the calcium-dependent metacaspase AtMCA-IIa releases the immunomodulatory peptide Pep1 from its precursor PROPEP1 (Hander *et al*, 2019; Shen *et al*, 2019). PROPEPs are a family of precursor proteins from which small signaling peptides, Peps, are derived and typically are perceived by the leucine-rich repeat receptor-like kinases (LRR-RLKs), PEP RECEPTOR 1 and 2 (PEPR1 and PEPR2), and its co-receptor BRI1 ASSOCIATED RECEPTOR KINASE 1 (BAK1) (Huffaker *et al*, 2006; Krol *et al*, 2010; Yamaguchi *et al*, 2006, 2010; Bartels *et al*, 2013). Recently, Wang and Xi *et al*. found that Pep7 can be perceived by another LRR-RLK, SUCROSE-INDUCED RECEPTOR KINASE 1 (SIRK1), and its co-receptor QIAN SHOU KINASE 1 (QSK1) (Wang *et al*, 2022). Strikingly, the *sirk1* and *propep7* mutants have a delayed LRP emergence, whereas application of synthetic Pep7 peptide speeds up LRP emergence. The mature Pep7 peptide was identified previously from root tip protein extracts through COFRADIC analysis (preprint: Liu *et al*, 2017) and PROPEP7 can be cleaved by AtMCA-IIa in vitro (Hander *et al*, 2019) and in Arabidopsis protoplasts (Shen *et al*, 2019). Therefore, it is possible that AtMCA-IIf and potentially Ca^2+^-dependent type II MCAs exert their effect on LRP emergence through cleavage and release of Pep7. Nevertheless, the molecular mechanism behind the effect of Pep7 on LRP emergence remains unclear and metacaspases likely mediate LRP emergence through cleavage of multiple different substrate proteins. Certainly, further studies are required to tease apart the potential developmental and immune-related effects of metacaspase substrate cleavage in LRP emergence.

## Conclusion

In summary, we report here on the structure of the calcium-independent metacaspase AtMCA-IIf and provide a rationale for the inhibitory action of a class of newly discovered TDP-containing small molecules against metacaspases. Importantly, some of the TDP-containing small molecules (such as TDP6) have a specific effect in plants on LRP emergence where, due to the specific expression pattern of AtMCA-IIf and AtMCA-IIb in the endodermal cells overlaying the LRP, metacaspases can control LRP emergence. Furthermore, the structure of AtMCA-IIf contributes to a better understanding of the activation mechanism of calcium-independent metacaspases and other clan CD proteases, including paracaspases, caspases, legumain and separase. The TDP-containing small molecule inhibitors provide an extra tool to study metacaspase function in diverse plant species and can initiate the development of new drugs against neglected diseases caused by *Trypanosoma* and *Plasmodium* where metacaspases – absent in humans – are considered as suitable therapeutic targets.

## Materials and methods

### Plant lines

T-DNA insertion lines of AtMCA-IIb (SAIL-284-C06) and AtMCA-IIf (GK-540H06) were crossed. The AtMCA-IIf KO line contains a T-DNA insert at cDNA base pair position 483 downstream of the start codon, present as an inverted repeat (LB-RB|RB-LB). The AtMCA-IIb KO line contains a T-DNA insert at cDNA base pair position 1111 downstream of the start codon (Appendix Fig. S3a). Homozygous F2 plants were selected by segregation analysis and T-DNA insertions were PCR-validated with gene specific and T-DNA insert specific primers (Appendix Table S6). Subsequently, absence of AtMC5 and AtMC9 transcripts from the single and double KO lines was determined by RT-PCR with primers that amplify a 978 bp cDNA fragment of AtMCA-IIf and a 402 bp AtMCA-IIb fragment (Appendix Fig. S3b).

Generation of multiple type-II metacaspase silenced lines was achieved by means of the artificial microRNA (amiRNA) technology (Schwab *et al*, 2006). The amiRNA construct hybridizes on a common sequence segment in *AtMCA-IIa, b, c, d* and *f*. Since *AtMCA-IIe* was not targeted, the construct was introduced into an *AtMCA-IIe* KO line (JII Gene Trap line GT_3_12679) of Landsberg erecta background (He *et al*, 2008) (Appendix Fig. S10a). The amiRNA construct was engineered with 4 oligonucleotide sequences (Appendix Table S6) and two modified Gateway^TM^ compatible A and B oligonucleuotides for recombination to the entry vector pDONR221. Subsequently, the construct was fused to the CAMV 35S promoter by LR reaction to the destination vector pB7WG2 (Karimi *et al*, 2002). The p35s:amiRNA fusion construct was transformed into the *Agrobacterium tumefaciens* strain C58C1RifR[pMP90]. Transgenic *Arabidopsis thaliana* (L.) Heynh., accessdion Landsberg *erecta* (L*er*) plants were obtained via floral dip transformation (Clough & Bent, 1998) and subsequent Basta selection. Independent T3 homozygous lines containing a single locus were selected for further analysis.

Transcriptional and translational reporter lines were made as follows. The promoter region of metacaspase genes was cloned into the binary vector pBGWFS7 (Karimi *et al*, 2002) giving rise to pAtMCA:GFP:GUS fusion reporter lines. The genomic region used to study promoter activity started at 1500 bp and ended 1 bp upstream of the start codon of each metacaspase gene. Forward and reverse PCR primers used for amplification of each promoter region are listed in Appendix Table S6. Extensions to complete Gateway^TM^ attB1 and attB2 cloning sites were obtained by a second PCR reaction. Constructs were introduced into *Arabidopsis thaliana*, accession Columbia-0 (Col-0) as previously described by *A. tumefasciens* mediated floral dip transformation.

### Tissue expression analysis

GUS staining of 7-day-old plants started with a 10 min incubation in 90% (v/v) acetone at room temperature. After washing in phosphate buffer pH 7.4, the material was incubated in 1 mg/ml 5-bromo-4-chloro-3-indolyl-D-glucuronide, 2 mM ferricyanide, and 0.5 mM ferrocyanide in 100 mM phosphate buffer, pH 7.4 at 37°C in the dark for 4 h to overnight. The material was cleared with 85% lactate solution and observed by light microscopy. Fluorescence microscopy of the pAtMCA:GFP:GUS fusion reporter lines was performed with a confocal microscope 100M and software package LSM 510 version 3.2 (Zeiss, Jena, Germany). Excitation was performed with a 488 nm argon laser. Emission fluorescence for GFP was captured via a 500-550 nm band-pass filter, and for propidium iodide (PI) via a 585 nm long-pass filter. Fluorescence microscopy of the pAtMCA-IIf:AtMCA-IIf-GFP translational fusion lines was performed on a Zeiss LSM 710 confocal scanning microscope, excitation with a 488 nm argon laser and appropriate emission windows for GFP and PI stain.

### Preparation and purification of native recombinant metacaspases

Recombinant AtMCA-IIa and AtMCA-IIf were produced as previously detailed in Vercammen et al., 2004 (Vercammen *et al*, 2004). An *AtMCA-IIb* CDS was cloned to a Gateway™ pDEST™17 vector and transformed into the *E. coli* strain BL21 (DE3). Protein expression was induced for 3 h with 0.2 mM isopropyl 1-thio-D-galactopyranoside at 37°C. Cultures were harvested, re-suspended in extraction buffer (50 mM HEPES, pH 7.5, 300 mM NaCl, 8 M urea) and sonicated. Cellular debris was removed by centrifugation, and the supernatant was mixed with TALON cobalt affinity resin (BD Biosciences) and incubated overnight at 4°C. Beads were washed eight times in the same buffer, while stepwise reducing the urea concentration from 8 to 0 M, loaded on a column, and washed twice again (50 mM HEPES, pH 7.5, 300 mM NaCl). Bound protein was eluted with extraction buffer supplemented with 150 mM imidazole.

*Escherichia coli* BL21(DE3)pLysE bacteria transformed with the pDEST17-AtMCA-IIf C147A expression plasmid were grown in shake flask cultures at 200 rpm and 37°C in an auto-induction medium containing 34 µg/ml chloramphenicol and 100 µg/ml ampicillin (Studier, 2005). After reaching OD_600_ value between 0.6 and 0.8, the incubation temperature was lowered to 16°C and the incubation was continued overnight. Cultures were harvested by centrifugation at 5000*g* for 20 min and the cell pellets were stored at −20°C until purification.

All following steps were performed at 4°C. Cell pellet was resuspended using a ratio of 1 g to 5 ml of binding buffer (20 mM Tris, 500 mM NaCl, 20 mM imidazole, pH 7.4). Resuspended cells were lysed using a cell disruptor at 20,000 psi and treated with DNaze I for 30 min. Insoluble cell debris was removed by centrifugation at 38,000*g* for 20 min and the supernatant was filtered through a 0.45 µm filter. Filtered supernatant was then applied to a 5 ml HiTrap Chelating HP column (Cytiva), Ni^2+^-loaded according to manufacturer’s instructions, connected to an ÄKTA FPLC and equilibrated with binding buffer. The affinity column was washed with approximately 10 column volumes of the binding buffer after which the protein was eluted using 75 ml linear gradient of 20-300 mM imidazole in the same buffer. Purity of the obtained protein fractions was visualised on 4-20% gradient SDS-PAGE gels, and the purest fractions were merged and concentrated to 2 ml using centrifugal concentrators (Vivaspin 10K, Cytiva). The protein sample was then purified using size exclusion chromatography on a HiLoad 16/600 Superdex 200 pg column (GE Healthcare), previously equilibrated with buffer containing 20 mM Tris-HCl, 100 mM NaCl, pH 7.4. Protein fractions were analysed with SDS-PAGE, pooled and concentrated to 14.2 mg protein/ml using centrifugal concentrator. Enzyme concentration was calculated from measured *A*_280_ nm on a NanoDrop TM 1000 Spectrophotometer (Thermo Scientific), and theoretical molar extinction coefficient, 12295 M^-1^ cm^-1^, determined by ProtParam tool on the ExPASy server (Wilkins *et al*, 1999).

### Crystallisation, data collection, and structure determination of inactive AtMCA-IIf C147A

Crystallisation screening was performed using Mosquito-LCP robot (SPT Labtech) in three-drop 96-well Protein Crystallography Plate (Corning). Crystals were grown in a RockImager 1000 system (Formulatrix) at 4 and 20°C. Four commercial screens were used: JCSG+ (Qiagen), Index, Crystal Screen HT, and PEGSuite (Hampton research). Three drops were set using 9.0 mg/ml protein solution (in 20 mM Tris-HCl, 100 mM NaCl, pH 7.4), with each crystallisation condition in the ratios: 150 + 150, 100 + 200, and 200 + 100 nl. The first crystals were obtained in two different conditions, both in second drop at 4°C: F6 Crystal Screen HT (0.01 M iron(III) chloride hexahydrate, 0.1 M sodium citrate tribasic dihydrate pH 5.6, 10% v/v Jeffamine M-600), and C3 JCSG+ (0.2 M ammonium nitrate, 20% w/v PEG 3350). The best diffracting crystal was obtained from hanging drop method by mixing 1 µl of 14.2 mg/ml protein solution and 1 µl of C3 JCSG+ condition (prepared in the laboratory) set up and incubated for 3 months at 20°C in EasyXtal 15-Well Tool X-Seal plate. Before flash-cooling in liquid nitrogen, the crystals were soaked in a cryo-protectant solution prepared by mixing 100 µl of C3 JCSG+ and 150 µl of 50% PEG 3350. Automated X-ray diffraction data collection was carried out at Diamond Light Source (Didcot, UK), on beamline i03 at 0.9763 Å wavelength. A total of 3600 images were collected, covering 360°, at 2.00 Å resolution. Statistics from diffraction data processing and structure refinement are summarized in Appendix Table S6.

Data processing was performed using XDS (Kabsch, 2010), and data scaling using Aimless (Evans & Murshudov, 2013) of the CCP4 software suite (Winn *et al*, 2011). To calculate R_free_, 5% of the reflections were randomly selected within the Aimless program and excluded from all refinements. The structure was solved by molecular replacement using a predicted structure of AtMCA-IIf as search model (obtained from AlphaFold [Jumper *et al*, 2021; Varadi *et al*, 2022], and the Phaser program [McCoy *et al*, 2007]) of the Phenix software package (Liebschner *et al*, 2019). The structure model was further improved by alternate use of COOT (Emsley *et al*, 2010), phenix.refine (Afonine *et al*, 2012), and PDBredo (Joosten *et al*, 2014). COOT was used for manually building, fitting and real space refinement of individual residues against σ_A_-weighted 2*F*_o_-*F*_c_ and *F*_o_-*F*_c_ electron density maps, while structure refinement was done in phenix.refine and PDBredo. Translation, rotation, and screw-rotation (TLS) parameterization of anisotropic displacement was used in the last few refinement steps. Coordinates and structure factors have been deposited in the Protein Data Bank under accession number 8A53. Protein structure figures were produced using PyMOL (DeLano, 2002).

### Plant material and total proteome extraction for COFRADIC analysis

Seeds of Arabidopsis Columbia-0 (Col-0), AtMCA-IIf T-DNA insertion line (GK-506H04-019739) and AtMCA-IIf overexpression (Vercammen et al., 2006) were overnight gas sterilized with HCl and NaOCl, then sowed on half strength Murashige and Skoog (MS) media and stratified at 4°C in the dark. After 3 days, seeds were transferred to 21°C with a 16 h light/8 h dark photoperiod with light intensity of 80-100 µmol m^-2^ s^-1^ and left to grow for 3 weeks. Roots were harvested, frozen in liquid nitrogen and ground into a fine powder using a mortar and a pestle. To achieve a total protein content of approximately 4 mg, 0.5 g of ground tissue was defrosted in 1 ml buffer of 1% (w/v) CHAPS, 0.5% (w/v) deoxycholate, 0.1% (w/v) SDS, 5 mM EDTA and 10% glycerol in PBS at pH 7.5, further containing the suggested amount of protease inhibitors mixture according to the manufacturer’s instructions (Roche). The sample was centrifuged at 16,000 g for 10 min at 4°C and guanidinium hydrochloride was added to the cleared supernatant to reach a final concentration of 4 M.

For the in vitro COFRADIC analysis, protein extracts were prepared in the AtMCA-IIf optimal activity buffer; 50 mM MES (pH 5.5), 150 mM NaCl, 10% (w/v) sucrose, 0.1% (w/v) CHAPS and 10 mM DTT. Four mg of total root protein extract was incubated at 30°C with recombinant AtMCA-IIf for a total duration of one hour; here, every 20 min 10 µg of fresh protease (855 units) was pre-activated in the same optimal activity buffer at room temperature for 15 min and then added to the protein extract (enzyme activity is 57 U/μl and one unit corresponds to the enzyme activity that catalyzes the hydrolysis of 1 µmol VRPR-AMC per min at 30°C). This was repeated twice. The same procedure was used for the control sample, but without adding AtMCA-IIf. Reactions were terminated by adding guanidinium hydrochloride to a final concentration of 4 M. Protein concentration measurements were performed using the DC protein assay (BioRad).

### N-terminal COFRADIC, peptide identification and quantification

Protein extracts were modified for N-terminal COFRADIC analysis as described in Staes et al. (2011), with few modifications. Briefly, in all three analyses, α and ε amines in the KO proteomes were 13C4-butyrylated, while amines of WT, OE or AtMCA-IIf treated-KO proteomes were 12C4-butyrylated. MS/MS-spectra were searched using the MASCOT algorithm against the TAIR protein database release 10. The following search parameters were set. Mass tolerance on peptide precursor ions was set to 10 ppm (with MASCOT C13 option set to 1) and that of fragment ions was set to 0.5 Da. Allowed peptide charges were 1+, 2+, 3+ and instrument setting was put on ESI-TRAP. Endoproteinase semi-Arg C/P was used, allowing for 1 missed cleavage. Two types of MASCOT searches were performed that differed in the set of variable and fixed modifications. The first type of searches included the variable modifications of pyro-glutamate formation of N-terminal glutamine and the fixed modifications of methionine oxidation to its sulfoxide derivative, S-carbamidomethylation of cysteine and butyrylation (12C4 or 13C4) of lysine and peptide N-termini. The second type of searches differed in that acetylation of peptide N-termini was set as a variable modification. Only peptides with a score higher than the MASCOT identity threshold score set at 99% confidence were withheld. Finally, Mascot Distiller (version 2.4) was used for peptide quantification.

### Screening of AtMC9 inhibitors

High-throughput screens of 10,000 compounds comprising the DIVERSet™ compound library of Chembridge were carried out in 384-well plates (Genetix X7001) and the reactions were performed in 50 µl final volume. 5 nM recombinant AtMCA-IIf, pre-activated for 15 min in 50 mM MES pH 5.5, 150 mM NaCl, 10% sucrose, 0.1% CHAPS and 1 mM DTT, was added to 10 µM of each compound in 10% DMSO. The compounds and AtMCA-IIf were co-incubated for 15 min at room temperature and then 10 µM Ac-VRPR-AMC was added to the assay mixture. As negative control, 10% DMSO was used, while 2.3 pmol of Ac-VRPR-FMK was used a positive control (90% inhibition). The fluorescence units measured for 10% DMSO are considered as 100% activity. Time-dependent release of free amido-4-methylcoumarin (AMC) was measured on a FLUOstar OPTIMA reader (BMG Labtechnologies, Offenburg, Germany) for 45 cycles of 1 min. Activity was expressed as the rate of fluorescence release per minute in each well and the assay was performed twice. A third screen was done in small scale (96-well plates) and involved 177 selected compounds with at least 75% inhibitory effect towards AtMCA-IIf in both previous screens. During the small-scale screen, besides AtMCA-IIf, the compounds were tested against AtMCA-IIa and AtMCA-IIb activities. These enzymes were used in concentrations that provide a signal similar in magnitude as AtMCA-IIf. For the AtMCA-IIa assay, 17 nM enzyme was pre-activated at room temperature for 15 min in 50 mM HEPES (pH 7.5), 150 mM NaCl, 10% glycerol, 100 mM CaCl_2_ and 10 mM DTT, while for the AtMCA-IIb assay, 72.5 nM enzyme was pre-activated at room temperature for 15 min in 10 mM HEPES (pH 7.5), 10 mM CaCl_2_ and 10 mM DTT. In each case, 10 µΜ compound was co-incubated with the enzymes for 15 min at room temperature and then 10 µM of the equivalent AMC substrate was added to the assay mixture.

### In silico analyses

IceLogo (Colaert *et al*, 2009) analyses were performed via the web application (http://iomics.ugent.be/icelogoserver/logo.html) using the Arabidopsis Swiss-Prot proteome as reference. Hierarchical clustering of the small-molecule metacaspase inhibitors was done via the open access software ChemMine (http://chemmine.ucr.edu/) based on structural and physicochemical similarities of the compounds using the Tanimoto similarity coefficient.

### In vitro HIT verification and full dose-response validation

The enzymatic activity of recombinant AtMCA-IIa, AtMCA-IIb and, AtMCA-IIf in the presence of the TDP substances were determined in vitro by following the cleavage of the fluorogenic metacaspase substrate Val-Arg-Pro-Arg-7-amino-4-methyl Coumarin (VRPR-AMC). The kinetic measurements were performed on a BioTek Synergy^TM^ H4 hybrid microplate reader, monitoring the change in fluorescence light at Ex355/Em460 nm. An isothiouronium salt, purchased from Sigma Aldrich, was used as a positive control towards AtMCA-IIa, AtMCA-IIb and, AtMCA-IIf (Appendix Fig. S5b).

Assay measurements on AtMCA-IIa and AtMCA-IIb were performed in HEPES buffer (0.2 M Hepes, 0.6 M NaCl, 40% Glycerol, 0.04% Fishgelatin, 0.02% Tween 20, 0.004 M dithiothreitol at pH 7.5). The assay reaction mixture contained 0.01 µM of AtMCA-IIa/AtMCA-IIb and the reaction was initiated by the addition of 50 mM of CaCl_2_ and 50 µM of VRPR-AMC after 10 minutes of pre-incubation. Measurements on AtMCA-IIf were performed in MES buffer (0.2 M MES, 0.6 M NaCl, 40% Glycerol, 0.04% Fish gelatin, 0.02% Tween 20, 0.004 M dithiothreitol at pH 5.5). The final assay concentration of AtMCA-IIf was 0.08 µM. The reaction was initiated by the addition of 50 µM of VRPR-AMC to each well. Inhibition curves for TDP1-TDP7 were measured for 12 concentrations between 0.3-250 µM for triplicate samples. The reaction was monitored for 10 minutes and the mean velocity (RFU/sec) was calculated within the linear range of the reaction (AtMCA-IIa: 1-6 min, AtMCA-IIb: 1-3 min, AtMCA-IIf: 1-3 min). Activities were normalized over the blank mean for each triplicate well. IC50 values were determined from a four-variable non-linear regression fit in GraphPad Prism, using triplicate data for AtMCA-IIf, and duplicate data for AtMCA-IIa and AtMCA-IIb.

### Compound characterization by nuclear magnetic resonance

Selected TDP-containing compounds were re-ordered from Chembridge (TDP1 to TDP7; Appendix Table S1). ^1^H Nuclear magnetic resonance (NMR) spectra of TDP1 to TDP7 were recorded on a Bruker 400 MHz Avance III spectrometer at 400 MHz (^1^H) at 25°C. Chemical shifts are reported in ppm and δ values were referenced to the residual solvent signal of DMSO-*d_6_* (2.50 ppm) as an internal standard (Appendix Table S3).

### Inhibition of GST-TEV-PROPEP1 cleavage

An orthogonal in vitro assay based on the cleavage of a recombinant MCA substrate protein was developed as alternative for the VRPR-AMC based activity assay. GST-TEV-PROPEP1 is a fusion protein of Glutathione-S-transferase (GST) with a Tobacco Etch Virus (TEV) protease cleavage site and a N-terminal truncation product of AtPROPEP1. GST-TEV-PROPEP1 was obtained as previously published (Hander *et al*, 2019) and diluted in dH_2_O to a concentration of 0.5 mg/ml. Optimal assay conditions with regards to MCA concentration (final concentrations of 0.1 µg/ml AtMCA-IIf, 0.2 µg/ml AtMCA-IIb, and 0.2 µg/ml AtMCA-IIa) and reaction time (5 minutes) were obtained beforehand to ensure that the reaction occurs in the linear range and GST-TEV-PROPEP1 cleavage is not saturated. MCAs were diluted in dH_2_O to four times the final concentration. Four-fold reactions buffers were prepared for AtMCA-IIa/b (200 mM HEPES-KOH, pH7.5, 600 mM NaCl, 40% (v/v) glycerol, 40 mM DTT, and 200 mM CaCl_2_) and AtMCA-IIf (200 mM MES-KOH, pH5.5, 600 mM NaCl, 40% (w/v) sucrose, 0.4% CHAPS, 40 mM DTT). Dilution series of a 10 mM TDP6 stock solution (solved in DMSO) were made in dH_2_O starting with a four-fold concentration of 1 mM and further dilution steps in dH_2_O (final concentrations in the reaction of 250, 125, 62.5, 31.25, 15.6, 7.8, and 3.9 µM). As empty control (0 µM TDP6), a volume of DMSO equivalent to the 1 mM TDP6 dilution was diluted in dH_2_O. Reactions were carried out in 8-well strips with 5 µl buffer, 5 µl MCA, 5 µl GST-TEV-PROPEP1, and 5 ul of the TDP6 dilutions at 30°C. AtMCA-IIf was pre-activated in buffer (1:1 of the four-fold concentrations) for 10 minutes at room temperature, after which it was added to a premix of GST-TEV-PROPEP1 and the TDP6 dilution series (1:1 of the four-fold concentrations) by multi-channel pipette. AtMCA-IIa/b were not pre-activated and added directly to a premix of buffer, GST-TEV-PROPEP1 and the TDP6 dilution series (1:1:1 of the four-fold concentrations) by multi-channel pipette. Reactions were stopped by the addition of 20 µl Laemmli SDS 2x buffer by multi-channel pipette and samples were heated for 10 minutes at 70°C. An unreacted control sample (C; Appendix Fig. S6) was prepared by mixing buffer, GST-TEV-PROPEP1 and dH2O (1:1:2) and heated for 10 minutes at 70°C. Samples were separated on 12% SDS-PAGE gels (Mini-Protean TGX Stain-Free Precast, BioRad) and GST-TEV-PROPEP1 protein bands were visualized on a BioRad Imager (stain-free). An equal amount of protein bands were manually appointed per lane and adjusted in the Lane profile menu of the Image Lab software (BioRad). Band intensity was quantified relative to the unreacted control sample.

### Molecular docking simulations

The X-ray crystal structure of *At*MCA-IIf was prepared for docking using the protein preparation wizard implemented in Maestro (Sastry *et al*, 2013). The autoinhibitory motif was removed from the active site (Glu174-Pro186), Ala147 was mutated to Cys147, water molecules were deleted, and termini were capped. This was followed by addition of hydrogens at pH 5.5 and optimization of the hydrogen bond network. The receptor grid was prepared based on the position of Arg183-Leu185 of the autoinhibitory motif, resulting in x-, y-, and z-coordinates of −16.5, −22.8, and −19.6. The inner and outer box dimensions were set to 10 Å and 33 Å, respectively. The scaling factor for the van der Waals radius was set to 0.9 to soften the potential for nonpolar parts of the receptor. The ligands were docked using Glide implemented in Maestro using standard precision mode (Friesner *et al*, 2004; Halgren *et al*, 2004). The Coulomb-vdW energy cutoff was increased to 1.0 kcal/mol and the RMS cutoff was decreased to 0 Å. The number of poses included during post-docking minimization was increased to 500, and the number of output poses was increased to 100, per ligand. Only poses where the ligand is placed in the S_1_ pocket, where Arg183 of the autoinhibitory motif is placed, were considered. Both the *E*- and *Z*-configurations of TDP2, TDP3 and TDP6 were included in the dockings. The Ser101-Lys109 loop was manually built with the Molecular Operating Environment (MOE) software (Chemical Computer Group ULC, Montreal, Canada), aided by the position of the corresponding loop of *At*MCA-IIa, followed by energy minimization of the added amino acid residues.

### RNA extraction, reverse-transcription PCR and real-time quantitative PCR

Total RNA was extracted from leaves as previously described (Bentsink *et al*, 2006). One µg of RNA was reverse-transcribed using the iScript™ cDNA Synthesis Kit (Bio-Rad). Real-time quantitative PCR was performed in a final volume of 5 µl including 10% (v/v) of 6-fold diluted cDNA as template for PCR, 200 nM of each primer and 50% (v/v) SYBR green mix (Invitrogen) using a LightCycler 480 II instrument (Roche Applied Science), according to the manufacturer’s instructions. The genes CBP20 (At5g44200) and ARP7 (At3g60830) were used as internal references for normalization of transcript levels. Metacaspase gene-specific primer sequences used for real-time quantitative PCR and reverse-transcription PCR are listed in Appendix Table S6.

### Plant treatments and growth conditions

Surface-sterilized seeds were grown on vertical plates containing half strength Murashige and Skoog (½MS) salts, 1% (w/v) sucrose, 0.8% (w/v) plant tissue culture agar (LabM, Bury, UK), 0.5 mg/l nicotinic acid, 0.5 mg/l pyridoxin and 1 mg/l thiamin. The seeds were stratified at 4°C for 2 nights in the dark and then left to germinate in continuous light (80-100 µmol m-2s-1) at 21°C. Three days after germination, seedlings were transferred to ½ MS agar plates containing 50 µM compounds and then 5 days later their root growth and phenotypical appearance was monitored. Staining of lateral root primordia and developmental staging was performed 10 days after germination as described (Malamy & Benfey, 1997).

For the TDP6 to DMSO transfer experiment, Arabidopsis (Col-0) seeds were sown on ½MS plates (same composition as above) in two parallel rows with 10 seeds per row. After stratification at 4°C for 3 days, the plates were placed vertical to allow the seedlings to germinate unperturbed in 16 h/8 h light/dark conditions at 21°C. After 3 days, the seedlings were transferred with sterile tweezers to ½MS plates containing 50 µM TDP6 and grown for 5 days in the presence of TDP6 (inhibitor treatment). Thereafter, the seedlings were transferred a second time with tweezers to a plate containing an equal volume of DMSO as control (relieve of inhibition), or were left to grow on the TDP6 containing plates, for another 6 days. Pictures of the plates were taken with an in-house fixed camera setup at 0 days after the second transfer (0 DAT) and 6 DAT. The position of the root tip at 0 DAT was indicated with a stripe, which was copied to the 6 DAT pictures, and the number of emerged lateral roots were counted above and below this stripe. Data visualization and statistics were performed with GraphPad Prism.

### Root bending assay and lateral root primordia staging

Surface-sterilized Arabidopsis seeds of Col-0, L*er*, and AtMCA-II amiRNA knockdown lines KD1 and KD1, were sown on ½ MS plates containing 1% (w/v) sucrose, 0.01% (w/v) myo-inositol and 0.05% MES-KOH, pH 5.7. The plates were stratified for 2 days and transferred to the growth chamber to grow vertically under continuous light at 22°C. After 48 hours, seedlings were transferred to ½ MS plates containing 25µM TDP6 or an equivalent volume of DMSO as control. After 24 hours, plates were rotated 90° for 48 hours. The seedlings were then fixed with FAA for 30 minutes and were rehydrated for 20 minutes each in 70%, 50%, 30%, and 10% ethanol and infiltrated overnight in 50% glycerol. The fixation and infiltration steps were taking place on ice and all solutions were kept cold. The seedlings were mounted in 50% glycerol under glass coverslips. Images of lateral root primordia were collected with an AxioCam MR3 (Carl Zeiss, Göttingen, Germany). The number and stage of lateral root primordia were registered according to Malamy & Benfey (1997).

### Data availability

All data porting this study are included in the main text and Appendix.

## Acknowledgements

Dr. Gerhard Hofer (Department of Materials and Environmental Chemistry, Stockholm University) is gratefully acknowledged for valuable advice and help to solve the phase problem in the initial steps of structure determination. We thank M. De Cock for help in preparing the manuscript. We acknowledge Diamond Light Source (UK) for time on Beamline i03 under Proposal MX23773. X-ray diffraction data processing computations were performed at NSC Tetralith/LUNARC Aurora provided by the Swedish National Infrastructure for Computing (SNIC) and PReSTO funded by the Swedish Research Council, grant 2018-05973 (SNIC) and 2018-06479 (PReSTO). This work was supported by the Research Foundation-Flanders (grant FWO14/PDO/166 to S.S.), the Knut and Alice Wallenberg Foundation (grant 2018.0026 to J.S., P.V.B., and A.L. and grant 2021.0071 to S.S.), and the Ghent University Special Research Fund (grant 01J11311 to F.V.B).

## Author contributions

All authors helped to conceive and design the analysis. S.S., I.S., D.A., T.A., L.T., R.P.K., A.V.-A., C.L., D.V., S.J., L.N., M.N., A.D.F.-F., T.B., and E.T. performed the experimental work. S.S., I.S., L.T., J.S., A.L., and F.V.B. wrote the manuscript. S.S., I.S., K.G., J.S., P.V.B, A.L., T.B. and F.V.B. revised the manuscript and were involved in the discussion of the work.

## Disclosure and computing interests statement

The authors declare that they have no conflict of interest.

---

**Supplementary Fig. 1:**
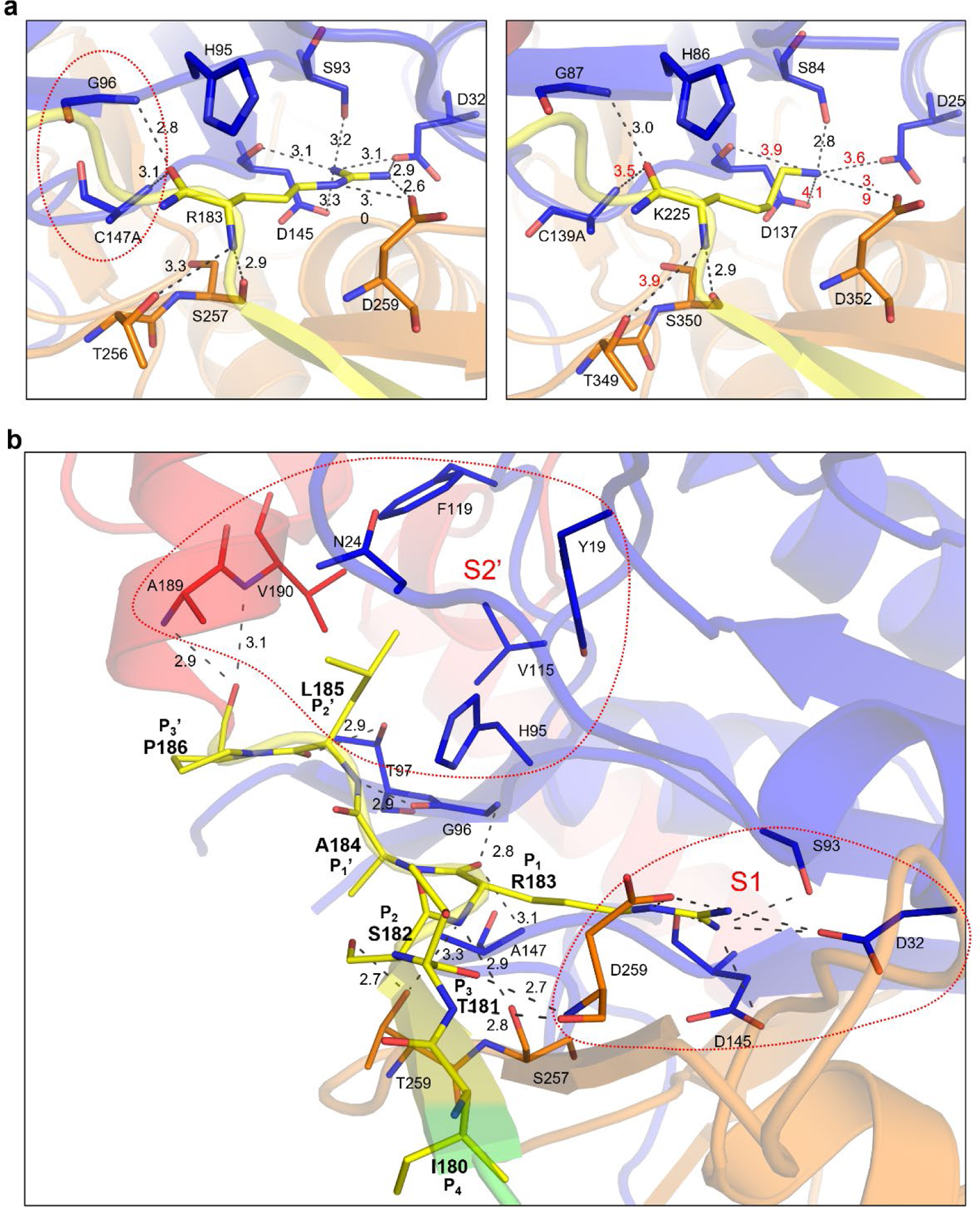
Close-up views of the catalytic centre and AIM (autoinhibitory motif) binding in the S1 and S2’ pockets. **a**, Polar interactions/hydrogen bonds between the P1 residue and protein at the catalytic centre and the S1 pocket, Arg183 in AtMCA-IIf (left panel) and Lys225 in AtMCA-IIa (right panel). Interaction distances in Å are shown by numbers, in red for weaker bonds (3.5 Å or longer). **b**, Binding of the AIM in the S1 and S2’ pockets in the AtMCA-IIf C147A structure. Different regions are color coded as: p20, blue; linker, green; AIM, yellow; UNK, red; p10, yellow. All residues in the AIM motif are shown, in stick representation, as well as selected and pocket-forming residues.

**Supplementary Fig. 2:**
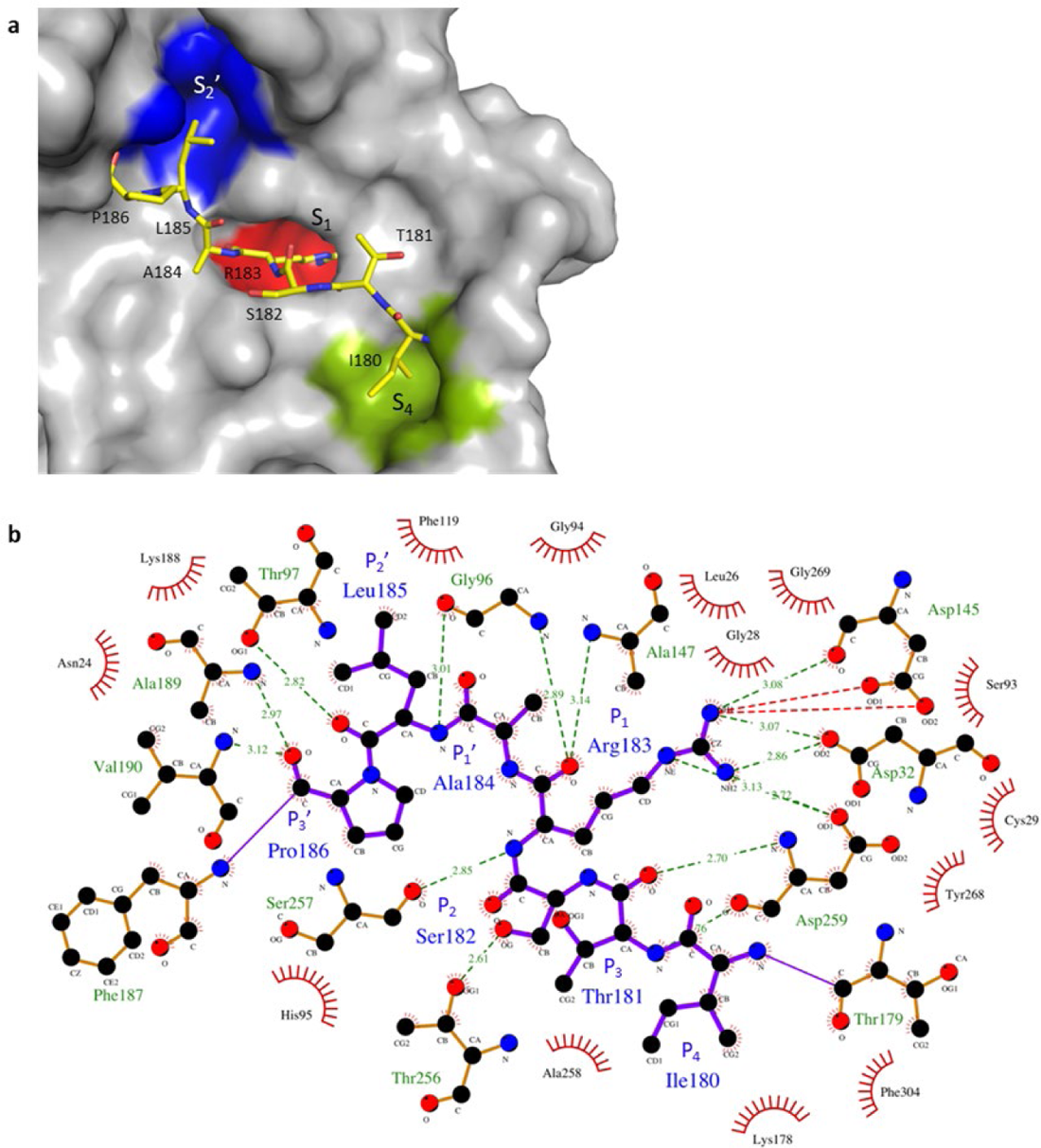
Binding of the AIM at the active site in the AtMCA-IIf C147A structure. **a**, Rendering of the water-accessible surface of the protein at the active site. The most clear sidechain-binding pockets S4, S1 and S2’ are shown in different colors and the AIM residues as sticks. **b**, LIG-plot of interactions between the AIM and surrounding protein residues [103].

**Supplementary Fig. 3:**
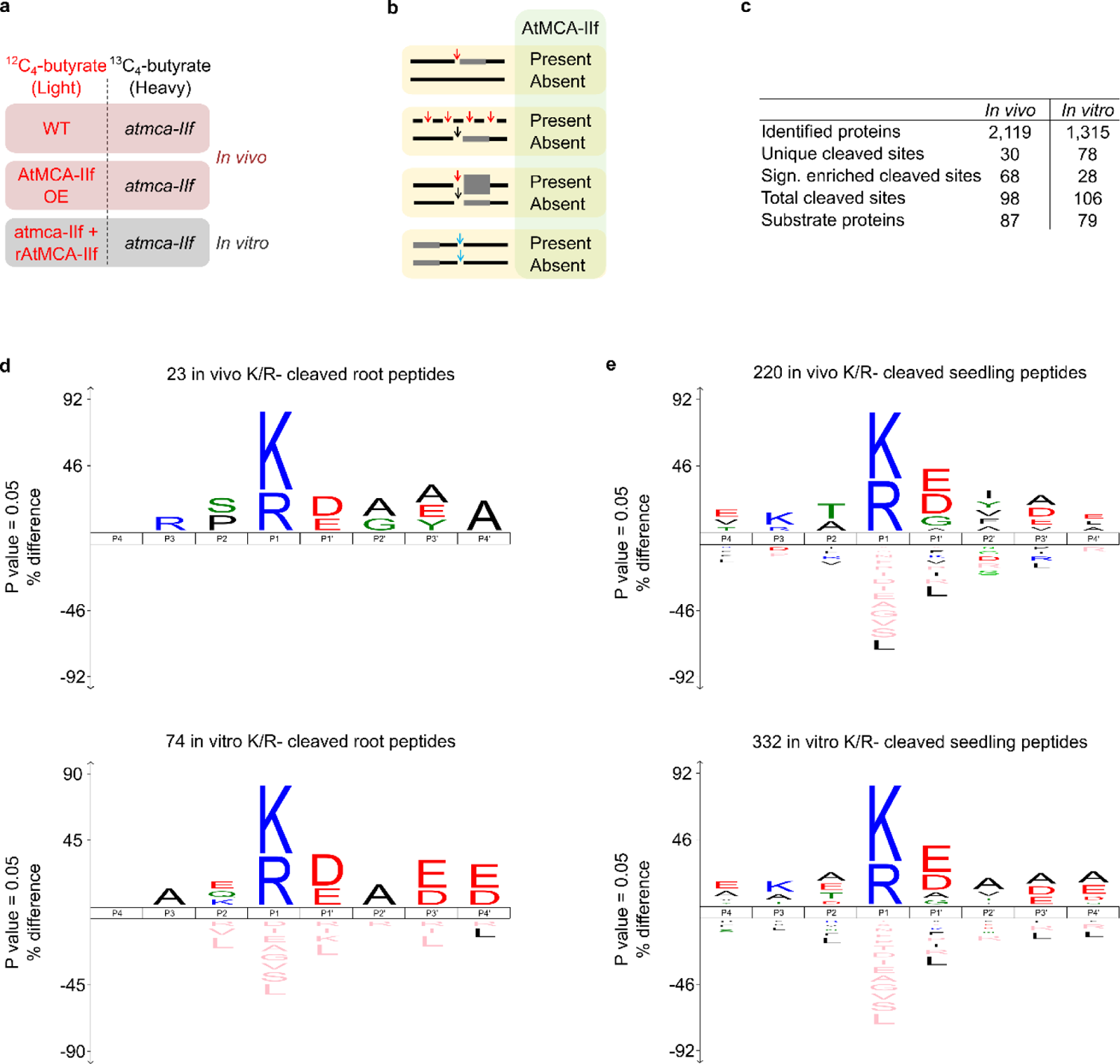
AtMCA-IIf in vitro and in vivo COFRADIC to identify substrate specificity and potential novel substrates. **a**, Experimental setup of the two in vivo and one in vitro proteomic analyses. The compared proteomes were differentially labelled with isotopic variants of NHS-esters of butyric acid (light, 12C4-butyrate; or heavy, 13C4-butyrate) and mixed in equal amounts. **b,** Possible proteolysis scenarios (left) and read-outs (right) in the presence or absence of AtMCA-IIf activity. From top to bottom: A protein is solely cleaved by AtMCA-IIf (red arrow) in the samples of active (WT or OE) or added (rAtMC9) protease, which generates a neo-N-terminal fragment. A protein is completely degraded by AtMC9 and no fragments are detected. A protein is cleaved in both samples but in a higher frequency in the sample with AtMC9, which generates a neo-N-terminal fragment in both samples but in a higher intensity in the protease sample. Proteolysis of the target in the sample without AtMC9 may have occurred by a protease with redundant activity to AtMC9 (black arrow). Fragments generated by trypsin (blue arrows) that bear the mature N-terminus of a protein are found in approximately equal intensities in both samples. **c,** Overview of the in vivo and in vitro cleaved sites and their converging proteins that were identified in this study. **d and e,** Sequence alignment (iceLogo [95]) of the in vitro (top) and in vivo K/R-cleaved proteins (bottom) found in this study (d) and in Tsiatstiani *et al*. [30] (e).

**Supplementary fig. 4:**
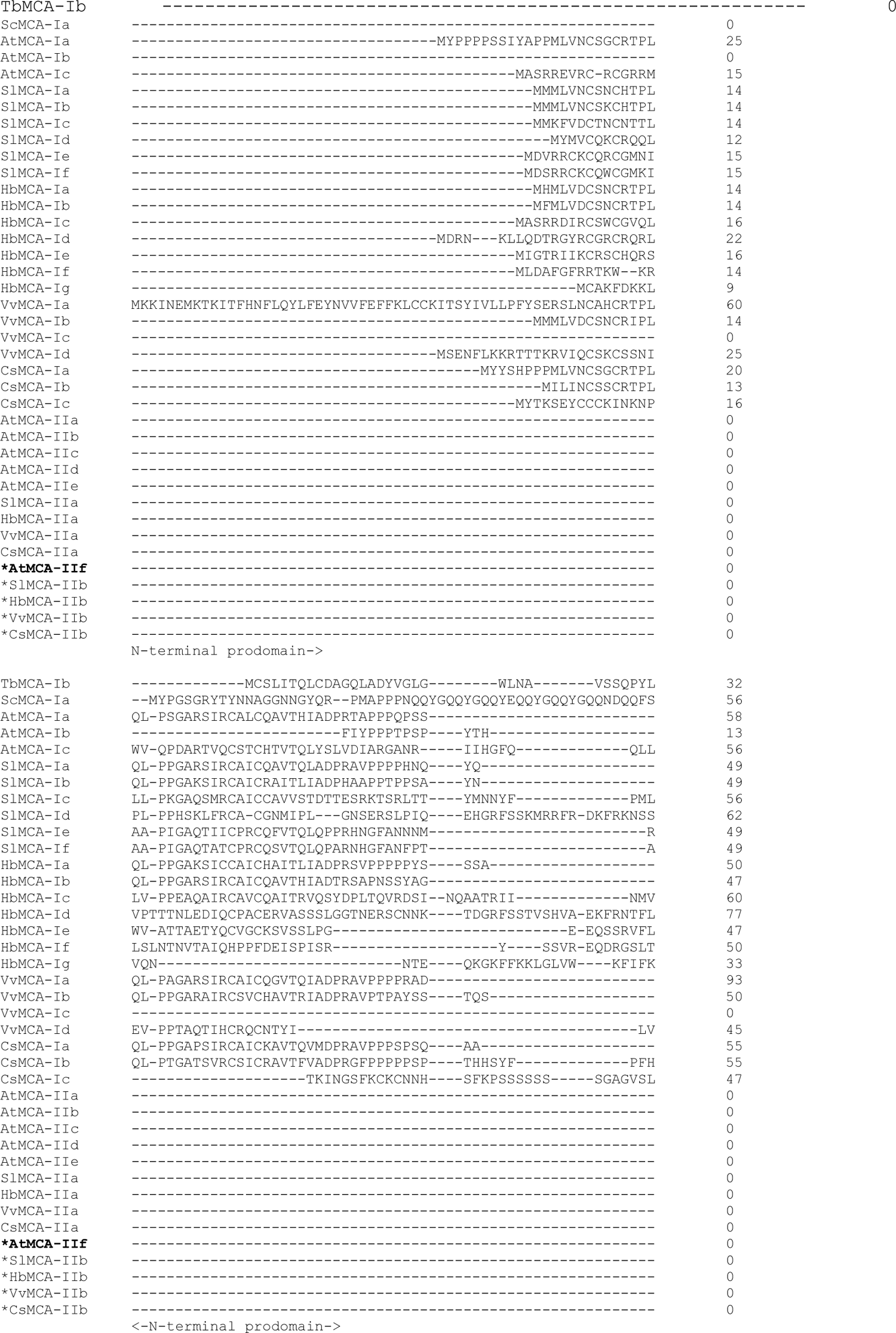
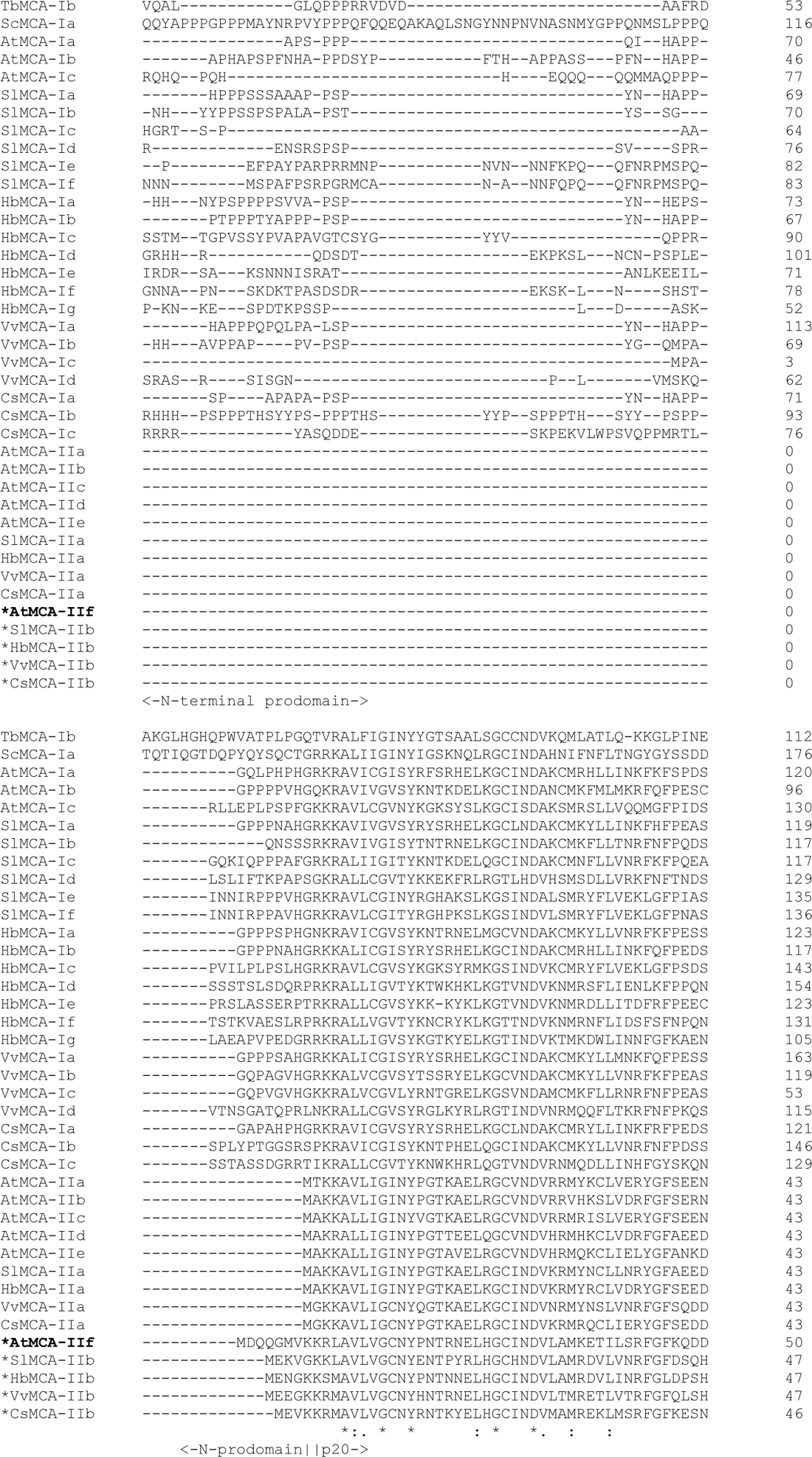
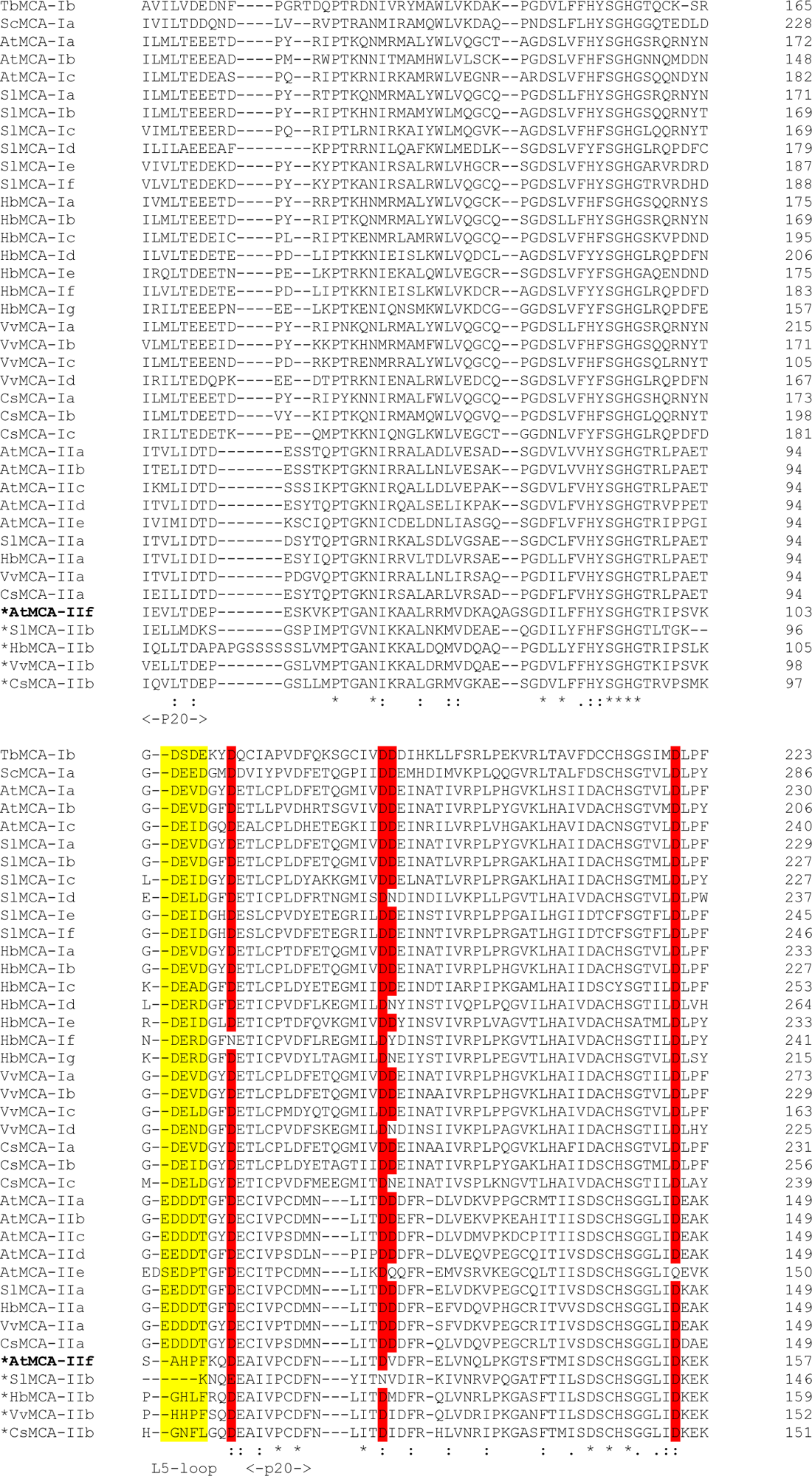
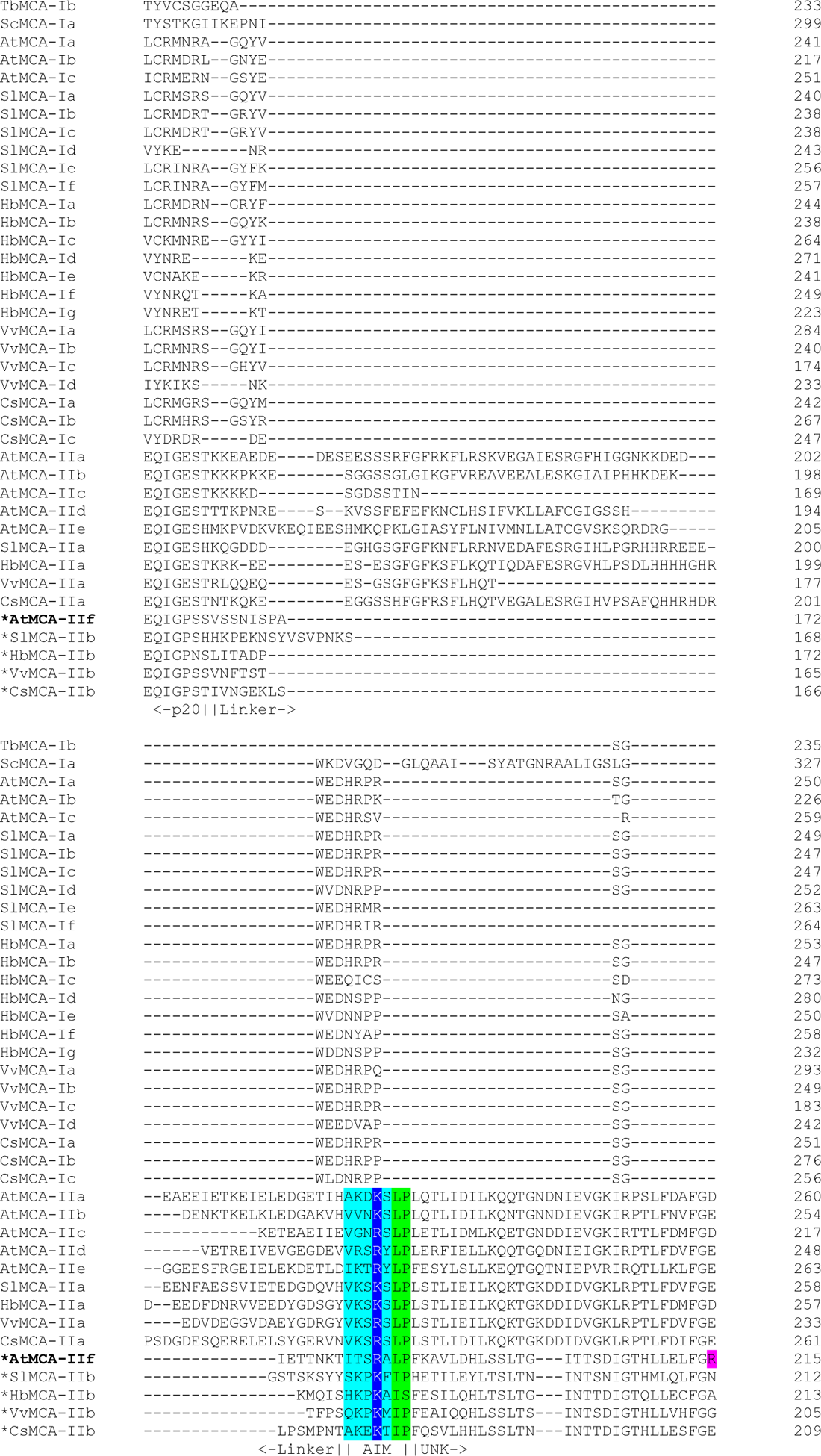
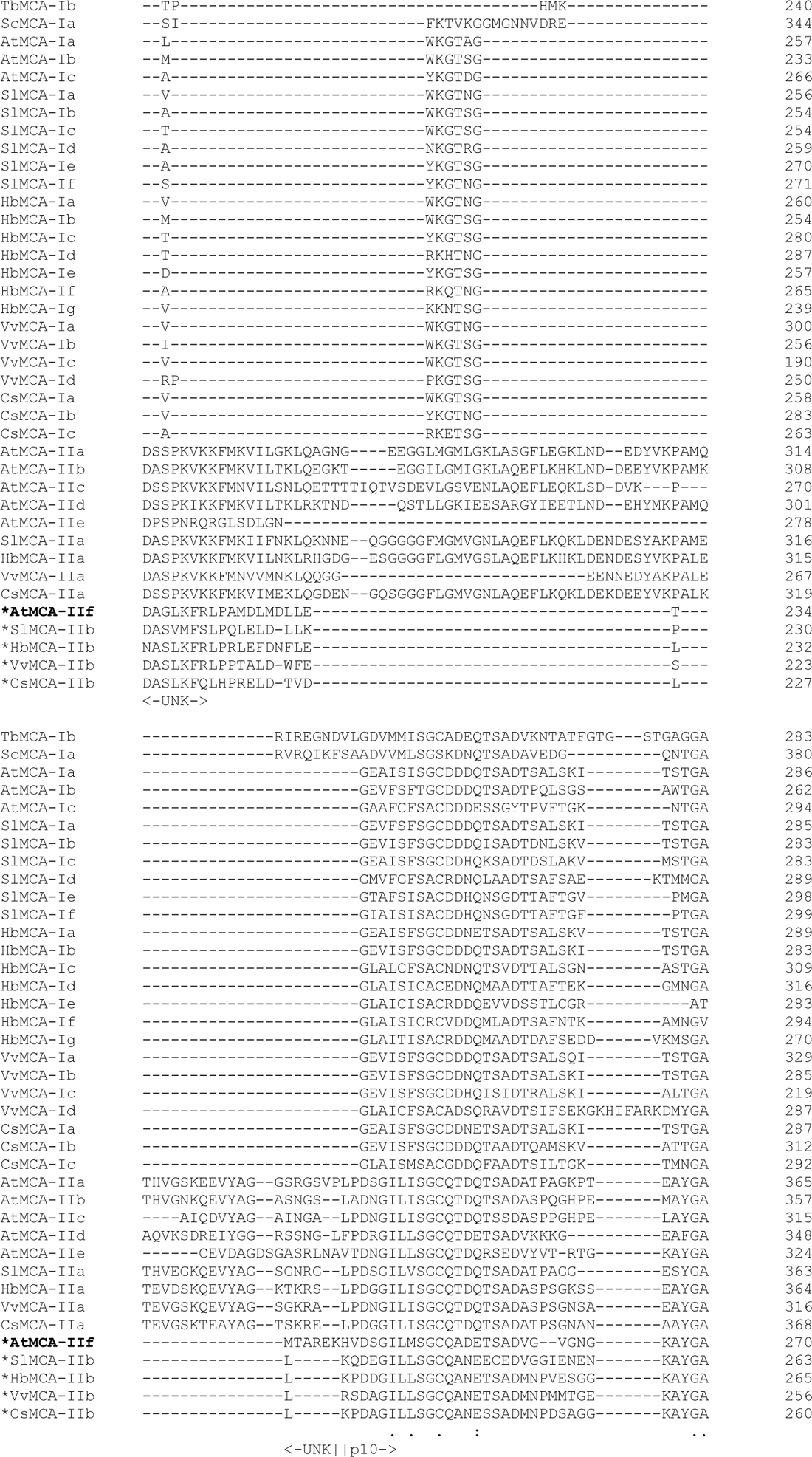
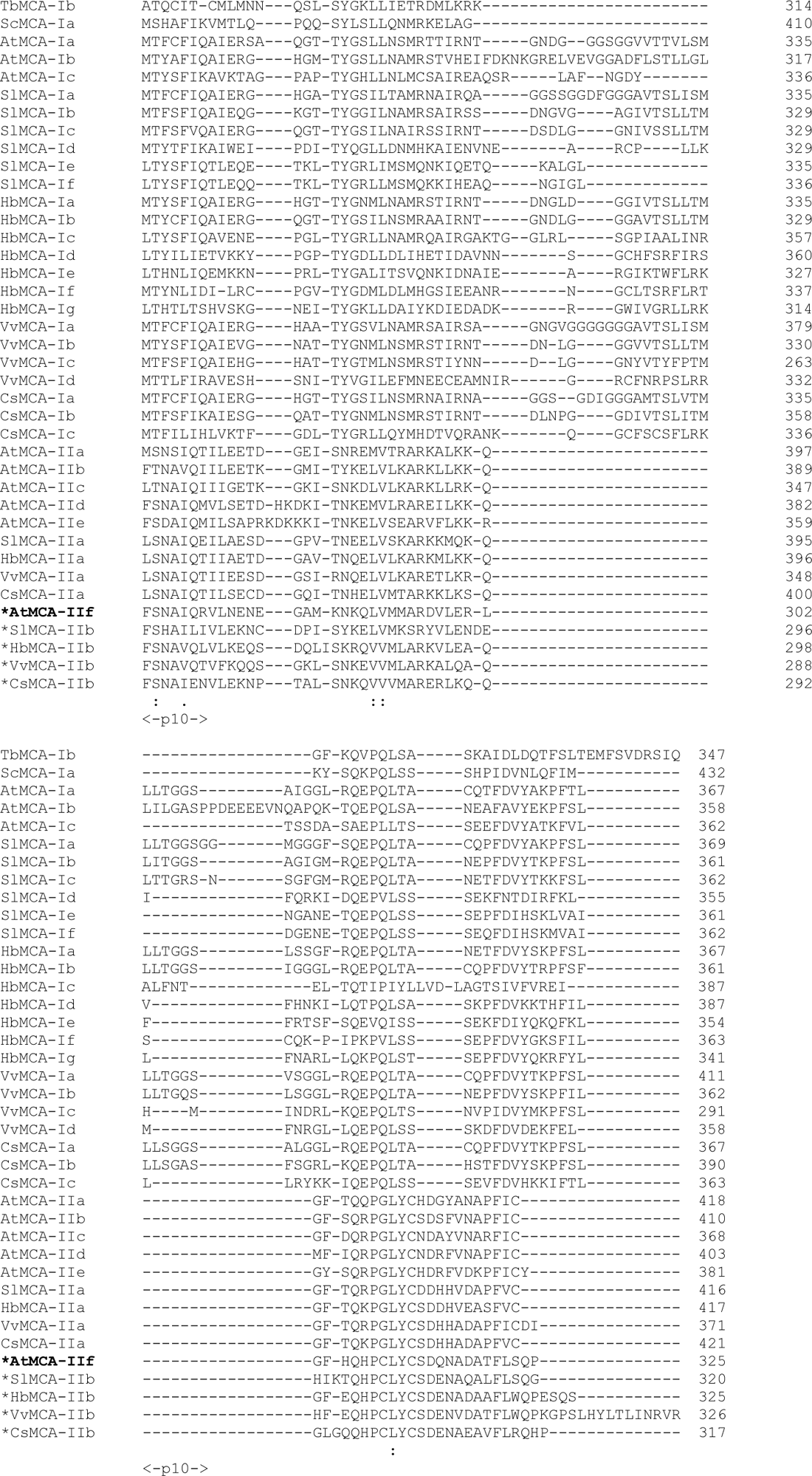
Multiple sequence alignment of type I and type II MCAs. The MCAs from angiosperms: *Arabidopsis thaliana*, *Solanum lycopersicum*, *Hevea brasiliensis*, *Vitis vinifera*, and *Cucumis sativus* were chosen based on [6], and type I MCA from *Trypanosoma brucei* (TbMCA-Ib) and *Saccharomyces cerevisiae* (ScMCA-Ia) were added to alignment. The asterisk symbol in front of the name indicates a calcium independent sub-group of type II metacaspases. The negatively charged cluster in L5 loop is highlighted in yellow. The amino acids involved in the calcium binding, based on TbMCA-Ib structure [20], are highlighted in red. The non-conserved part of the AIM is colored light blue, the conserved part is colored green, and the cleavage site is colored dark blue. The secondary cleavage site in the AtMCA-IIf is highlighted in magenta. Sequence alignments were performed using Clustal Omega program [104].

**Supplementary Fig. 5:**
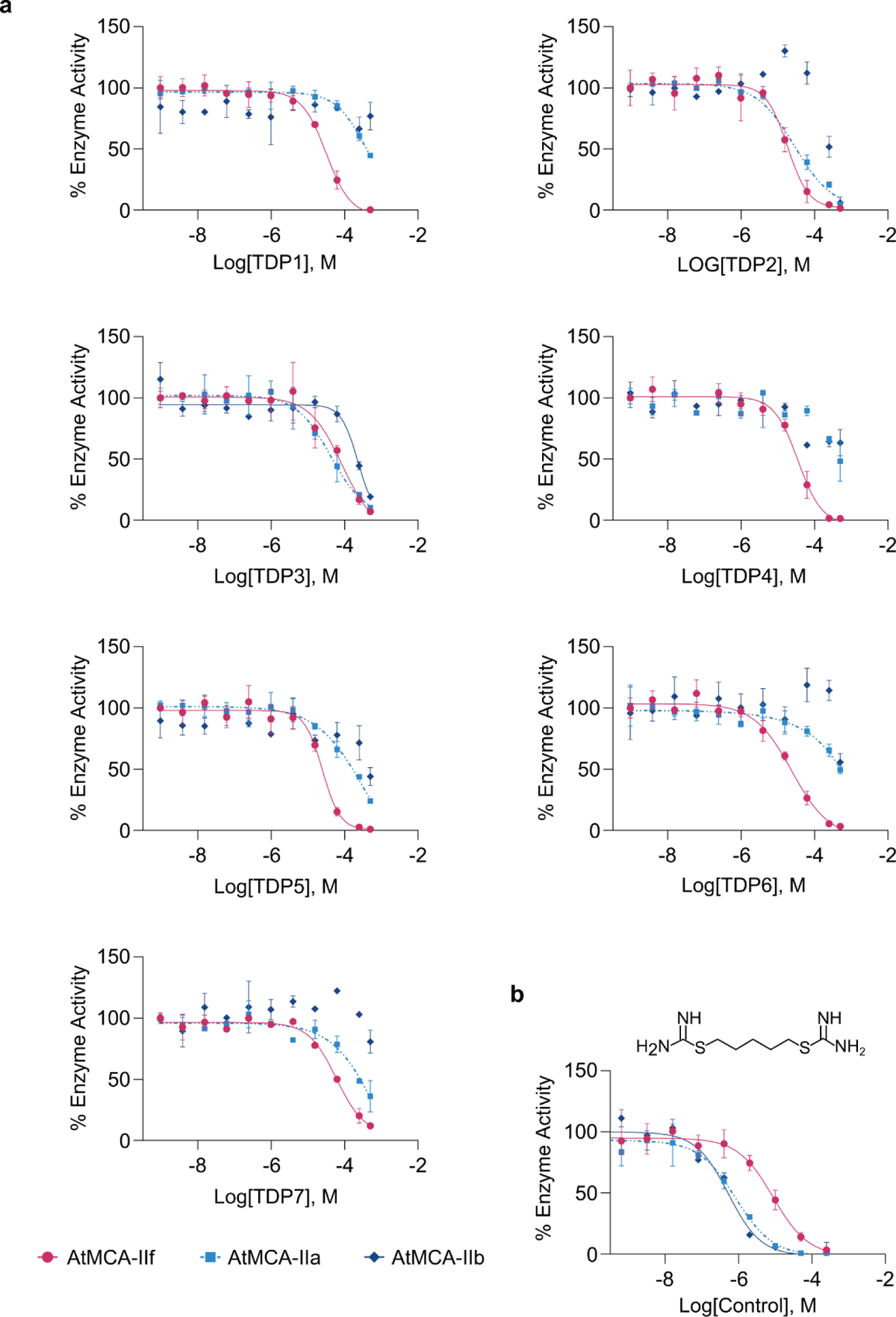
Dose response analysis of AtMCA inhibition by the TDP compounds. **a**, Dose response curves for inhibition by TDP1 to TDP7 of AtMCA-IIa, -IIb and -IIf. The data for TDP6 are also displayed in Fig. 2e. **b,** Dose response analysis of a positive control (an isothiouronium salt) with activity against AtMCA-IIa, -IIb and -IIf.

**Supplementary Fig. 6:**
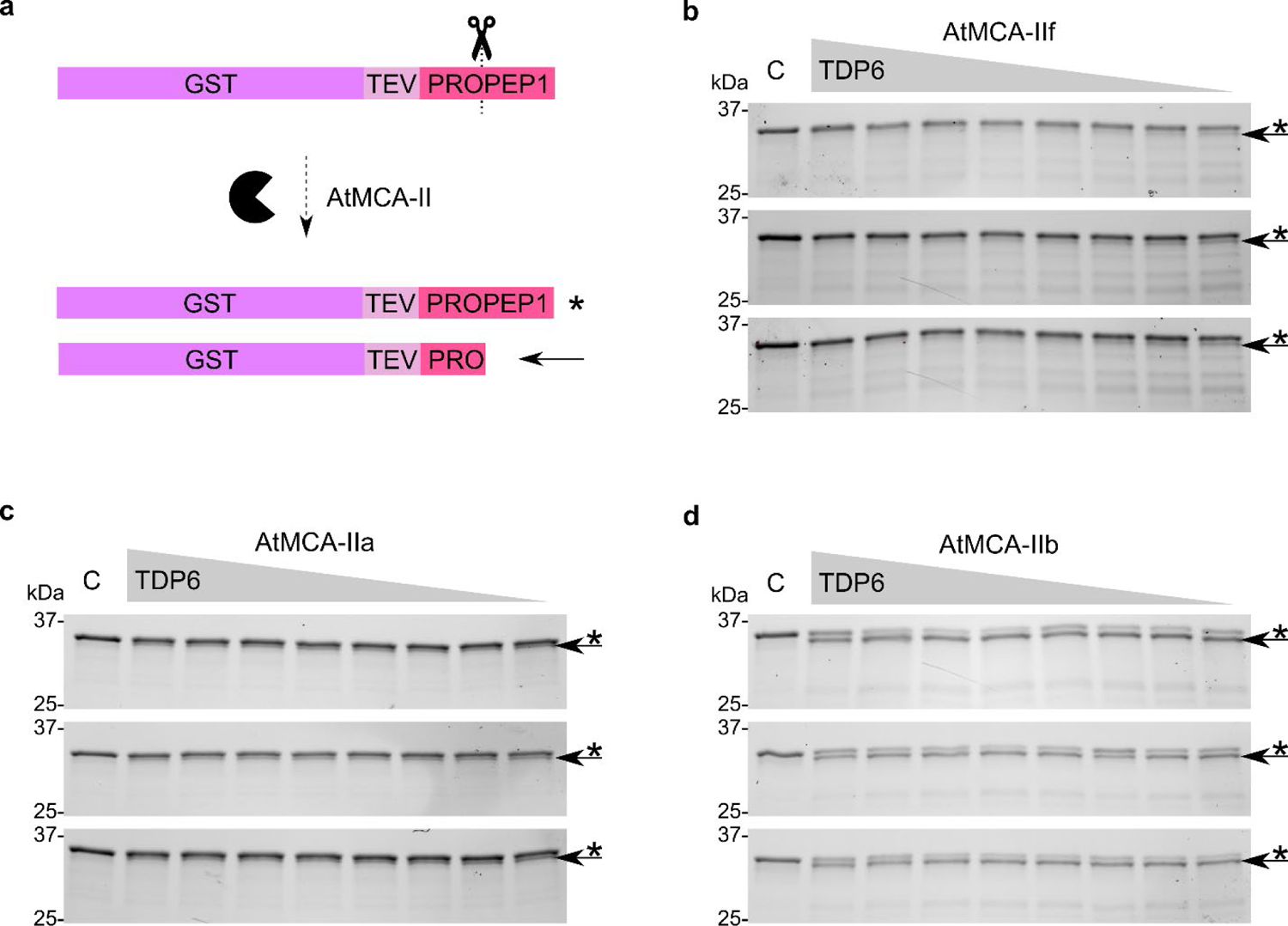
Inhibition of GST-TEV-PROPEP1 cleavage by TDP6. **a**, Schematic representation of the GST-TEV-PROPEP1 fusion protein (*) and cleavage product (arrow) by MCA. **b-d**, Visualization of GST-TEV-PROPEP1 on SDS-PAGE gels of the GST-TEV-PRO cleavage band (indicated with arrow) in the presence of a TDP6 concentration gradient for AtMCA-IIf (**b**), AtMCA-IIb (**c**), and AtMCA-IIa (**d**). The experiment was performed in triplicate, hence the three SDS-PAGE results per protease.

**Supplementary Fig. 7:**
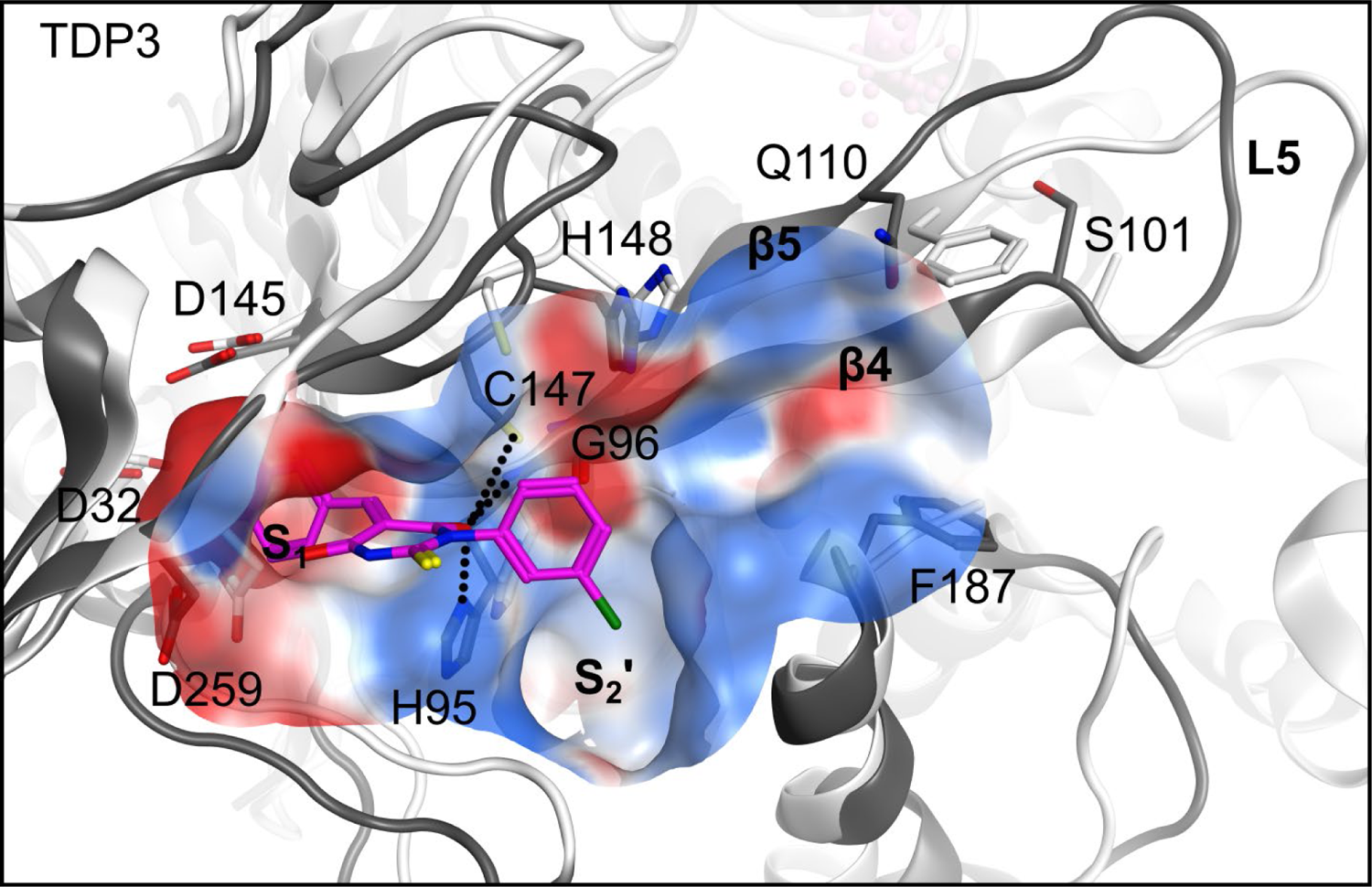
Docking of TDP3 to AtMCA-IIa and f. The tertiary structure and selected residues of AtMCA-IIf (PDB 8A53) are displayed in dark grey and AtMCA-IIa (PDB: 6W8S) in white. Possible hydrogen bonding interactions to the carbonyl oxygens of the thiobarbituric acid moiety are shown with dotted lines. The electrostatic surface of the active site is displayed. The L5 loop of AtMCA-IIf has been modelled for visualization purposes. Residue numbers for AtMCA-IIf are shown.

**Supplementary Fig. 8:**
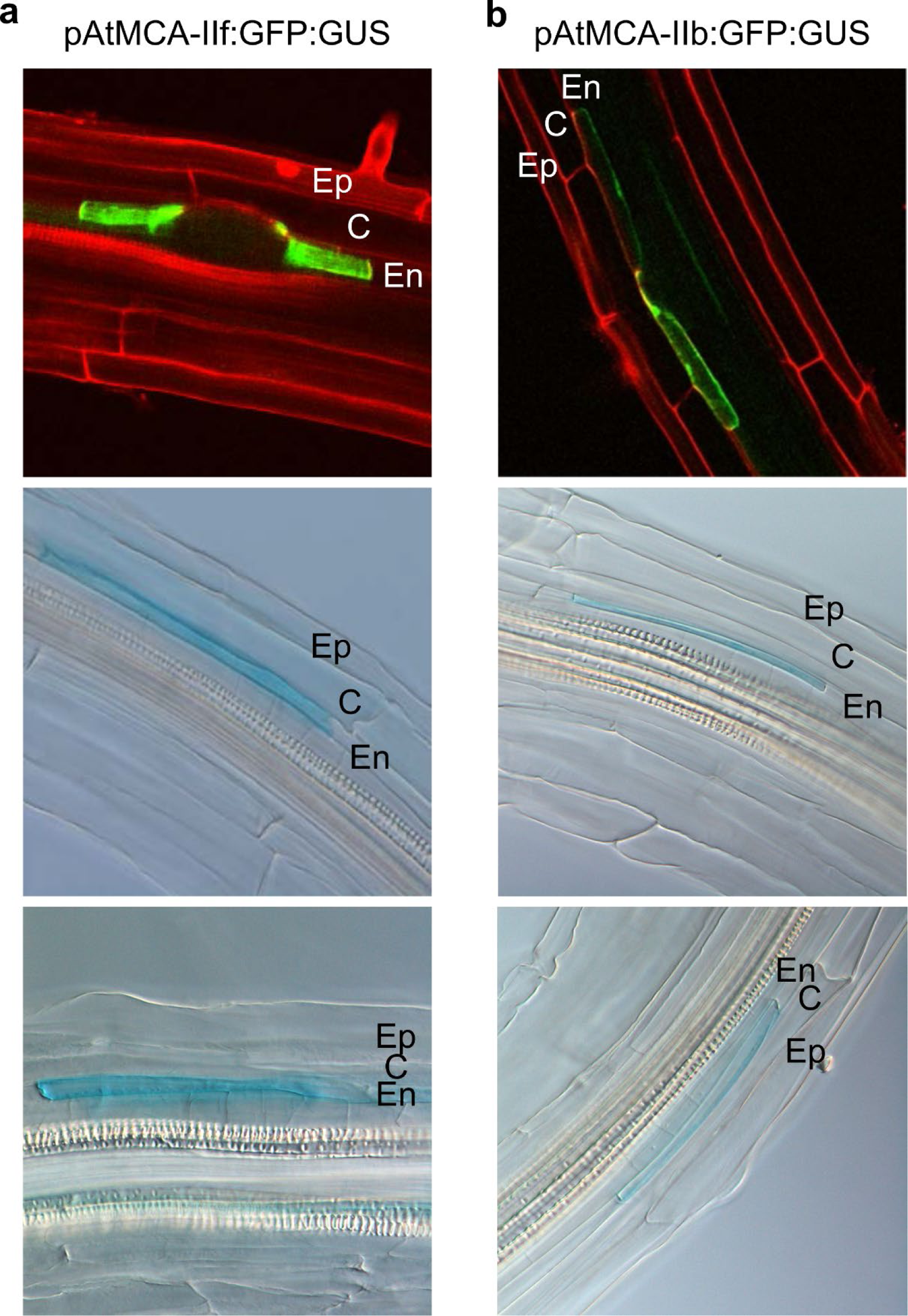
GFP (top panels) and GUS (bottom two panels) expression pattern of AtMCA-IIf. (**a**) and AtMCA-IIb (**b**) in endodermis cells that overlie early stage lateral root primordia. AtMCA promoter fusions to GUS and GFP (pAtMCA:GFP:GUS). En: endodermis, C: cortex, Ep: epidermis.

**Supplementary Fig. 9:**
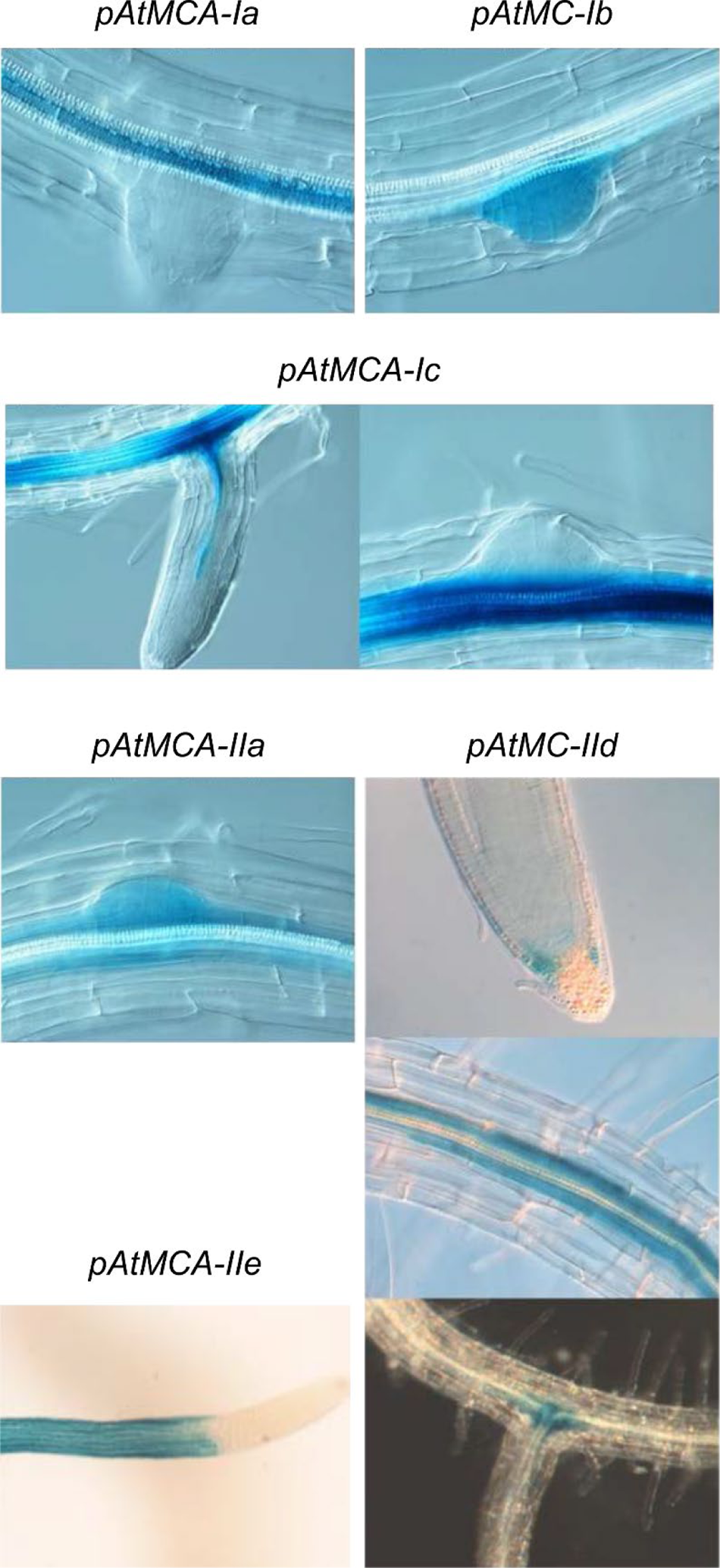
Expression patterns of metacaspase genes in *Arabidopsis* roots. *AtMCA* promoter fusions to GUS and GFP (pAtMCA:GFP:GUS). AtMCA-Ia/AtMC1, AtMCA-Ib/AtMC2, AtMCA-Ic/AtMC3, AtMCA-IIa/AtMC4, AtMCA-IId/AtMC7, AtMCA-IIe/AtMC8.

**Supplementary Fig. 10:**
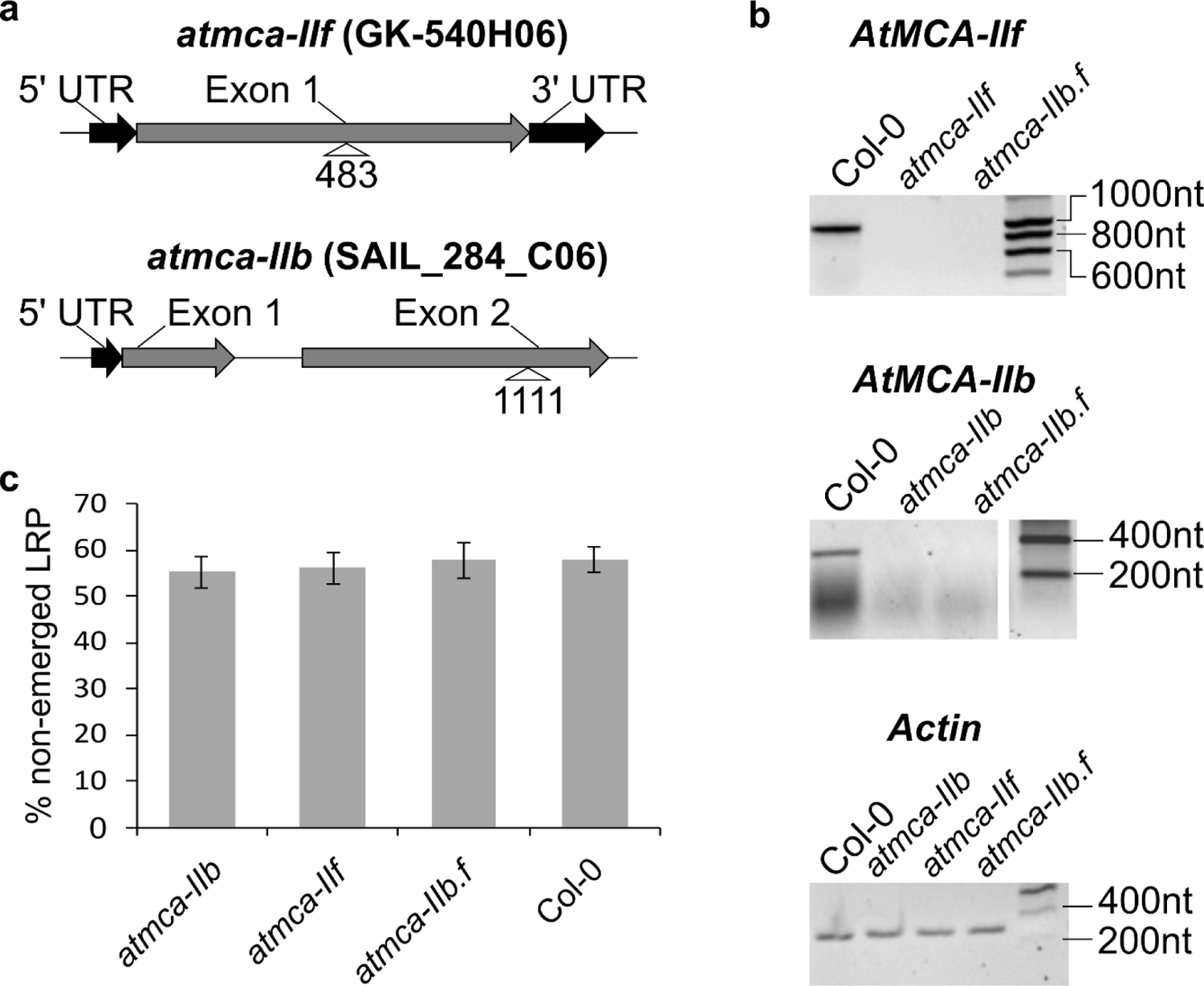
Molecular characterization of the T-DNA insertion lines used in this study. **a**, Schematic representation of the *AtMCA-IIf* and *AtMCA-IIb* genes indicating the location of the T-DNA insertion. Numbering indicates cDNA base pairs downstream of the start codon. **b**, RT-PCR of AtMCA-IIf and AtMCA-IIb mRNA transcripts in the single and double (*atmca-IIb.f*) knock-out lines compared to wild type control (Col-0). Actin was used as a loading control. **c**, No significant differences were observed for non-emerged LRPs in the AtMCA-IIf and AtMCA-IIb single and double knock-out lines compared to Col-0. Values are the mean of 10 measurements ± 95% confidence interval.

**Supplementary Fig. 11:**
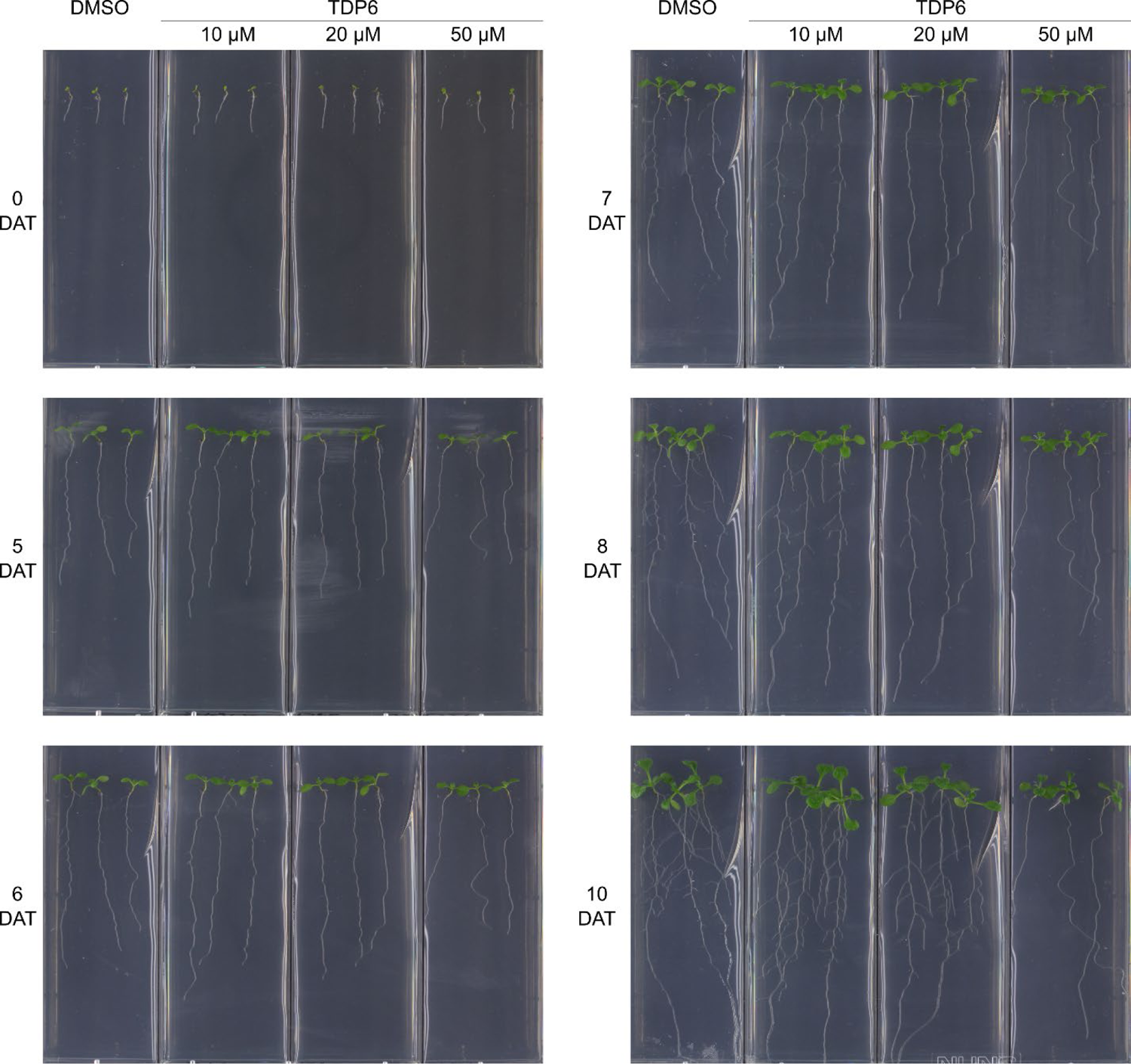
Inhibition of lateral root emergence by TDP6 over time. Wild type Col-0 seedlings transferred after three days of growth on normal medium to medium containing 10, 20, or 50 µM TDP6. The same amount of DMSO as in the 50 µM TDP6 was added to the medium as a control. Assay was repeated twice with similar results. Plates were photographed at the indicated days after transfer (DAT). The image from 7 DAT is used in Fig. 5b.

**Supplementary Fig. 12:**
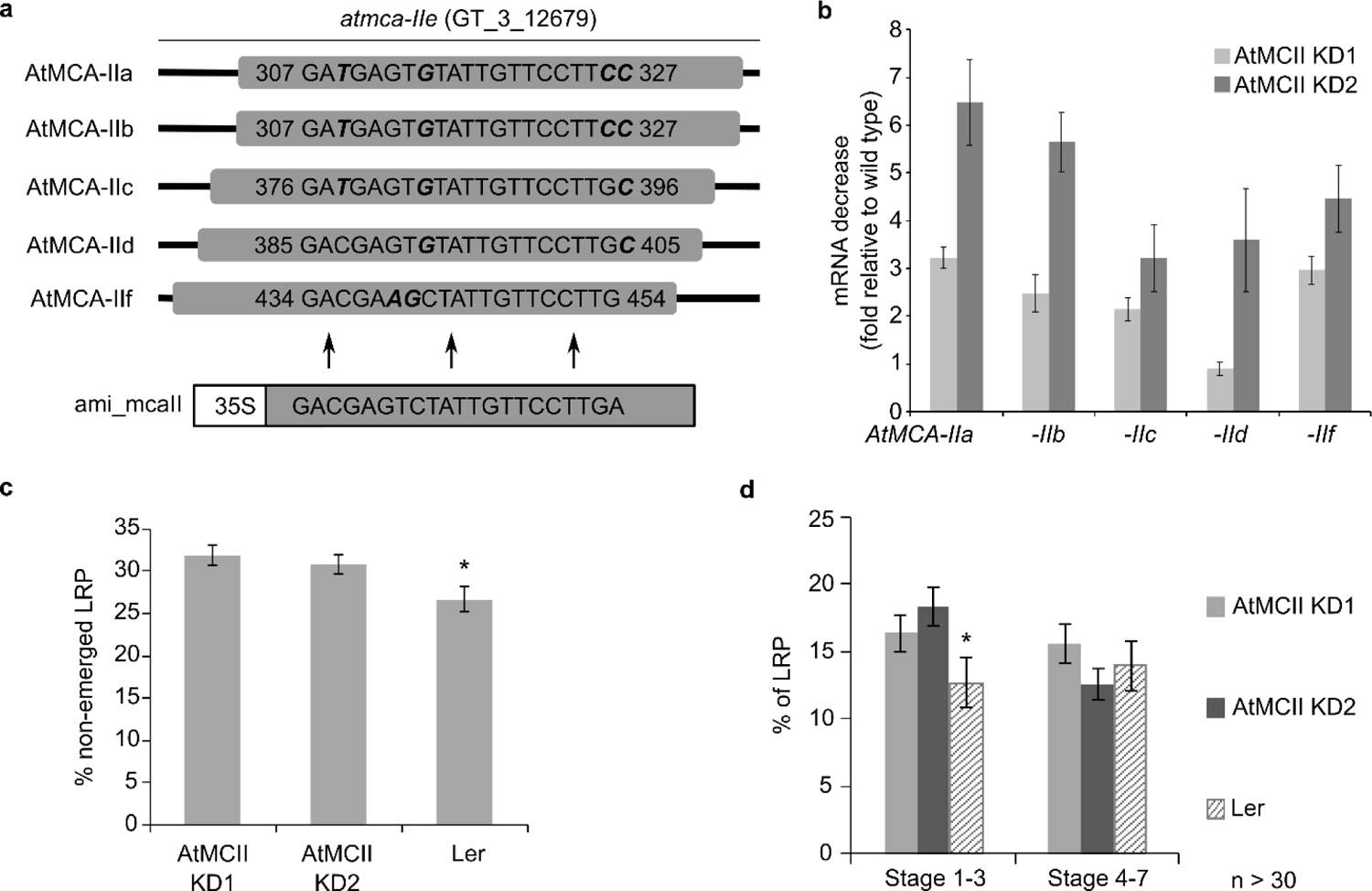
Characterization of LRP development in the metacaspase knock-down lines. **a**, Sequence of the amiRNA construct and its targets in the Arabidopsis type-II metacaspase genes. The amiRNA construct was fused to the CAMV 35S promoter and introduced in an atmca-IIe T-DNA insertion line (JII Gene Trap line GT_3_12679; Landsberg erecta ecotype (Ler)). Mismatched base pairs are indicated in bold and italics. **b**, Decrease fold of each type II metacaspase mRNA in two independent knock-down lines (KD). Values represent the mean ± SE of three biological replicates. **c**, The type-II MC knock-down lines show a significant increase in non-emerged LRPs in comparison to wild type (Ler). **d**, Quantification of LRP stages reveals an increase of in the early stages of LRP development (stages 1-3), when the primordia have not yet protruded the endodermis cell layer, in the knock-down lines. N>30, ± 95% confidence interval. *; p<0.05.

**Supplementary Fig. 13:**
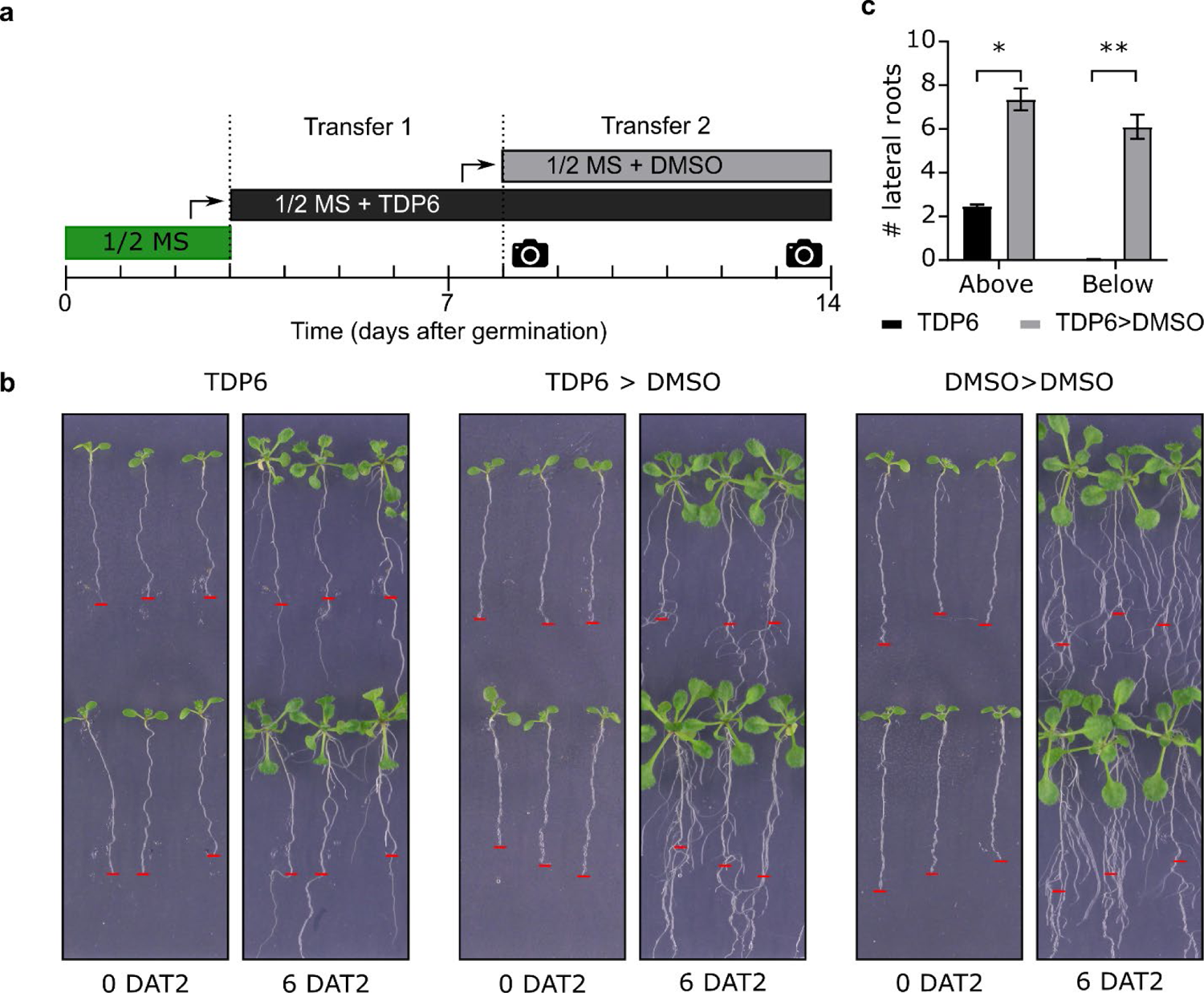
Inhibition of lateral root emergence by TDP6 can be undone by transfer of seedlings to DMSO. **a**, Schematic representation of the seedling transfer experiment. Seedlings were allowed to germinate on plates with standard half-strength Murashige and Skoog (1/2 MS) growth medium before a first transfer to plates containing 50 µM TDP6 and a second transfer to DMSO. Pictures in (b) were taken at the indicated timepoints by a camera symbol. **b**, Representative pictures of the transfer experiment at 0 and 6 days after transfer 2 (DAT2). The position of the root tip at 0 DAT2 was indicated with a red stripe and transposed to the 6 DAT2 picture. **c**, The number of lateral roots visual per seedling above and below the red stripe were counted at 6 DAT (two biological repeats each containing 20 seedlings; unpaired t test, p value < 0.05* or < 0.01**). As control, a transfer experiment from DMSO to DMSO is displayed in (b), but this was not quantified.

**Supplementary Fig. 14:**
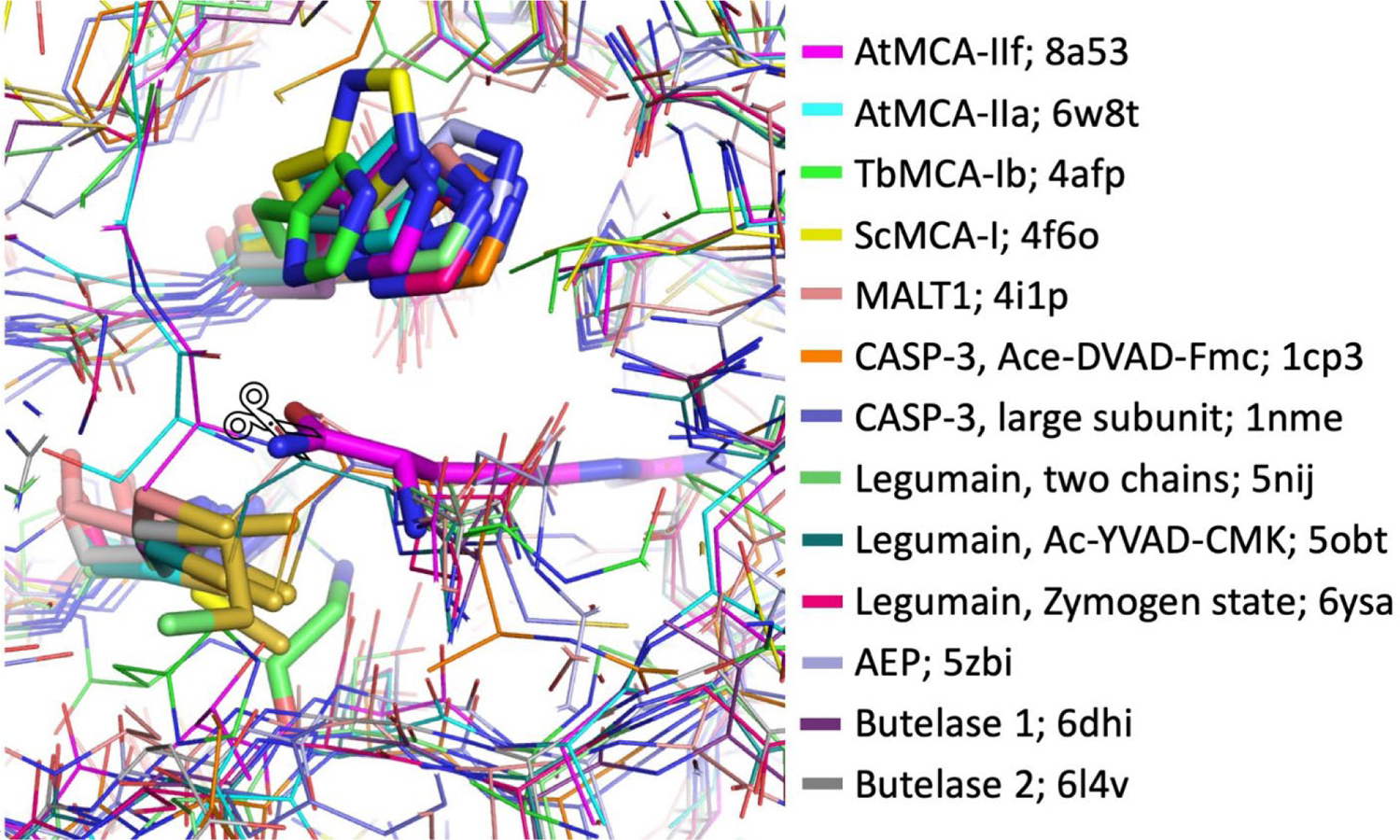
Structure superposition of the His-Cys dyad of various CD-clan members. No major structure changes occur at the catalytic center between inactive mutants, zymogen forms, activated enzyme or trapped covalent intermediates. Histidine-cysteine dyads are shown in stick representation, as is Arg183 from AtMCA-IIf and the scissile peptide bond to be cleaved, indicated by the scissors. For inactive Cys-mutants, the residue is shown that replaces the catalytic cysteine, such as glycine in TbMCA-Ib. AtMCA-IIf, calcium independent type II metacaspase from *A. thaliana*; AtMCA-IIa, calcium dependent type II metacaspase from *A. thaliana*; TbMCA-Ib, type I metacaspase from *T. brucei*; ScMCA-I, type I metacaspase from *S. cerevisiae*; MALT1, paracaspase from *Homo sapiens*; CASP-3, caspase-3 from *H. sapiens*; Legumains are from *A. thaliana*; Butelases are from *Clitoria ternatea*; AEP, asparaginyl endopeptidase from *Viola canadensis*.

