## Supplemental Table 1 for "Structure-function analysis of a calcium-independent metacaspase reveals a novel proteolytic pathway for lateral root emergence"

**Supplementary Table 1. Crystal parameters and statistics from diffraction data processing and structure refinement^a^.**

|  | | | AtMCA-IIa C147A |
| --- | --- | --- | --- |
| Crystal parameters | | |  |
|  | Resolution (Å) | | 1.95 (44.38) |
|  | Space group | | *P*12_1_1 |
|  | Unit cell parameters | |  |
|  | *a*, *b*, *c* (Å) | | 81.2, 88.1, 82.5 |
|  | *α*, *β*, *γ* (°) | | 90.0, 102.3, 120 |
|  | Matthews coefficient (Å^3^/Da) | | 1.88 |
|  | Solvent contentc (%) | | 34.6 |
| Data collection | | |  |
|  | Completeness (%) | | 98.3 (96.8) |
|  | No. of unique reflections | | 81454 |
|  | *I*/σ(*I*) | | 21.4 (3.2) |
|  | R_merge_ (%) | | 4.4 (59.0) |
|  | CC_1/2_(%) | | 100 (92.1) |
|  | Redundancy | | 7.1 (7.30) |
|  | Wilson B-factor (Å^2^) | | 32.9 |
| Refinement | | |  |
|  | *R* (%)/ *R*_free_ (%) | | 17.4/20.8 |
|  | RMSD bonds length (Å) | | 0.004 |
|  | Average *B* factor | | 45.0 |
|  | Number of molecules in AU | | 4 |
|  | No. of atoms | |  |
|  | | Protein | 9284 |
|  | | Nitrate | 12 |
|  | | Water molecules | 416 |
|  | Ramachandran analysis^b^ | |  |
|  | | Favored (%)/*n* | 98 |
|  | | Allowed (%)/*n* | 2 |
|  | | Outlier (%)/*n* | 0 |

^a^ Data for the highest resolution shell is given in parentheses.

The abbreviations RMSD and AU stand for root-mean-square deviation and asymmetric unit, respectively.

^b^ Defined by validation program MOLPROBITY (Richardson et al., 2010).
