## Supplemental Table 4 for "Structure-function analysis of a calcium-independent metacaspase reveals a novel proteolytic pathway for lateral root emergence"

|  |  | IC_50_ (µM) ^a^ |  |
| --- | --- | --- | --- |
|  | AtMCA-IIf | AtMCA-IIa | AtMCA-IIb |
| TDP1 | 31  (24-42) | >200 | Inactive^b^ |
| TDP2 | 19  (14-27) | 29  (17-77) | Inactive^b^ |
| TDP3 | 86  (43-120) | 41  (21–250) | >200 |
| TDP4 | 35  (27-51) | Inactive^b^ | Inactive^b^ |
| TDP5 | 25  (19-33) | >200 | Inactive^b^ |
| TDP6 | 24  (16–43) | >200 | Inactive^b^ |
| TDP7 | 60  (43-100) | >200 | Inactive^b^ |
| Positive control | 0.6  (0.4–0.9) | 0.8  (0.5–1.5) | 8.7  (5.4–18) |

^a^ 95 % confidence interval is given within parentheses;

^b^ Inactive within the tested concentration span
