## Supplemental Table 5 for "Structure-function analysis of a calcium-independent metacaspase reveals a novel proteolytic pathway for lateral root emergence"

#### NMR characterization

TDP1 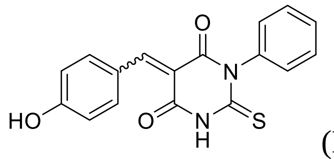

(E/Z)-5-(4-hydroxybenzylidene)-1-phenyl-2-thioxodihydropyrimidine-4,6(1H,5H)-dione. E/Z mixture 1:1. $^{1}$H NMR (400 MHz, DMSO-$d_6$) δ 12.64 (s, 1H, E+Z, NH), 10.97 (s, 1H, E+Z, OH), 8.49 – 8.27 (m, E+Z, 2H), 8.24 (s, 1H, E+Z, -CH=C), 7.55 – 7.36 (m, 3H, E+Z), 7.27 (t, *J* = 7.3 Hz, E+Z, 2H), 6.89 (dd, *J* = 21.3, 8.5 Hz, E+Z, 2H).

TDP2 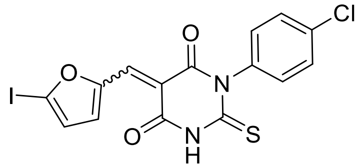

(E/Z)-1-(4-chlorophenyl)-5-((5-iodofuran-2-yl)methylene)-2-thioxodihydropyrimidine-4,6(1H,5H)-dione. E/Z mixture 1:1. $^{1}$H NMR (400 MHz, DMSO-$d_6$) δ 12.79 (s, *2*H, E+Z, NH), 8.42 (d, *J* = 3.8 Hz, 1H, E or Z), 8.20 (d, *J* = 3.8 Hz, 1H, Z or E), 8.08 (s, 1H, -CH=C, E or Z), 7.99 (s, 1H, -CH=C, Z or E), 7.57 – 7.49 (m, 4H, E+Z), 7.38 – 7.29 (m, 4H, E+Z), 7.22 (dd, *J* = 3.9, 0.8 Hz, 1H, E or Z), 7.14 (dd, *J* = 3.9, 0.8 Hz, 1H, Z or E).

TDP3 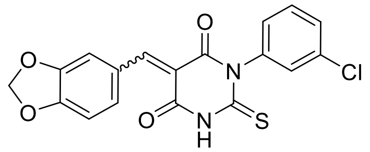

(E/Z)-5-(benzo[d][1,3]dioxol-5-ylmethylene)-1-(3-chlorophenyl)-2-thioxodihydropyrimidine-4,6(1H,5H)-dione. E/Z mixture 1:1. $^{1}$H NMR (400 MHz, DMSO-$d_6$) δ 12.78 (s, 1H, NH, E or Z), 12.70 (s, 1H, NH, Z or E), 8.34 (s, 1H, -CH=C, E or Z), 8.33 (d, *J* = 1.8 Hz, 1H, E or Z), 8.24 (s, 1H, -CH=C, Z or E), 8.08 (d, *J* = 1.7 Hz, 1H, Z or E), 7.84 – 7.82 (m, 1H, E or Z), 7.82 – 7.80 (m, 1H, Z or E), 7.51 – 7.43 (m, 6H, E+Z), 7.30 (tt, *J* = 6.7, 2.3 Hz, 2H, E+Z), 7.12 (d, *J* = 8.2 Hz, 1H, E or Z), 7.08 (d, *J* = 8.2 Hz, 1H, Z or E), 6.22 (s, 2H, E or Z), 6.17 (s, 2H, Z or E).

TDP4 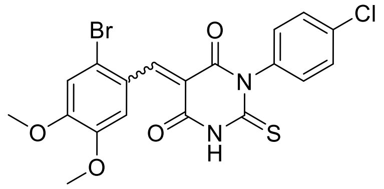

(E/Z)-5-(2-bromo-4,5-dimethoxybenzylidene)-1-(4-chlorophenyl)-2-thioxodihydropyrimidine-4,6(1H,5H)-dione. E/Z mixture 1:1. $^{1}$H NMR (400 MHz, DMSO-$d_6$) δ 12.84 (s, 1H, NH, E or Z), 12.71 (s, 1H, NH, Z or E), 8.42 (s, 2H, E+Z), 8.07 (s, 1H, E or Z), 7.78 (s, 1H, Z or E), 7.52 (dd, *J* = 13.0, 8.7 Hz, 4H, E+Z), 7.39 – 7.29 (m, 6H, E+Z), 3.90 (s, 3H, E or Z), 3.86 (s, 3H, Z or E), 3.78 (s, 3H, E or Z), 3.63 (s, 3H, Z or E).

TDP5 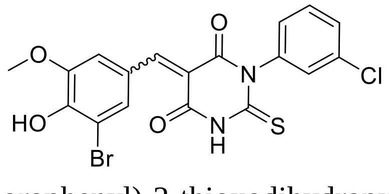

(E/Z)-5-(3-bromo-4-hydroxy-5-methoxybenzylidene)-1-(3-chlorophenyl)-2-thioxodihydropyrimidine-4,6(1H,5H)-dione. E/Z mixture 1:1, described as a single isomer. $^{1}$H NMR (400 MHz, DMSO-$d_6$) δ 12.71 (bs, 1H, NH), 8.47 (bs, 1H), 8.26 (bs, 1H), 8.20 – 7.99 (m, 1H), 7.56 – 7.41 (m, 3H), 7.30 (dt, *J* = 6.8, 2.1 Hz, 1H), 3.87 (s, 3H).

TDP6 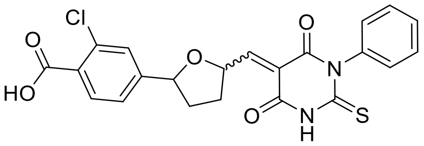 (E/Z)-2-chloro-4-(5-((4,6-dioxo-1-phenyl-2-thioxotetrahydropyrimidin-5(2H)-ylidene)methyl)furan-2-yl)benzoic acid. E/Z mixture 1:1. $^{1}$H NMR (400 MHz, DMSO-$d_6$) δ 13.71 (bs, 2H, COOH, E+Z), 12.77 (s, 1H, NH, E or Z), 12.72 (s, 1H, NH, Z or E), 8.67 (d, *J* = 4.0 Hz, 1H, E or Z), 8.51 (d, *J* = 4.0 Hz, 1H, Z or E), 8.32 (t, *J* = 2.2 Hz, 2H, E+Z), 8.26 (s, 1H, E or Z), 8.15 (s, 1H, Z or E), 8.11 (ddd, *J* = 8.2, 5.5, 2.3 Hz, 2H, E+Z), 7.72 (dd, *J* = 8.5, 3.7 Hz, 2H, E+Z), 7.66 (d, *J* = 4.0 Hz, 1H, E or Z), 7.59 (d, *J* = 3.9 Hz, 1H, Z or E), 7.51 – 7.45 (m, 4H, E+Z), 7.41 (ddd, *J* = 5.8, 3.3, 1.8 Hz, 2H, E+Z), 7.29 (ddd, *J* = 9.8, 7.6, 1.4 Hz, 4H, E+Z).

TDP7 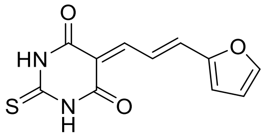 (E)-5-(3-(furan-2-yl)allylidene)-2-thioxodihydropyrimidine-4,6(1H,5H)-dione. $^{1}$H NMR (400 MHz, DMSO-$d_6$) δ 12.30 (s, 1H, NH), 12.26 (s, 1H, NH), 8.21 (dd, *J* = 15.0, 12.4 Hz, 1H), 8.04 (s, 1H), 8.02 (d, *J* = 10.2 Hz, 1H), 7.63 (d, *J* = 15.1 Hz, 1H), 7.09 (d, *J* = 3.5 Hz, 1H), 6.75 (dd, *J* = 3.5, 1.8 Hz, 1H).

### NMR spectra

#### TDP1

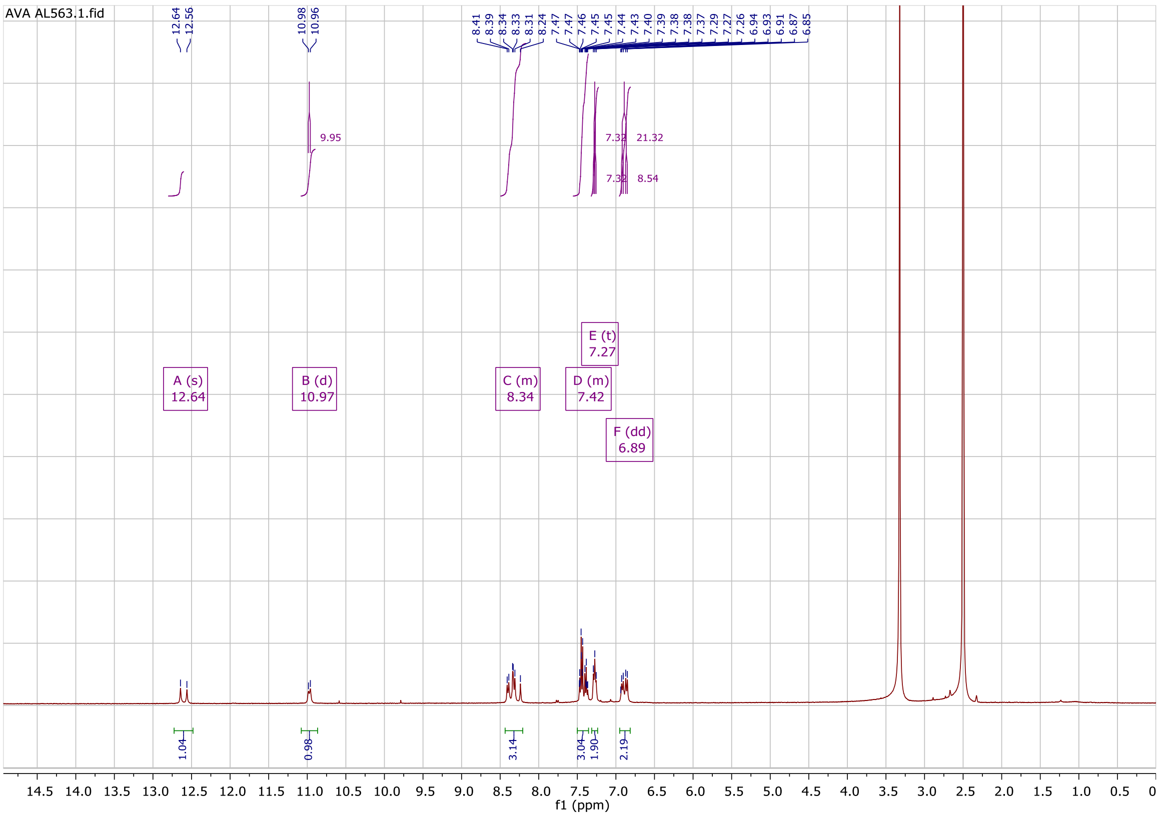


#### TDP2

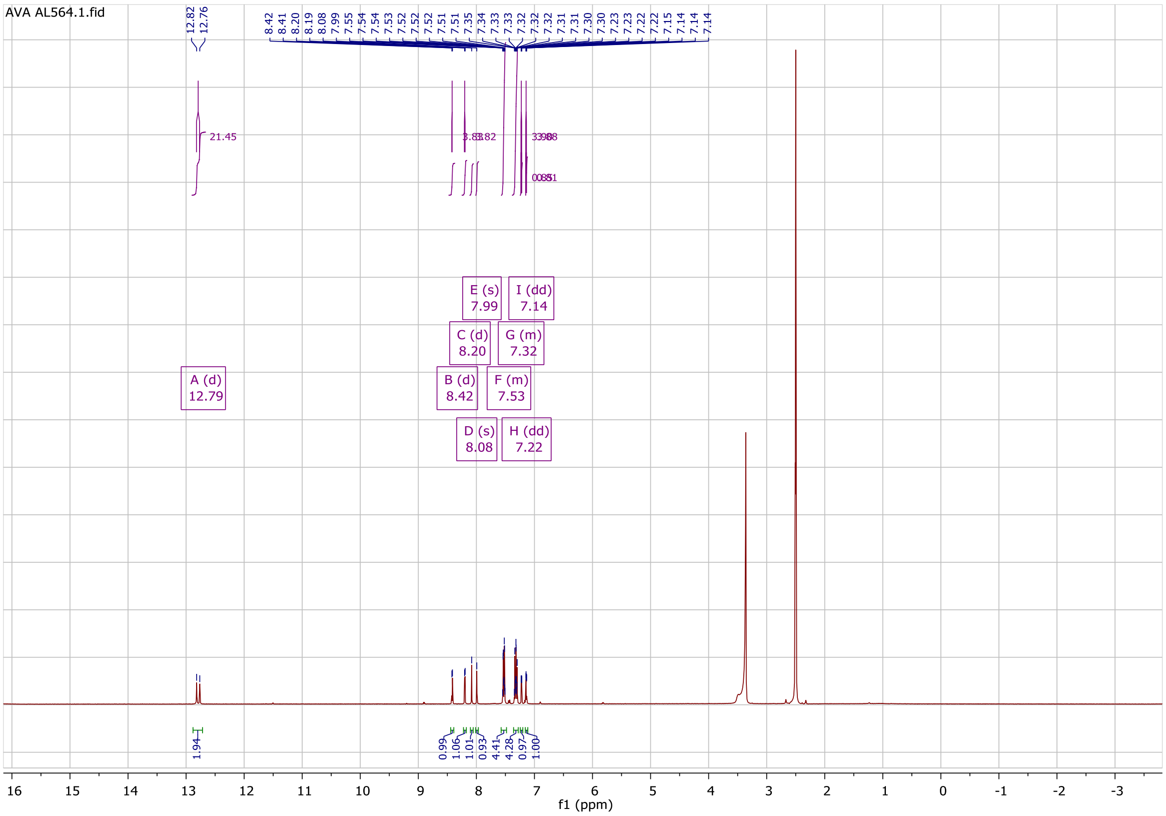

TDP3

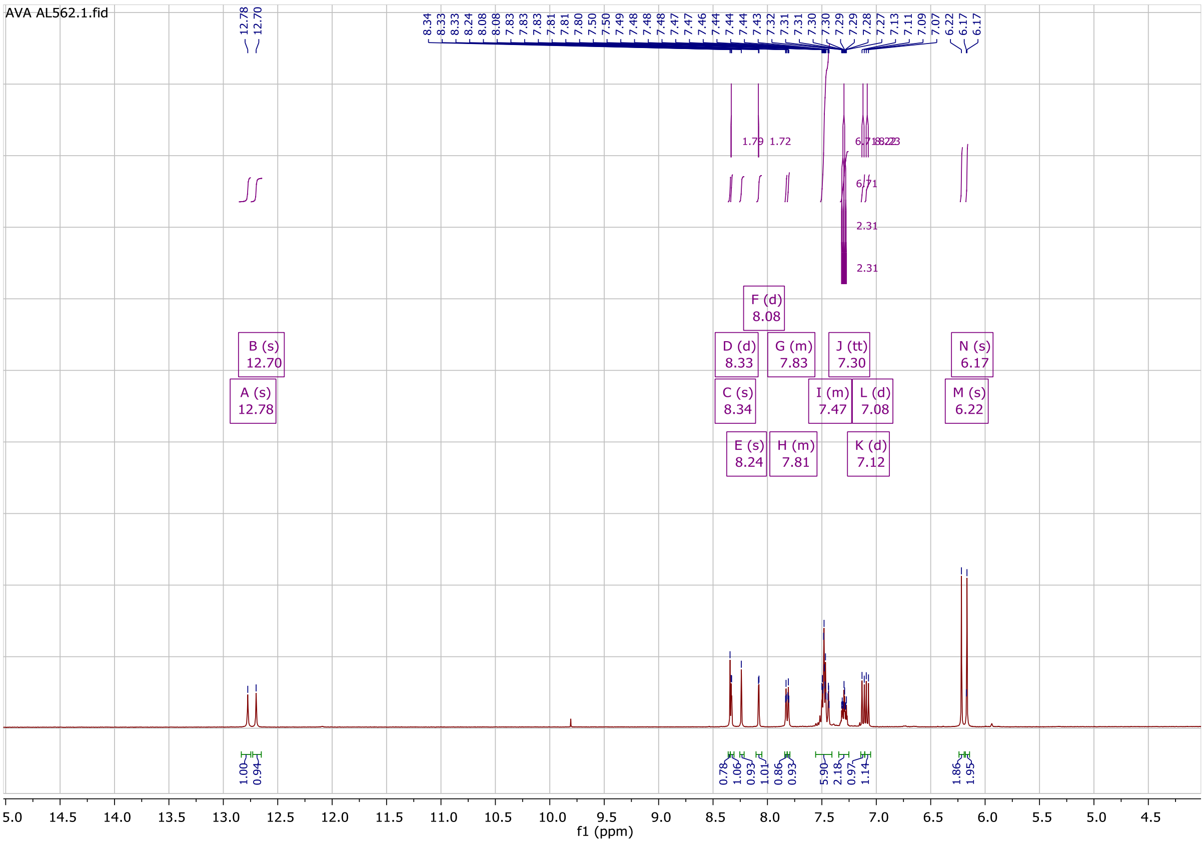
AVA AL562.1.fid
A (s) 12.78
B (s) 12.70
C (s) 8.34
D (d) 8.33
E (s) 8.24
F (d) 8.08
G (m) 7.83
H (m) 7.81
I (m) 7.47
J (tt) 7.30
K (d) 7.12
L (d) 7.08
M (s) 6.22
N (s) 6.17
f1 (ppm)

TDP4

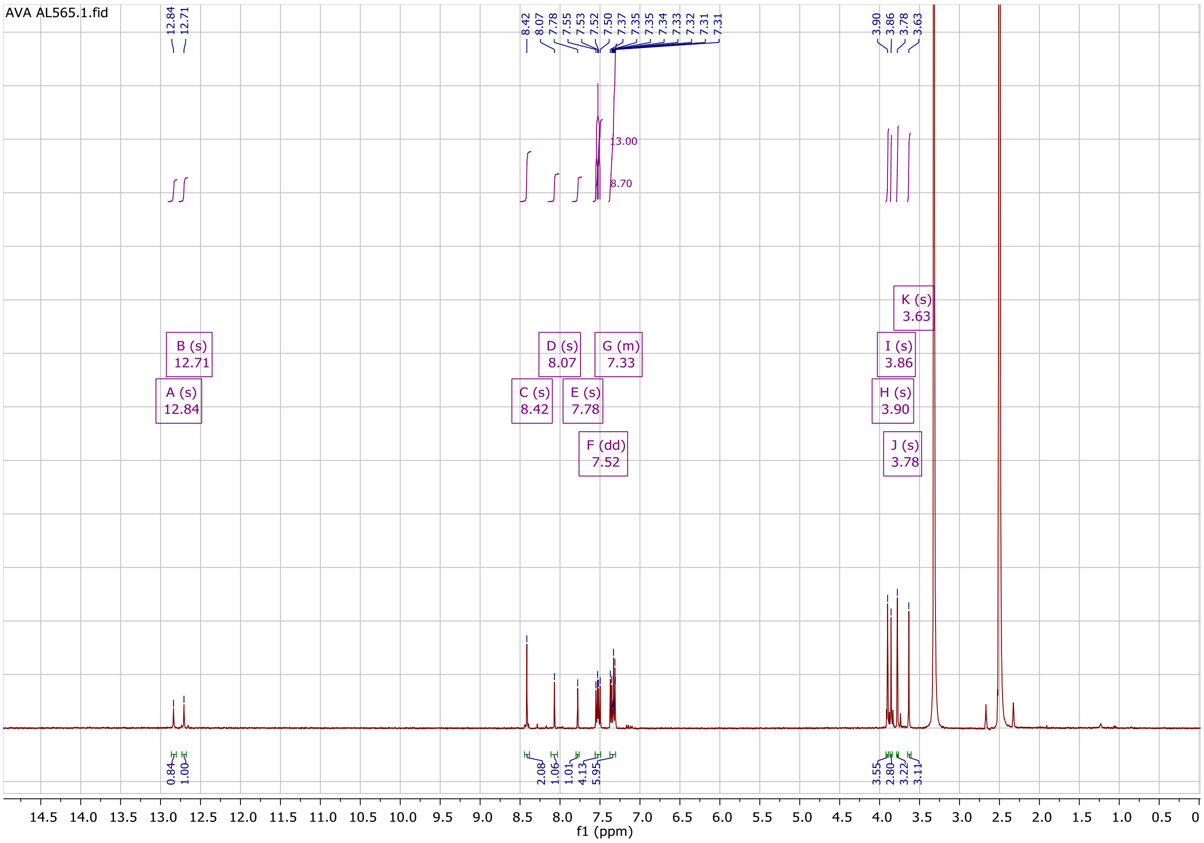
AVA AL565.1.fid
A (s) 12.84
B (s) 12.71
C (s) 8.42
D (s) 8.07
E (s) 7.78
F (dd) 7.52
G (m) 7.33
H (s) 3.90
I (s) 3.86
J (s) 3.78
K (s) 3.63
f1 (ppm)

TDP5

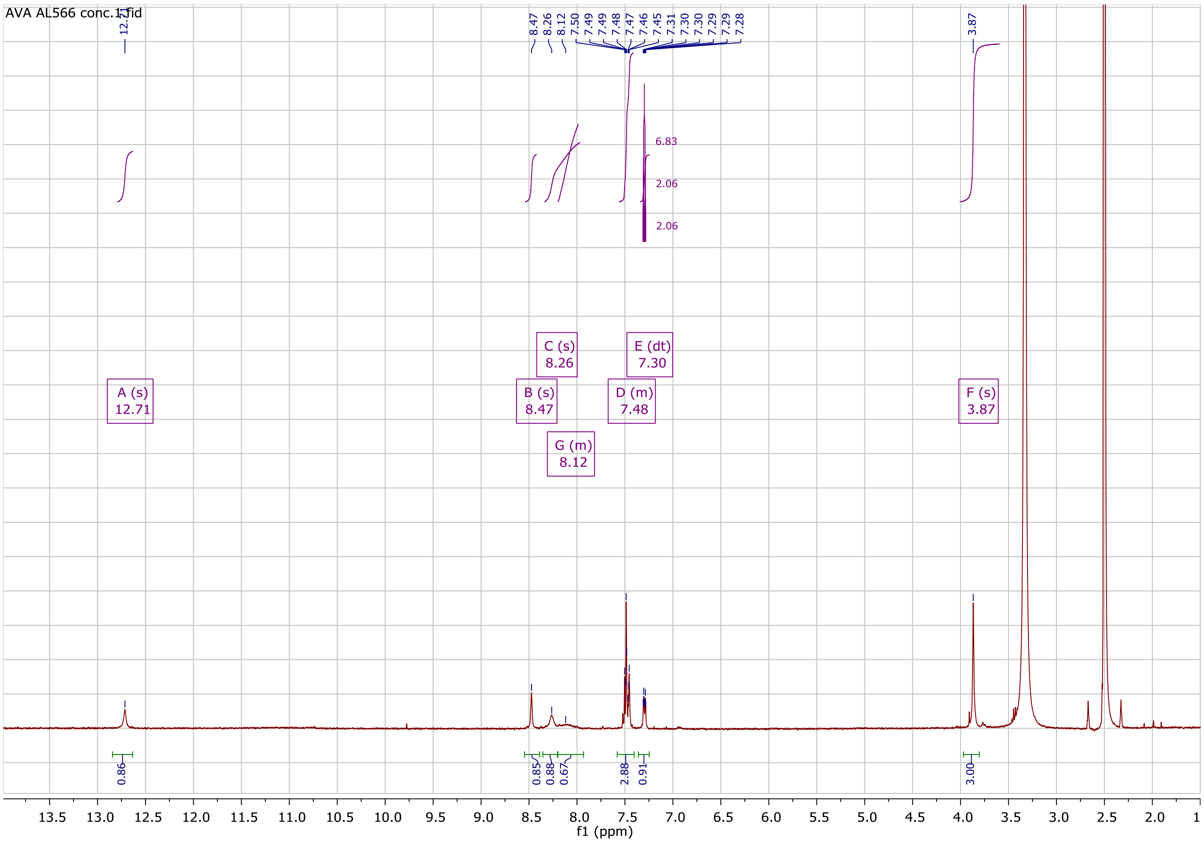
AVA AL566 conc.1.fid
A (s)
12.71
B (s)
8.47
C (s)
8.26
G (m)
8.12
D (m)
7.48
E (dt)
7.30
F (s)
3.87
f1 (ppm)

TDP6

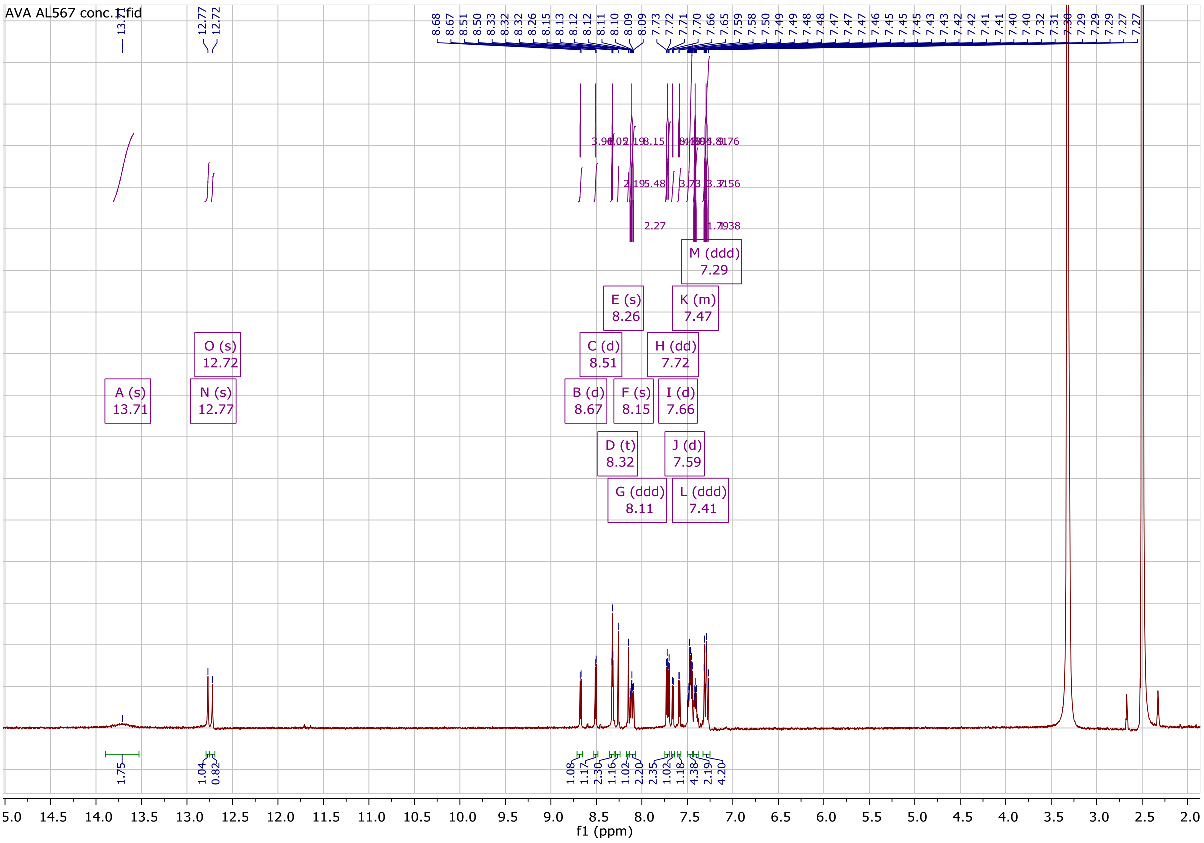
AVA AL567 conc.1.fid
A (s)
13.71
N (s)
12.77
O (s)
12.72
B (d)
8.67
C (d)
8.51
D (t)
8.32
E (s)
8.26
F (s)
8.15
G (ddd)
8.11
H (dd)
7.72
I (d)
7.66
J (d)
7.59
K (m)
7.47
L (ddd)
7.41
M (ddd)
7.29
f1 (ppm)

TDP7

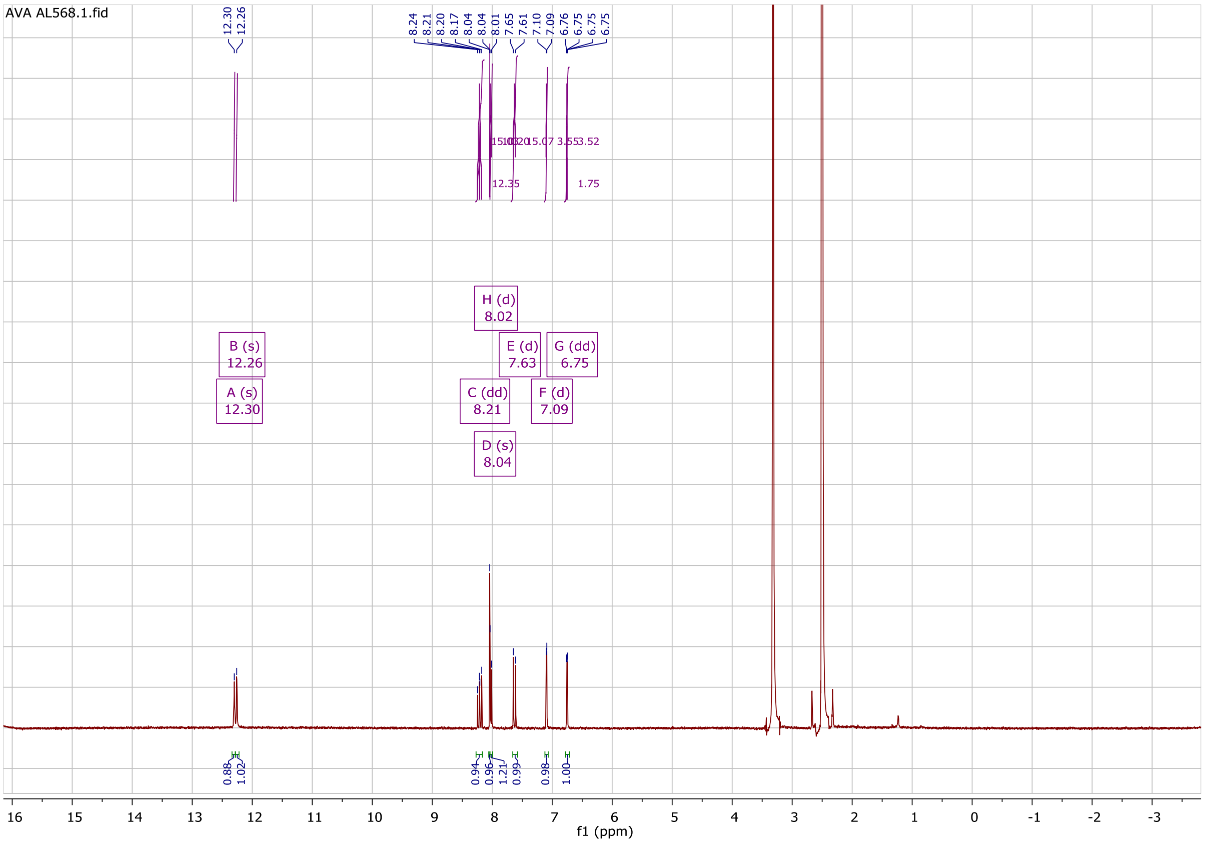
AVA AL568.1.fid
B (s) 12.26
A (s) 12.30
H (d) 8.02
E (d) 7.63
G (dd) 6.75
C (dd) 8.21
F (d) 7.09
D (s) 8.04
f1 (ppm)

COSY

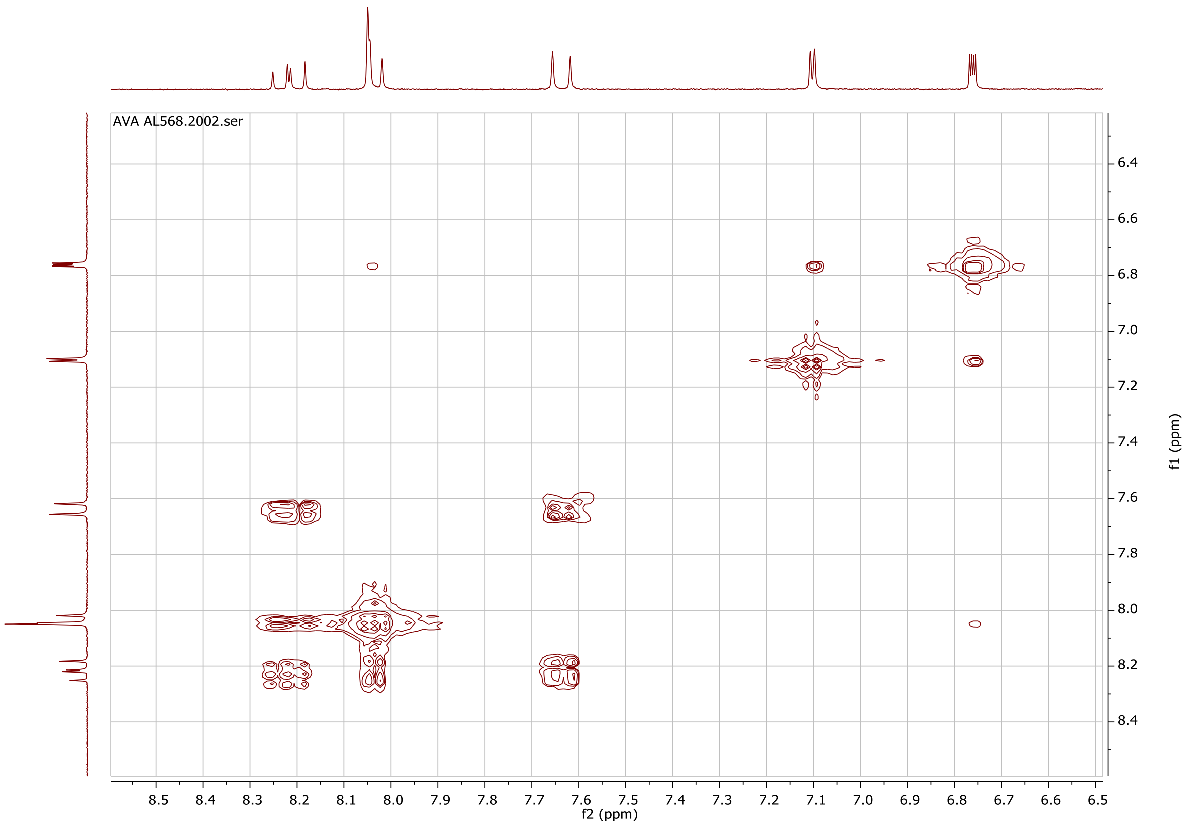
AVA AL568.2002.ser
f2 (ppm)
f1 (ppm)
